# Intrinsically Disordered Regions Promote Protein Refoldability and Facilitate Retrieval from Biomolecular Condensates

**DOI:** 10.1101/2023.06.25.546465

**Authors:** Philip To, Atharva M. Bhagwat, Haley E. Tarbox, Ayse Ecer, Hannah Wendorff, Zanya Jamieson, Tatjana Trcek, Stephen D. Fried

## Abstract

Many eukaryotic proteins contain intrinsically disordered regions (IDRs) that intersperse globular folded domains, in contrast with bacterial proteins which are typically highly globular^1, 2^. Recent years have seen great progress in identifying biological functions associated with these elusive protein sequence: in specific cases, they mediate liquid liquid phase separation^3^, perform molecular recognition^4^, or act as sensors to changes in the environment^5^. Nevertheless, only a small number of IDRs have annotated functions^6^ despite their presence in 64% of yeast proteins,^7^ stimulating some to question what ‘general purpose’ they may serve^8, 9^. Here, by interrogating the refoldability of two fungal proteomes (*Saccharomyces cerevisiae* and *Neurosporra crassa*), we show that IDRs render their host proteins more refoldable from the denatured state, allowing them to cohere more closely to Anfinsen’s thermodynamic hypothesis^10, 11^. The data provide an exceptionally clear picture of which biophysical and topological characteristics enable refoldability. Moreover, we find that almost all yeast proteins that partition into stress granules during heat shock are refoldable, a finding that holds for other condensates such as P-bodies and the nucleolus. Finally, we find that the Hsp104 unfoldase^12^ is the principal actor in mediating disassembly of heat stress granules and that the efficiency with which condensed proteins are returned to the soluble phase is also well explained by refoldability. Hence, these studies establish spontaneous refoldability as an adaptive trait that endows proteins with the capacity to reform their native soluble structures following their extraction from condensates. Altogether, our results provide an intuitive model for the function of IDRs in many multidomain proteins and clarifies their relationship to the phenomenon of biomolecular condensation.

## Main

Intrinsically disordered regions (IDRs) play well-understood roles such as presenting short linear motifs that are recognized by other proteins,^13^ sensing and responding to the environment, or folding upon binding to a partner; yet these accepted functions only account for a small fraction of the disorder in the *Saccharomyces cerevisiae* (hereafter referred as yeast) proteome, for which 64% of proteins are predicted to contain disordered regions of thirty residues or more and the median protein is 19% disordered. Increasingly, IDRs have been assigned an important role in promoting the formation of biomolecular condensates^14^; however, this relationship is also unclear, as some well-known phase-separating proteins contain little disorder whilst many disordered proteins are known to be highly soluble^15^. Hence, a general function for disordered regions has not been clearly established, though the identification of one could prove important for explaining their pervasive presence in eukaryotic proteomes.

Global refolding assays analyzed with structural proteomics have recently provided new insights into which types of proteins can locate their native conformations spontaneously^16^, or require assistance from chaperones^17^, or struggle to refold at all. So far, these studies have been limited to *E. coli* whose proteins are generally very ordered (the median protein is 3.5% disordered). Here, we present global refolding assays on two fungi, the single-celled *Saccharomyces cerevisiae* and filamentous *Neurospora crassa*. The data show that disordered regions promote the spontaneous refolding of their host proteins – enabling them to hew more closely to Anfinsen’s thermodynamic hypothesis – and suggesting an adaptive function in higher organisms for which proteins frequently need to transit between compartments, a process that sometimes involves unfolding them.

Heat stress granules are biomolecular condensates that rapidly form in response to elevated temperature, but which can also be efficiently dispersed in an ATP dependent manner^18^. We find that yeast proteins associated with this condensate evince unusually high levels of refoldability, and that during recovery they are processed by the Hsp104 disaggregase, with no detectable participation from degradative complexes. This observation provides a biophysical rationale for why proteins in heat stress granules can be efficiently retrieved without needing to get degraded, potentially explaining why yeast are so well-adapted to survive the commonly encountered stressor of elevated temperature. Moreover, we find that high refoldability is a common feature to proteins associated with other dynamic cellular compartments, including P-bodies and the nucleolus. Hence, these studies establish an adaptive role for reversible refoldability initially observed by Anfinsen and shed light on a potential general purpose for IDRs in fungi, and possibly all eukaryotes.

## Yeast Proteins are Surprisingly Refoldable

Drawing upon previously established methods^16, 17^, we carried out global refolding experiments on the soluble yeast proteome, and interrogated the conformational ensemble populated by yeast proteins via limited proteolysis mass spectrometry (LiP-MS^19^; Figure 1A). Briefly (see Experimental Methods), in these experiments we lyse yeast by cryogenic pulverization, unfold the extract by supplementing it with denaturant and reductant, and concentrating *in vacuo* (to 6 M guanidinium chloride and 10 mM DTT), and refold by 100-fold dilution. Pulse proteolysis via 1 minute incubation with proteinase K (PK) enables only the most solvent-accessible and unstructured portions of proteins to be cleaved by PK (Extended Data Figure 1A-C). Subsequent trypsinolysis allows sites where PK cleaved to be resolved by identifying half-tryptic peptides, sequenced and quantified with liquid chromatography tandem mass spectrometry (LC-MS/MS). The same procedure is carried out on a native extract that was never unfolded but is compositionally identical to the global refolding reactions (hence, these two samples differ only in history). By comparing the proteolysis profile of each protein in its native and refolded forms, we thereby assess whether a protein can refold spontaneously or not. Proteins are labeled nonrefoldable if we detect two or more sites with significant changes in proteolytic susceptibility (P < 0.01 by t-test with Welch’s correction) in the refolded form.

**Figure 1.**
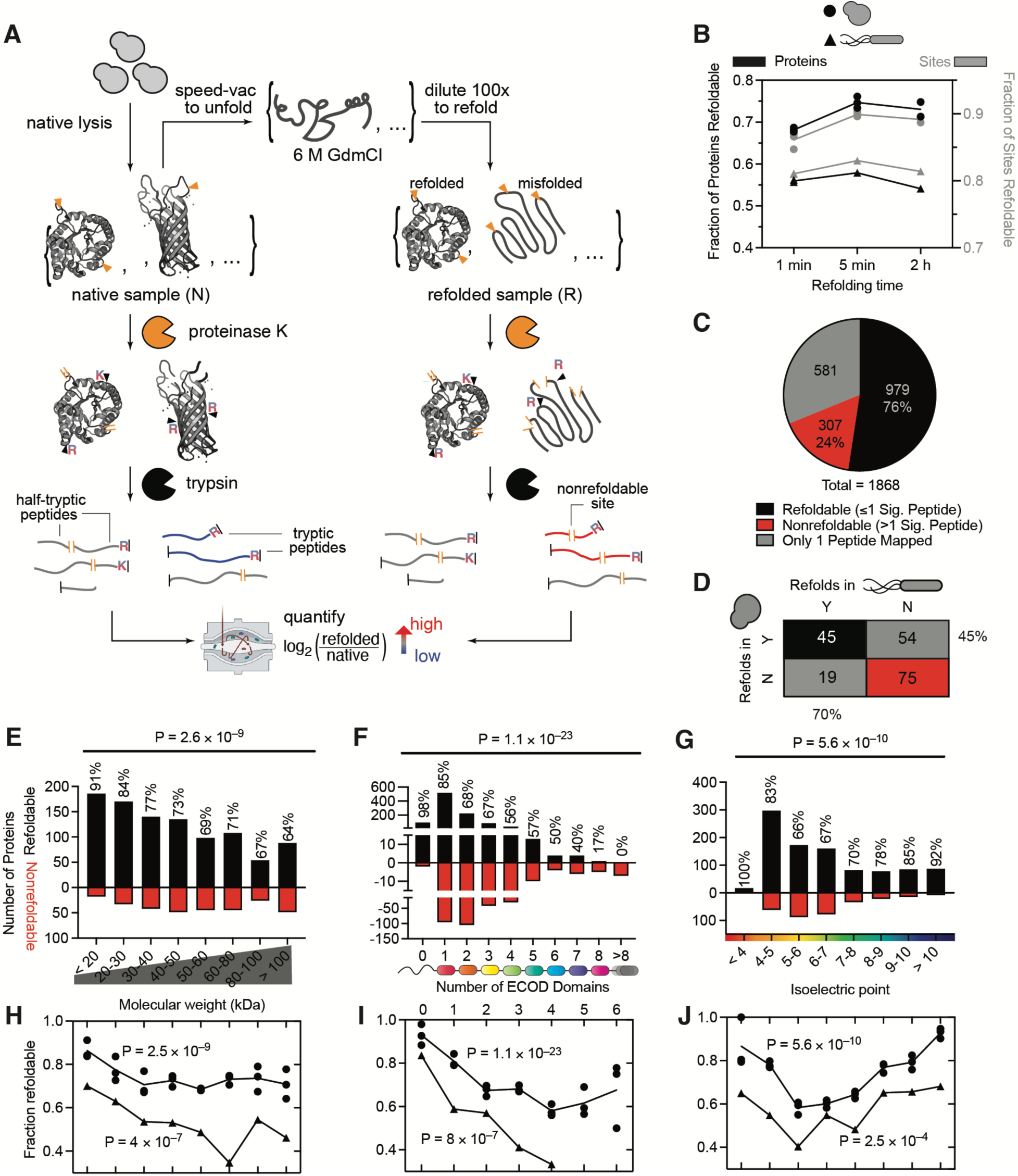
The Yeast Proteome Spontaneously Refolds More Efficiently than *E. coli*’s. (A) Schematic of experiment in which global refolding reactions on cell extracts are analyzed by limited-proteolysis mass spectrometry (LiP-MS). (B) Fraction of sites (i.e., mapped peptides; gray) and proteins (black) that are refoldable in yeast (circles) or *E. coli* (triangles), based on if their susceptibility to proteinase K is the same or different after unfolding/refolding compared to a native reference. Refolding reactions were analyzed at three times following dilution from denaturant. (C) Pie chart showing (non)refolding protein counts after 5 min. (D) Contingency table showing the number of *E. coli*/yeast orthologous pairs with a given refoldability status in either organism (5 min refolding time). (E-G) The number of refoldable (black) and nonrefoldable (red) proteins in yeast divided by (E) molecular weight, (F) number of folded domains, and (G) isoelectric point at the 5 min refolding time. (H-J) Fractional refoldability for proteins in the categories shown in E-G, for *E. coli* (triangles) and yeast (circles). Three separate performances of the assay were performed for yeast, each time with three biological replicates. P-values are from chi-square test; for yeast all P-values come from replicate 1.

After one minute of refolding, 68% ± 0.5% of yeast proteins had returned to their native state, and this increased to 75% ± 1% (977 out of 1286) after five minutes (Figure 1B-C, Extended Data 1D-G; error bars are std. devs from three separate experiments each of which was performed by refolding on three biological replicates). This overall refolding propensity remained unchanged after two hours (75%; 983 out of 1311). In similar experiments conducted on *E. coli* extracts^20^, the refolding frequency peaked at 5 minutes (58%; 649 out of 1121) and then slightly decreased after two hours. Hence, these studies revealed the surprising result that the yeast proteome is substantially *more* refoldable than the *E. coli* proteome despite being larger (average molecular weight of 53 kDa vs. 34 kDa), more multi-domain (28% multi-domain vs. 21%), and possessing aggregation-prone proteins at a 3-fold higher frequency than the *E. coli* proteome^21^. This ∼20% uptick in refoldability is also apparent among yeast proteins which possess *E. coli* orthologues (Figure 1D), and hence cannot be attributed purely to eukaryotic-specific proteins that lack bacterial counterparts. These assays were reproducible upon separate performances of the refolding experiment, each which involved refolding reactions on three biological replicates (Extended Data Figure 2); both at the level of quantifications for individual peptides (Extended Data Figure 2A-F), as well as at the level of protein refoldability calling (Extended Data Figure 2G-J).

Classifying yeast proteins by simple biophysical metrics such as molecular weight (MW, Figure 1E), number of domains (*N*Dom, Figure 1F), and isoelectric point (pI, Figure 1G) recapitulated correlations to refoldability that were previously identified in *E. coli*^16^; however, with stronger statistical significance. For instance, we found an inverse monotonic relationship between MW and *N*Dom with refoldability (P-value = 2.6×10^−9^ and 1.1×10^−23^, respectively by chi-square test). These results are intuitive to interpret from the perspective that *in vivo* proteins initially fold cotranslationally^22, 23^, during which vectorial synthesis provides a means to fold one domain at a time, decoupled from others, an advantage that is absent during refolding from a denatured state. As in *E. coli*, we find that mildly-acidic proteins (pI between 5–6) are the worst refolders whereas the most basic proteins (pI > 10) are the best (P-value = 5.6×10^−10^ by chi-square test). All these trends were highly significant and fully recapitulated at the level of individual PK cut-sites (Extended Data Figure 1H-J, P-value = 4.7×10^−29^, 1.4×10^−32^, and 1.3×10^−41^ respectively by chi-square test) implying that the findings are robust and not biased by differences in coverage or abundance amongst these categories. We also found that yeast proteins which do not engage CCT (chaperonin) or Ssb (Hsp70) co-translationally were typically refoldable (83% ± 2% and 88% ± 2%, respectively), whilst those which bind extensively (>4 sites) generally are not (35% ± 5%, 33% ± 7%), suggesting that many proteins which do not refold spontaneously in our assay require chaperone assistance (Extended Data Figure 3A-B)^24^. Overall, yeast proteins echo the same refoldability trends as the *E. coli* proteome (Figure 1H-J), but in any one of these categories, yeast proteins are ∼20% more refoldable than their *E. coli* equivalents (similar to the overall increase, Figure 1B). This suggests the existence of some global feature inherent to the yeast proteome that confers an improved capacity to refold from the denatured state.

## Intrinsic Disorder is a Key Predictor of Spontaneous Refoldability

When we classified yeast proteins based on their disorder content^25^, we obtained a striking correlation (Figure 2B; the way % Disorder is defined for each protein is shown in Figure 2A). Proteins that are fully disordered (% Disorder > 80%, called intrinsically disordered proteins, IDPs) are all (49 out of 49, 100%) “refoldable.” This result is – in some sense – expected, given that chemical denaturation should have a minor effect on IDPs, and thus serves as a positive control for our approach. On the other end, proteins that are fully folded/globular (Disorder < 5%) had the lowest refolding rate (53 out of 106, 50%), and as disorder content increases, refoldability increases in lockstep (Figure 2B, P-value = 5 × 10^−29^ by chi-square test). The trend was even more significant at the level of individual PK cut-sites, as assessed in two ways. Firstly, PK cut-sites mapped to regions that were assessed as disordered were significantly less likely to be altered by global unfolding/refolding (P-values by Fisher’s exact test shown with contingency tables in Figure 2F), as well as PK cut-sites associated with proteins with greater disorder content (P-value = 5 × 10^−68^ by chi-square test, Extended Data Figure 4A), disproving a potential bias arising from sequencing coverage or differences in PK’s cleavage activity on (dis)ordered regions. Hence, this correlation very neatly explains why the yeast proteome refolds more efficiently than *E. coli*’s: it is because it is more disordered (Figure 2C, E). Indeed, if *E. coli* proteins are subdivided by the same disorder categories, a very similar set of refolding propensities are obtained (Figure 2C). The difference is that the *E. coli* proteome has very few proteins with greater than 20% disorder, whereas yeast has many. We encountered a similar correlation when dividing proteins based on what percent of the sequence is part of an annotated domain (Extended Data Figures 3C, 4B) – the greater the proportion of the protein that is within domains, the lower its propensity to refold.

**Figure 2.**
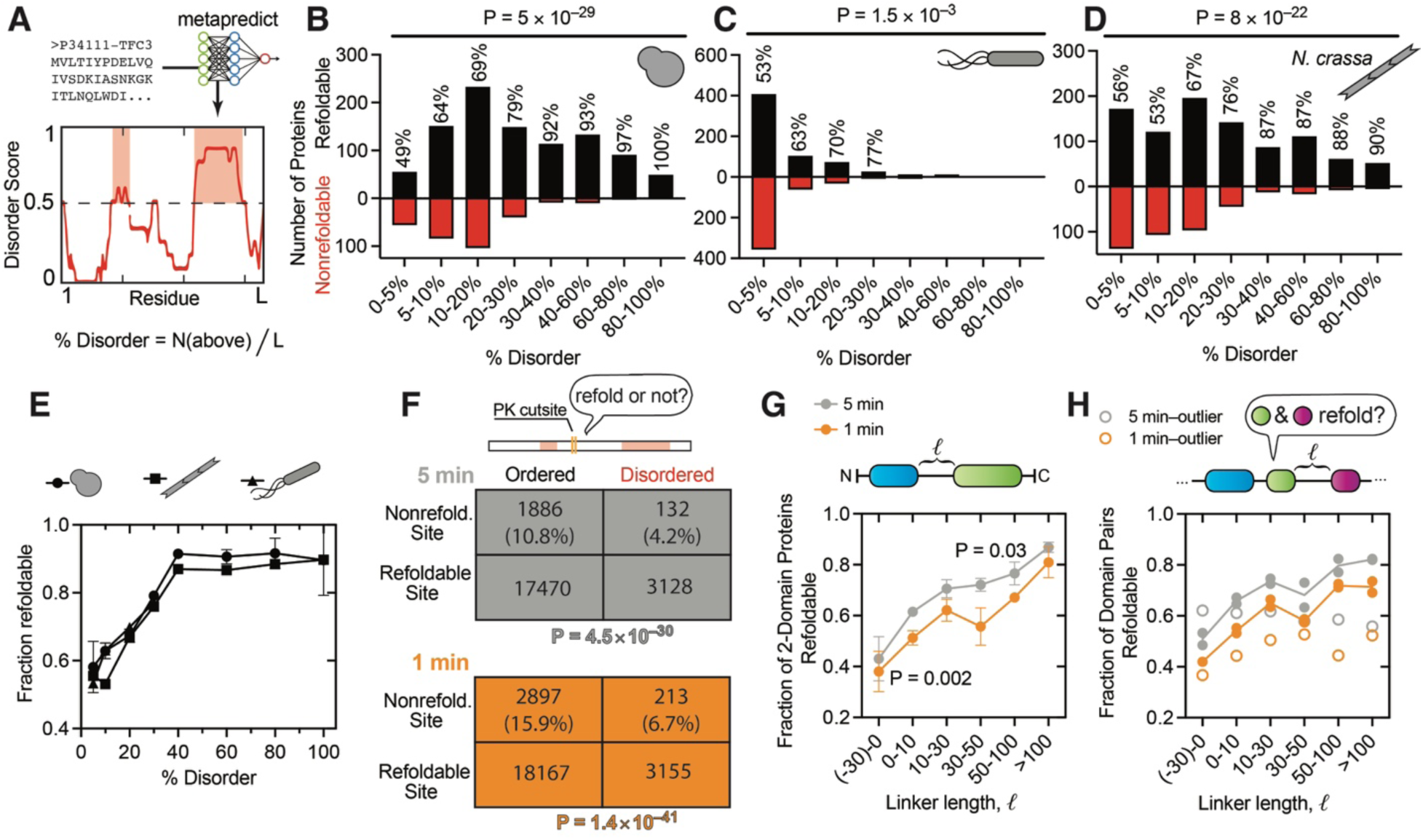
Intrinsic Disorder Underlies Refolding Propensities Within and Between Species. (A) Schematic diagram of how % disorder is predicted for each protein sequence, calculated with metapredict. (B-D) The number of refoldable (black) and nonrefoldable (red) proteins in yeast (B), *E. coli* (C), and *Neurospora crassa* (D) as a function of percent disorder in each protein at the 5 min refolding time. P-values are from chi-square test. (E) Fractional refoldability for proteins in the same % disorder categories for yeast (circles), *N. crassa* (squares), *E. coli* (triangles). (F) Contingency tables showing the number of assessed sites that are refoldable or not in terms of whether they fall in an ordered or disordered region. Sites in disordered regions are substantially (and according to Fisher’s exact test, significantly, P-values given) more likely to refold. Colour denotes refolding time (5 min, gray; 1 min, orange). (G) Fractional refoldability of two-domain proteins in yeast as a function of linker length between domains, calculated with DomainMapper. (H) Fractional refoldability of consecutive domain pairs within all yeast proteins with ≥2 domains as a function of linker length between pair of domains. Signal was not detected in one of three datasets (shown as empty circles).

**Figure 3.**
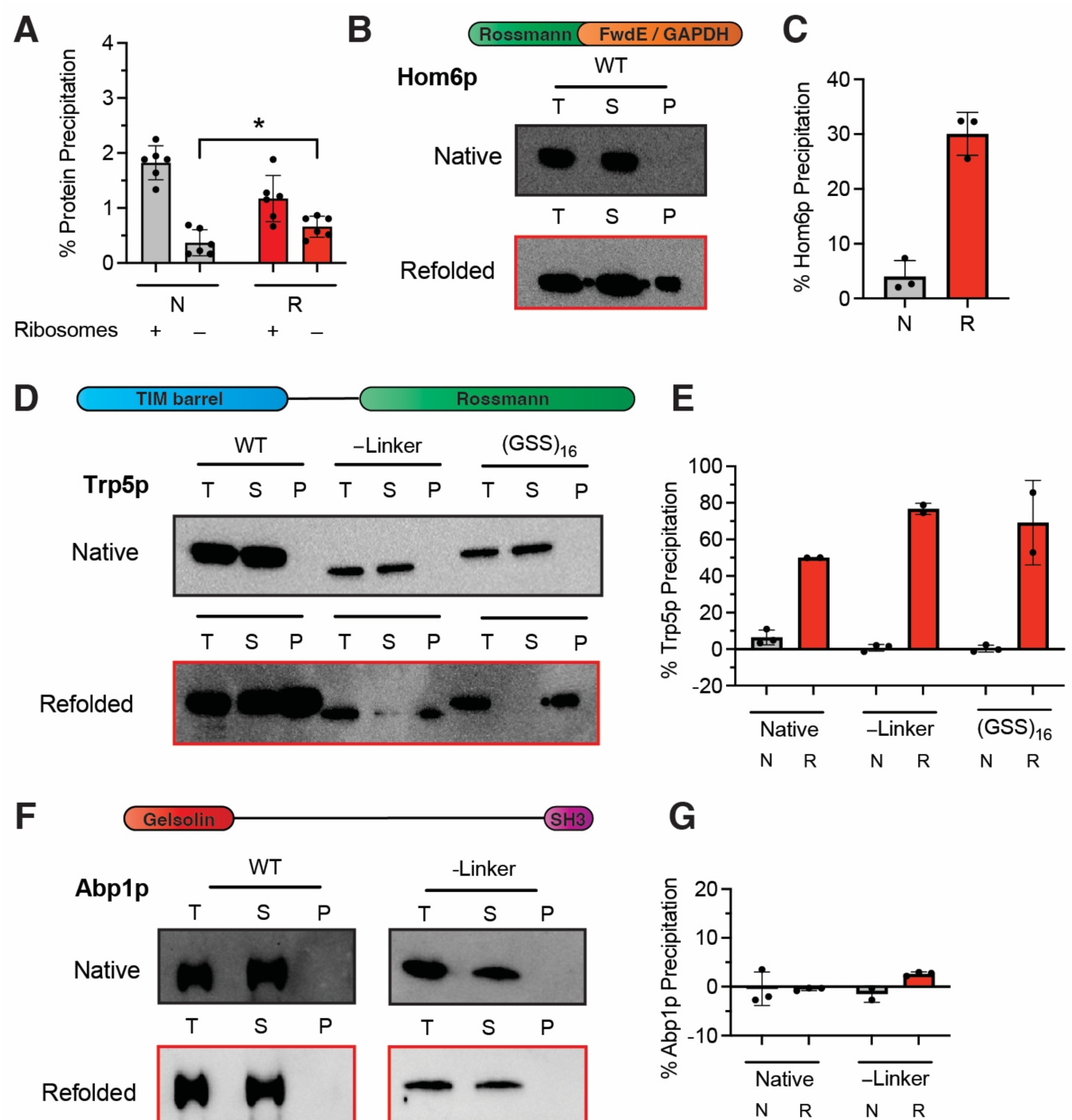
Aggregation Assays on Wild-type Yeast or Overexpression Strains. (A) Yeast extracts were assessed for precipitating aggregates (upon centrifugation at 16,000 *g* for 15 min) via the bicinchoninic acid (BCA) assay, either in their native form (N) or following a cycle of global unfolding/refolding (R). There was no significant level of aggregation in the refolding reactions. (B,C) A strain of yeast that overexpresses Hom6p (domain map given) was grown, and its extract retained either in a native form (N) or subject to global unfolding/refolding (R). (B) Aggregation of Hom6p was assessed by Western Blot of total extract (T), soluble portion (S), or pelleting portion (P). Representative image of three biological replicates (see Figure S8). (C) Densitometry analysis of western blots on biological triplicates (n = 3). (D, E) As for panels B & C, except for a strain of yeast that overexpresses Trp5p. Three variants of Trp5p were tested, wild-type (WT), a variant in which the linker between the two domains was removed (-Linker), and a variant in which the linker between the two domains was replaced with a flexible sequence of equal length, (GSS)16. (F, G) As for panels B & C, except for a strain of yeast that overexpresses Abp1p. Two variants of Abp1p were tested, WT and a variant in which the linker between the two domains was removed.

**Figure 4.**
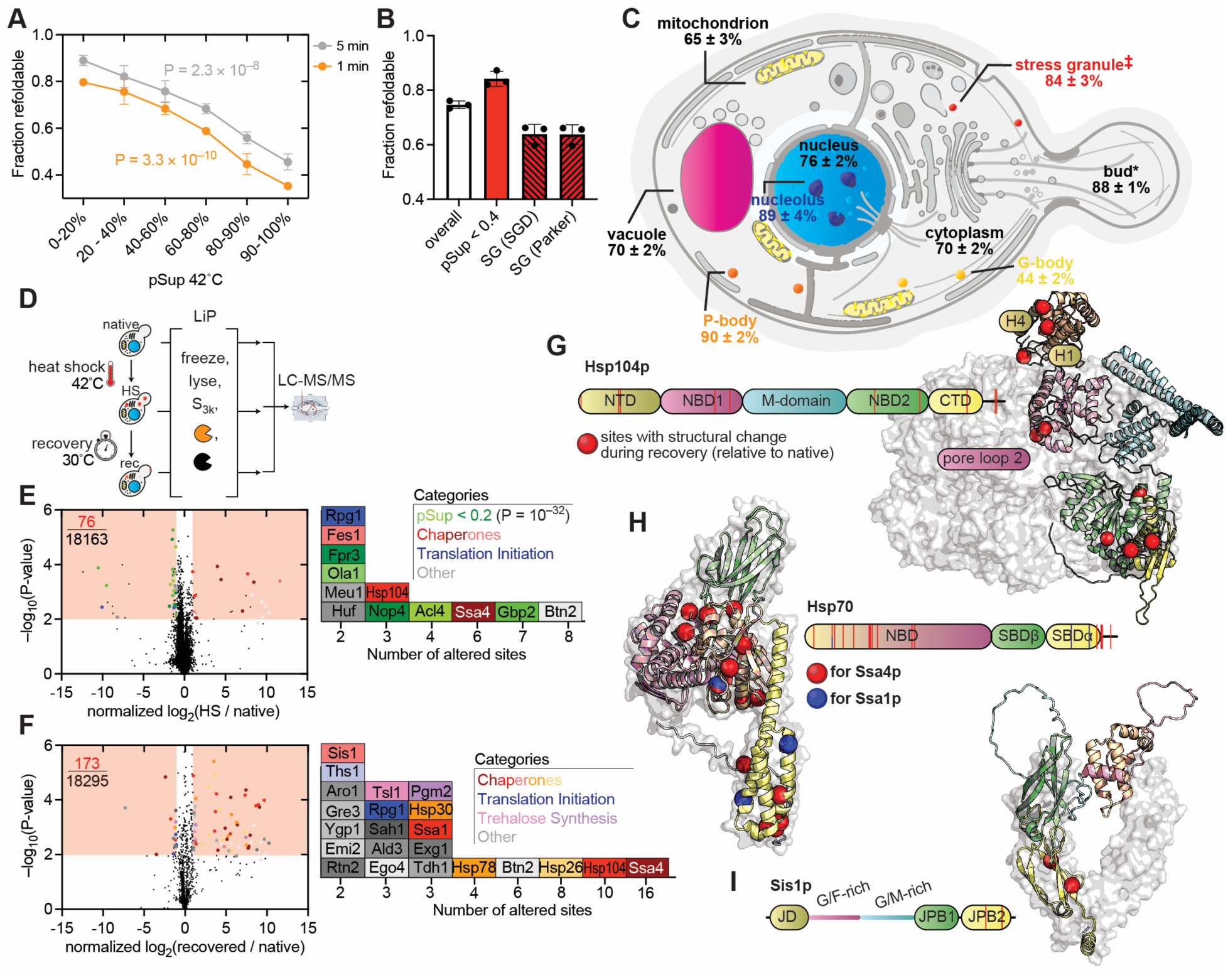
Heat Stress Forms Native-like Condensates Comprised Primarily of Refoldable Proteins. (A) Fraction of yeast proteins that refold after either 1 min (orange) or 5 min (gray) as a function of pSup42°C (proportion in the supernatant after heat shock at 42 °C), a measure of the protein’s tendency to partition to heat stress granules. P-values are from chi-square test from replicate 1 of the experiment. (B) Fraction of yeast stress granule proteins that are refoldable, depending on the definition used for stress granule resident proteins (pSup < 0.4, less than 40% of the protein is soluble after heat shock; SGD, according to the *Saccharomyces* Genome Database under Cellular Component; Parker, according to ref. 42 in which yeast were subject to oxidative stress with azide. (C) Fraction of yeast proteins that refold after 5 min as based on their cellular component (according to SGD), or according to ref. 46 in the case of G-bodies. (D) Schematic of experiment to use LiP-MS to assess proteins that undergo structural changes during heat shock (HS) or recovery. (E, F) Peptide volcano plots comparing extracts from cells subjected to (E) heat shock, or (F) subjected to heat shock and then allowed to recover. Regions in red represent cut-offs for significance; in upper-left corner is the fraction of total peptides quantified that report on significant structural changes. To the right are all proteins with significant structural changes, sorted by the number of significant peptides (columns) and functional category (colours). (G) Domain map and structure of Hsp104 (AlphaFold) imposed on cryoEM structure of hexamer (PDB: 5VJH). Red spheres represent sites with structural change during recovery from HS. (H) As panel G, except of Ssa4. Blue spheres represent Ssa1 sites with structural changes aligned onto Ssa4 positions. (I) As panel G, except of Sis1 imposed on x-ray structure of dimer (PDB: 2B26).

We wondered what role disordered regions could play in facilitating refolding. To consider a simple set of cases, we examined all two-domain proteins with a tail-to-head topology and asked what impact the spacing between two domains has on the refoldability of the protein^26, 27^ (Figure 2G). A clear trend emerged: the longer the spacing between two domains are, the more expeditious and efficient the refolding of the protein is. In the most extreme case where the two domains ‘feathered’ into each other, the refolding rate was very low (35% ± 4%) after both 1 min and 5 min, but this increased to 85% ± 1% for the subset of two-domain proteins with 100 amino acids of spacing or more (Figure 2G). The latter quantity is striking because it is identical to the overall refolding rate among single-domain proteins (Figure 1F). For linkers of intermediate length (10–50 residues), refolding was worse after 1 min, but improved after 5 min.

To generalize this concept to proteins with any number of domains, we analyzed pairs of consecutive domains in the context of multidomain proteins, and asked for each pair is there a relationship between the linker that intervenes between the two domains and their refoldability statuses. Though the trend was weaker, it qualitatively reproduced the same correlation amongst two-domain proteins (Figure 2H).

As for the linker regions themselves, we found that PK cut-sites that were far from any folded domain (>100 residues) *or* that were located in a long linker region (>250 residues) were close to 100% refoldable, whereas PK cut-sites in linkers in proximity to domains followed trends closer to the folded regions themselves (Extended Data 4D-E). Interestingly, this trend only held for inter-domain linkers (that is, those that are sandwiched between two domains), but not for “leading” or “trailing” non-globular regions. Such an observation supports the hypothesis that IDRs with independent functional roles are more likely to be found on the N- or C-termini, but that the primary function of inter-domain linkers is to act like a spacer or lubricant between their flanking domains, thereby improving the ability of complex multi-domain proteins to refold^2^. Previous works have suggested that non-native interdomain contacts constitute the primary way that multi-domain proteins misfold^28, 29^; our proteome-wide refolding data support this view and suggest that interdomain linkers facilitate refolding by lowering their chance of forming.

Since we were not aware of prior studies documenting this strong linkage between disordered regions and protein refoldability, we wondered whether this effect might be specific to *S. cerevisiae* (or budding yeast). Hence, we performed analogous global refolding experiments on the distantly-related fungus, *Neurospora crassa*^30^. Like yeast, *N. crassa* proteins were also highly refoldable (69% after 5 min of refolding; 75% after 2 h), and satisfyingly, all the trends between refoldability and intrinsic disorder were recapitulated (Figure 2D-E, Extended Data Figure 5A-B). All the basic biophysical trends governing refoldability were nearly identical in yeast and *N. crassa* (Extended Data Figure 5D-I), as well as the connection between refoldability of multidomain proteins and linker lengths (Extended Data Figure S6A-B). Refoldability status was 71% conserved between orthologous pairs (Extended Data Figure 5C). But quite interestingly, one feature that we found to be highly conserved is linker lengths interspersing domains between these two species (*R*^2^ = 0.70 for two-domain proteins; *R*^2^ = 0.78 for multidomain proteins) suggesting that this property is retained by evolution (Extended Data Figure 6E-F). Moreover, for the small subset of proteins whose yeast orthologue could refold but whose *N. crassa* orthologue could not, the yeast orthologue had greater disorder content (Extended Data Figure 6D).

## Nonrefolding Proteins Aggregate when Overexpressed

In global refolding assays on whole yeast extracts, we could not detect any precipitation above background (Figure 3A, 0.29 ± 0.30%). Hence, we reasoned that the yeast extract could be used as a ‘blank slate’ for aggregation studies of selected yeast proteins, a traditional biochemical approach for measuring refoldability or lack thereof^31, 32^. We cloned Hom6 (homoserine dehydrogenase, involved in biosynthesis of Met^33^) driven by a GAL1,10 promoter onto a plasmid with a 2µ origin for over expression. Hom6 possesses several properties representative of yeast nonrefolders: it is a metabolic enzyme with two partially overlapping domains (i.e., a negative linker length), one of which is a Rossmann fold (a fold-type which is a relatively poor intrinsic refolder, 65% vs 83% overall, see Extended Data Figure 7 for refoldability levels of common fold-types). Cells were induced, grown to the end of log-phase, lysed, and then their extracts globally unfolded and refolded as previously described. Over expression of Hom6p resulted in extracts that were much more aggregation-prone during global refolding (32.2 ± 9%, Extended Data Figure 8A). Moreover, Western blots specific for Hom6p confirmed that one-third of the protein (30.1 ± 4%) precipitated after being diluted from denaturant (Figure 3B-C). These results suggest that at endogenous levels, Hom6p populates soluble misfolded conformations (which LiP-MS differentiates from native conformations) that do not aggregate but also cannot self-correct (at least on the timescale of the experiment, upto 2 h); at the higher concentrations from over expression however, Hom6p aggregates instead.

A similar set of results were obtained when Trp5p (tryptophan synthase) was over-expressed. Like Hom6, Trp5 is also a two-domain protein involving a Rossmann fold, but that has a longer structured linker that connects it to a TIM barrel (one of the poorest spontaneous refolders^18, 34^). Upon over-expression, Trp5p’s inherent nonrefoldability also results in aggregation upon attempted refolding: by Western blot, 50.0 ± 0.7% of Trp5p precipitates, as well as 15.1 ± 9% of all protein in the lysate (Figures 3D-E, Extended Data Figure 8D). Moreover, when Trp5p’s structured linker is removed the phenotype is exacerbated and virtually all of Trp5p precipitates, consistent with the general trends from proteome-wide refolding (Figure 2).

To find supporting evidence that refoldable proteins do *not* precipitate upon refolding, even when present at high concentrations from over-expression, we focused on Abp1 (actin-binding protein). Like Trp5, Abp1 is a two-domain protein, but similar to other refolders, it possesses a long interdomain linker. Moreover, one of its domains is an SH3, which has high refolding propensity (96% ± 5%, Extended Data Figure 7)^35, 36^. Yeast was less tolerant to Abp1p over-expression: cells grew slowly after induction and Western blot revealed a range of degradation products (see Extended Data Figure 8 for all raw Western blots). Nevertheless, Abp1p and even all its partially degraded forms remained fully soluble after dilution from denaturant (Figure 3F-G, Extended Data 8J).

These experiments enable us to connect the formation of soluble misfolded conformations (probed in LiP-MS assays) with the more prototypical measure of nonrefoldability – aggregation. Each individual protein in diluted lysates has a concentration that is too low to aggregate^37, 38^, but at elevated concentrations, the soluble misfolded conformations can aggregate.

Having established that two fungal proteomes are generally adept at spontaneously refolding following chemical denaturation, that this property is linked to their higher levels of disorder, and that disordered regions have been retained in fungal multidomain proteins, we were stimulated to wonder whether there is any *biological* reason for proteins to be refoldable. This question is not obvious given that: (i) proteins generally fold cotranslationally on the ribosome during primary biogenesis; and (ii) 6 M guanidinium is not a physiologically relevant stressor. To answer this question, we next considered how yeast responds to a frequently encountered stress, namely heat shock, and whether that is connected to refoldability.

## Proteins Associated with Most Biomolecular Condensates are Highly Refoldable

During heat shock, many yeast proteins form transient foci that are visible by fluorescence microscopy^18, 39^. In a study by Wallace and co-workers,^40^ biochemical fractionation was performed on transiently heat-stressed yeast (42°C for 8 min) and LC-MS/MS used to quantify what percent of each protein remains soluble (i.e., in the supernatant, pSup). When we cross-correlated pSup values for 747 yeast proteins for which refoldability assessments were also available, we obtained the result that yeast proteins which completely condense upon heat shock (pSup≤20%) can generally refold (89% ± 2%), whereas those which stay soluble (pSup≥90%) generally cannot (45% ± 3%; Figure 4A). The effects are more pronounced when considering heat shock at higher temperature (46°C) or the shorter refolding time (1 min), suggesting that the more a protein condenses, the more it has also been selected to refold quickly (Extended Data Figure 9A). The same trends are found when refoldability is correlated with pSup’s filtered for baseline levels of protein sedimentation (e.g., ΔpSup 30°C→42°(Extended Data Figure 9B-C)) and are robustly recapitulated at the level of individual PK cut-sites, disproving any artefact from coverage bias (Extended Data 9D-E).

A high association between refoldability and stress granule localization is detected when we use a dataset based on *heat* stress^40, 41^ (Figure 4B). The trend is not apparent when the definition is applied to proteins that condense in response to oxidative stress^42^, or when used more expansively to refer to particles formed in response to various stressors^43, 44^. This is intriguing, because heat is a physiologically salient stressor yeast experience regularly and is also one from which they can recover rapidly^40, 45^. Strikingly, we find high refoldability is associated with other *dynamic* biomolecular condensates (Figure 4C) – that is, those which can disperse rapidly – such as P-bodies (90 ± 2%) and the nucleolus (89 ± 4%). As a counterexample, G-bodies are a condensate which yeast form in response to severe hypoxia and which persist for long periods of time (hours or days) after normoxia is restored^46, 47^. Consistently, their proteins (as well as glycolytic proteins in general^16, 17^) are mostly nonrefoldable (44 ± 2%, Figure 4C). Moreover, one of the more prominent resident proteins in G-bodies is Cdc48p (which acts upstream of the proteasome), consistent with G-body clearance occurring by way of degradation. In contrast, proteins in heat stress granules (HSGs) have been shown to escape degradation following thermal stress, instead getting restored to the cytoplasm^40^. These observations lead to the question of which biophysical properties consign a protein in a condensate/aggregate to get degraded or enable it to be restored/refolded when the aggregate is dispersed.

## Heat Stress Granules Are Processed by Hsp104, not by Degradative Machines

To better understand the molecular players involved during the recovery from heat stress and how it might relate to refoldability, we performed a separate set of LiP-MS experiments in which yeast were subject to heat shock at 42°C (for 8 min), and either immediately frozen and lysed by cryogenic pulverization or first allowed to recover at 30°C for 1 hr before freezing (Figure 4D). Pulse proteolysis of PK was conducted in the lysate under conditions that maintain HSGs. Hence, these limited proteolysis (LiP) profiles identify which proteins are structurally altered after HSGs have formed or during their disassembly. In marked contrast to our refolding experiments, protein conformations are virtually unchanged by heat stress (Figure 4E), with only 12 (out of 1532, <1%) proteins structurally altered (using the same criteria and cut-offs). This finding is broadly consistent with one previous report that suggested proteins in HSGs are not misfolded and can sometimes still execute their native functions^40^. The set of 12 contains three chaperones (Hsp104p, Ssa4p, Fes1p), a translation initiation factor (Rpg1p/eIF3a)^48^, and a set of five proteins with very low pSup values (Ola1p, Fpr3p, Nop4p, Acl4p, Gbp2p)^40^ previously termed “super-aggregators”. The last group highly overrepresents the total observed structural changes (20/76=26%, despite representing 5/1532=0.3% of proteins; Extended Data Figure 9F-G). Together with the observations that the super-aggregators form foci very rapidly^40^ and disperse slowly (shown in the following), we hypothesize that these five proteins in their altered conformations formed during heat shock serve as scaffolds^42, 49^ for HSGs. Moreover, Nop4 and Gbp2 consist of many RNA-binding domains which suggests they could produce a multivalent network with RNA. The overall abundance of scaffold proteins is not increased by heat shock (Extended Data Figure 9J), consistent with the view that their condensation is protein-autonomous, not mediated through gene regulation.

During recovery, slightly more structural differences are present (22 altered proteins out of 1526; 173 sites with significant changes in PK susceptibility out of 18,295, Figures 4F). The “scaffold” proteins have completely returned to native-like conformations, but now the structural differences are concentrated on two small heat shock proteins and two chaperones associated with HSGs disassembly: Hsp104p (a hexameric AAA+ ATPase) and Ssa4p (an Hsp70). These two proteins account for 15% (26/173) of all detected structural changes. We also find structural changes in Ssa1p (the housekeeping paralogue of Ssa4), Hsp78 (the mitochondrial equivalent to Hsp104^50^), and the trehalose synthesis pathway (Figure 4F). On Hsp104p (Figure 4G), we identified three altered sites on the N-terminal domain (NTD) on helices 1 and 4, consistent with this region’s recently assigned role in loading substrates and interacting with Hsp70s^51, 52^ as well as two altered sites on the second pore loop of the first ATPase domain (called a nucleotide-binding domain, NBD) involved in substrate translocation^53^. All the structural perturbations faced into the pore based on cryo-EM models of the hexamer^54^. Ssa4p is completely remodeled (Figure 4H), particularly near the ATP binding pocket and the alpha-helical portion of the substrate binding domain (SBD), oftentimes considered as the “lid” that clamps down on bound clients^55^. Ssa1p is modified to a lesser extent, albeit in the same regions as Ssa4p (Figure 4H, blue spheres). Sis1p (but not its paralogue, Ydj1p), a J-domain protein, is also structurally altered in its second J-domain peptide binding domain (JPB, Figure 4I), consistent with its recently-clarified role as the key J-domain protein in disaggregation^56, 57^.

Cdc48p and the proteasome were *not* detectably structurally altered by heat shock or during recovery despite high coverage (52 and 251 peptides quantified respectively) nor even upregulated, suggesting they are not activated over baseline levels. Neither yeast’s chaperonins (CCT, Hsp60) nor Hsp90 were structurally altered (Extended Data Figure 9K). Taken together, these studies point to Hsp104 as the primary chaperone responsible for disassembling HSGs *in vivo*,^58^ consistent with its demonstrated efficacy to disperse condensates of pure Pab1p (an abundant protein in HSGs) *in vitro*^57^.

There is an intuitive appeal for HSG proteins to be refoldable if they are destined to become clients of Hsp104p. Because Hsp104p acts as a processive unfoldase^12^, refoldability could serve an adaptive function because it would enable HSG-proteins to reactivate spontaneously after being processed once by Hsp104 (Figure 5A). Hence, the focused outcome of these structural proteomics experiments (cf. Extended Data Figure 9F-I) reinforces the view that HSGs can be dispersed without their resident proteins getting degraded and identify which chaperones are active in this process^40^.

## Refoldability Facilitates Retrieval of Proteins from Heat Stress Granules

What properties make a protein an amenable client for Hsp104 under native physiological conditions? To address this question, we generated a yeast strain in which Pab1 (an HSG marker protein) was doubly labeled with a fluorescent protein (mClover3) and a FLAG tag at its chromosomal locus. The cells were subjected to heat stress at 42°C or 46°C, recovered at 30°C for various time intervals, lysed under conditions that maintain HSGs, and then HSGs were purified by differential centrifugation and anti-FLAG immunoprecipitation (Figure 5B). Isolated stress granules were then interrogated with bulk spectroscopy, fluorescence microscopy, and all their resident proteins identified and quantified with LC-MS/MS. Fluorescence micrographs of the isolated HSGs confirmed that their dimensions were consistent with their reported size *in vivo* (median radius 400 nm), and their numbers declined as recovery proceeded, with most vanishing after 30 min for the 42°C heat shock, but for the 46°C heat shock, their numbers do not drop precipitously until 1 h, as expected (Extended Data Figure 10).

**Figure 5.**
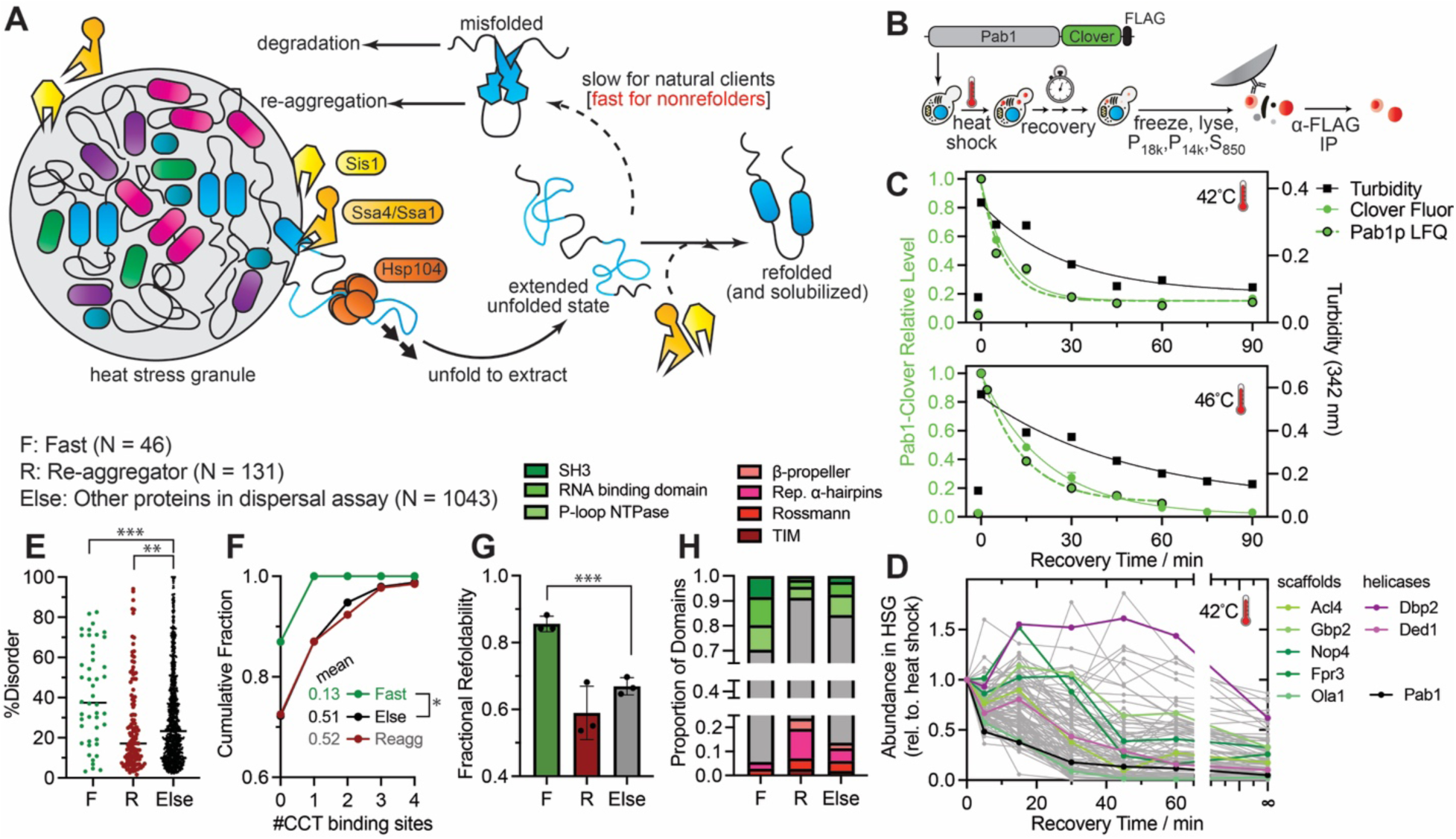
Refoldability is Associated with Efficient Retrieval from Heat Stress Granules during Recovery. (A) Model for heat stress granule (HSG) disassembly by Hsp104 and the importance of refoldability to complete retrieval to cytoplasm. (B) Schematic of experiment to isolate fluorescent affinity-tagged HSGs after stress and during recovery, whose time evolution are monitored with turbidity, fluorimetry, and quantitative mass spectrometry by label free quantification (LFQ). (C) Relative turbidity (black squares), Pab1-Clover fluorescence (green circles), and Pab1 abundance by LFQ (green circles with black borders) of the HSG fraction as a function of recovery time after 42°C (top) and 46°C (bottom) heat shocks (15 min). (D) Dispersal curves for all HSG proteins (pSup < 0.4) during recovery (30°C) from heat stress at 42°C. Scaffold proteins highlighted in shades of green; RNA helicases in shades of violet; Pab1p in black. “Infinite” time refers to control sample not subject to heat shock. (E-H) Properties associated with “fast” (F) proteins that are rapidly evicted from HSGs, “reaggregating” (R) proteins that experience large returns to the HSG fraction, and all other proteins assessed in the dispersal assay (else). (E) %Disorder for proteins in the three dispersal categories. ***, P<0.001; **, P<0.01 by Mann-Whitney test. (F) Number of CCT binding sites for proteins in the three dispersal categories, shown as cumulative density functions. *, P=0.015 by Mann-Whitney test. Average number of CCT sites per protein for each category given. (G) Fractional refoldability for each dispersal category (dots correspond to three separate performances of refolding experiment). ***, P<0.001 by two-tailed t-test. (H) Proportion of domains for proteins in each dispersal category belonging to each X-group.

Fluorimetry reveals that Pab1p is efficiently evicted from HSGs with single exponential kinetics (half-life 6.1 ± 1 min and 15.0 ± 1 min after heat stresses at 42°C and 46°C; Figure 5C), a rate in good agreement with *in vitro* Hsp104 dispersal assays (8.5 min) further supporting Hsp104 as the chaperone responsible for driving eviction *in vivo*. Moreover, the fluorescence-based dispersal rates are indistinguishable from those determined for Pab1p by quantitative mass spectrometry, implying the dispersal dynamics from our high-throughput assay are reliable (for which 1231 proteins’ dispersal curves were obtained). Pab1p’s dispersal is faster than the disappearance of HSGs as a whole (as determined by UV-Vis turbidity (half-life 19 ± 6 min and 34 ± 10 min, Figure 5C)) and is faster than the majority of individual HSG-proteins (as determined by LC-MS/MS (Figures 5D, Extended Data Figure 12A-D)). On the other hand, the preponderance of HSG proteins is solubilized faster than Hsp104’s activity on the “model” aggregation-prone protein luciferase (for which 1% is restored after 2 h)^57, 59^. These observations therefore beg the question as to what makes HSG proteins “amenable” clients for Hsp104 (Figure 5A).

To begin dissecting this, a qualitative comparison of Pab1 and luciferase is instructive. Pab1 consists primarily of RNA-binding domains (a fold with high inherent refoldability, 87% ± 1%) separated by medium-sized linkers (36 residues on average), and is 25% disordered. Luciferase, on the other hand, has 4 Rossmann-like domains that feather into each other (nonrefoldable traits) and is 1% disordered. In general, fold types and their spacings are independent predictors for protein refoldability (Extended Data Figures 7, 11A-B), as is isoelectric point, percent disorder, and number of domains (Extended Data Figure 11C-D). Moreover, we find that refoldable fold-types are enriched in all of yeast’s dynamic condensates (e.g., jelly-rolls are 3.9-fold enriched in HSGs, SH3-folds are 6.8-fold enriched in P-bodies, and OB-folds are 2.8-fold enriched in the nucleolus) whereas nonrefoldable fold-types are depleted from them (Extended Data Figure 11E).

To see if these ideas could generalize to our proteome-wide dispersal assay, we delineated two archetypal categories (Extended Data Figure 12E-F). We defined “fast” proteins as those which, after 5 min of recovery, have been more than 50% removed from the immunoprecipitated HSGs, and then continue to decrease thereafter until an hour of recovery. “Re-aggregators” are those which between any two consecutive timepoints during the hour of recovery, experience a large (>2-fold) *increase* in their abundance in the HSG fraction. The 46 fast proteins display a noticeably higher level of intrinsic disorder (41% on average), are very rarely clients for CCT during their primary biogenesis, and exhibit high levels of refoldability from denaturant (86% ± 2%) – even with only 1 min refolding time (89% ± 3%) (Figure 5E-G, Extended Data Figure 12G).

Two alternative (but not mutually exclusive) explanations for the relationship between intrinsic disorder and retrievability from HSGs are: (i) disordered regions are more readily loaded into Hsp104; and (ii) disordered regions facilitate refolding after translocation through. In support of the idea that refoldability is specifically important for efficient dispersal, we find refoldable domains (such as SH3 folds and RNA-binding domains) are also over-represented amongst rapidly-retrieved proteins (Figure 5H). Hence, in general, proteins which are most expeditiously retrieved from HSGs share features in common with Pab1. The 131 re-aggregators have the opposite attributes: they have less disorder, greater CCT dependency, fewer SH3 folds and RNA-binding domains, and lower levels of refoldability (Figure 5E-H).

Non-monotonicity in HSG disassembly during recovery is widespread, supporting the model in Figure 5A that some proteins re-aggregate following an unfolding cycle by Hsp104 (or alternatively, are recruited to HSGs during the recovery phase). In some cases, this behaviour might be functional. For instance, the stress granule-associated RNA helicases Dbp2p and Ded1p are replenished to HSGs between 5 and 15 min of recovery (Figure 5D), possibly because it becomes more important to actively unwind RNA to maintain HSG fluidity during the recovery process (since the elevated temperatures which stimulated HSG formation also inherently melts base-pairing). On the other hand, the majority of translation initiation factors (12 out of 14, excluding eIF1 and eIF1A, Extended Data Figure 12I) are evicted from stress granules relatively quickly (more than 80% dispersed by 30 min), as are the constituents of the ribosome associated complex (RAC) and multi-synthetase complex (Extended Data Figure 12J-K), consistent with rapid resumption of translation during recovery. Stress granule scaffold proteins all disperse more slowly (with the exception of Ola1), consistent with their rapid initial condensation and putative localization in the core of the condensate (cf. Figure 5D). Dispersal curves following heat shock at 46°C recapitulate all of these findings except are on the whole somewhat slower (Extended Data Figure S12H,J,K). Hence, proteome-wide dispersal assays show that spontaneous refoldability governs, in part, the compositional trajectory of HSGs during recovery, with proteins that refold well getting evicted more rapidly.

## Discussion

What is the relationship between intrinsically disordered regions (IDRs) and biomolecular condensates? IDRs are commonly invoked as playing the defining role in mediating condensate formation, a theory supported by defining work on several disordered RNA-binding proteins (e.g., FUS^60^ and hnRNPA1^3^) in human diseases that are now well-established model systems. On the other hand, folded domains provide the key scaffolding functions in other condensates, such as in multi-synthetase complexes^40, 41, 61^, *C. elegans* germ-granules^15, 62^, and pyrenoids from algae^63^. An emerging view holds that condensation can be mediated through electrostatic complementarity or multimerizing domains without any requirement for IDRs^64^. Why do certain condensate-resident proteins possess high levels of intrinsic disorder whilst others do not? Our proteome-wide assessments of spontaneous refoldability, mapping of structural changes associated with heat shock, and HSG-wide dispersal assays provide some insight into this question. Intrinsically disordered regions between folded domains promote refoldability by insulating domains from each other, enabling multi domain proteins to recapitulate the capacity of single-domain proteins to reversibly refold as demonstrated by Anfinsen’s foundational studies. Because processive unfoldases are employed to extract proteins from condensates^42, 57, 58^, refoldability is a useful trait for proteins that partition into dynamic condensates which must disperse rapidly in response to a signal or change in the environment. In other words, disorder is equally important for getting proteins *out* of condensates as it is to get proteins *into* them.

The existence of physiological processes that actively *unfold* certain proteins could produce an evolutionary pressure for such proteins to *refold* in a translation independent and energetically-inexpensive manner. The optimal outcome of such evolutionary selection would be, of course, spontaneous refoldability. Given that both processive unfoldases and 6 M guanidinium generate extended conformations^65^, it is possible that the yeast proteome’s high refoldability from denaturant is a byproduct of the preeminent role yeast’s proteostasis network assigned to Hsp104. On the other hand, most proteins fail to reversibly refold following thermal denaturation. But as has been pointed out^40, 57^, temperatures that denature mesophilic proteins (ca. 60–70°C) are typically much higher than any temperature the organism is adapted for (even during acute heat stress), so there would not have been evolutionary pressure for proteins to refold from such extreme thermal stress. Refolding from denaturant, on the other hand, may phenocopy the natural process of refolding following translocation through AAA+ unfoldase machines.

Lastly, these proteome-wide measurements can put biological context to several outstanding debates surrounding the mechanism of Hsp104-like unfoldases (Figure 6), for which there is not consensus about substrate topology, translocation rate, or processivity. Structural studies have so far only provided evidence for single-pass substrate threading^54^, though recent single-molecule experiments point to a processive loop-extrusion model^66^. Most early work took the view that Hsp104 fully translocates its clients^12, 67^, though studies since have provided evidence for partial threading^57, 68^. A theory is frequently discussed as to whether the vectorial appearance of substrate on the *trans* side of Hsp104 could recapitulate cotranslational folding (cf. Figure 6), which is worth reflecting on. Finally, various experiments have placed very different values to the translocation rate (between 5-500 residues/s), with the upper end supported by single-molecule studies^66^.

**Figure 6.**
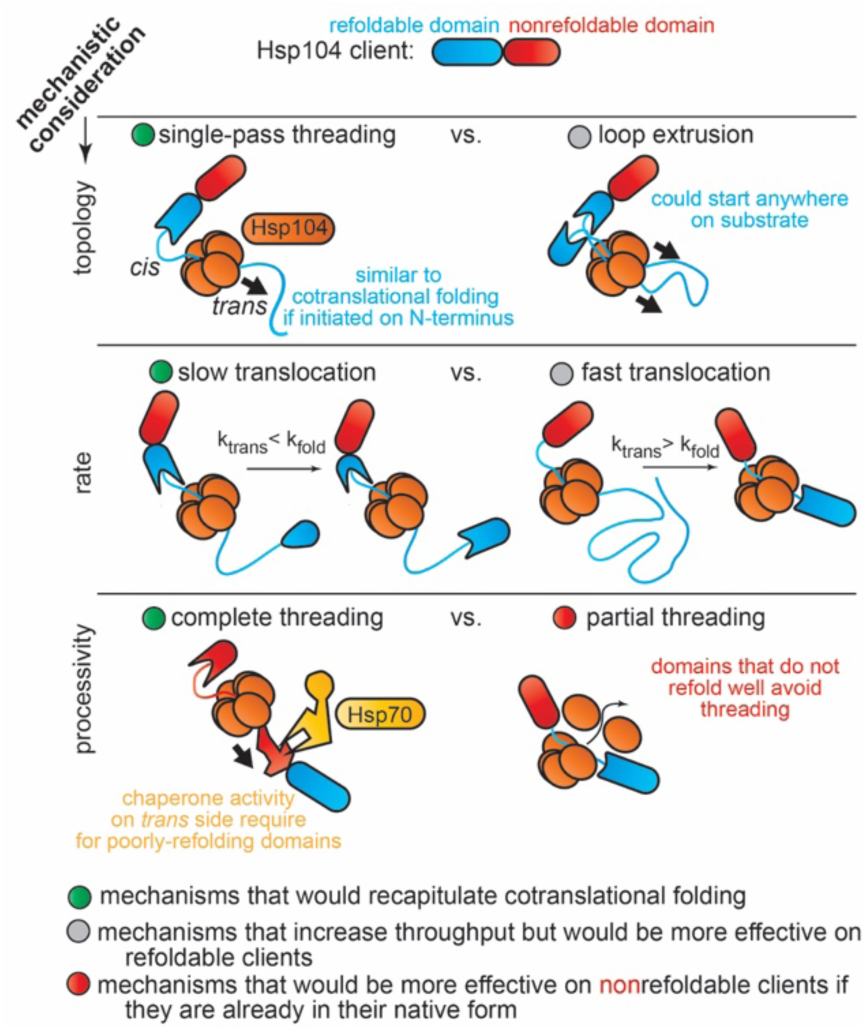
Comparison of Hsp104 Unfoldase Mechanisms. Top row, two models for the topology of threading. Middle row, two models for the translocation rate and how depending on its speed relative to folding rates Hsp104 would either recapitulate cotranslational folding or spontaneous refolding from denaturant. Bottom row, two models for processivity and their implications for a client that does not refold efficiently: further chaperone intervention would be required if such a portion were completely threaded.

Though our assays are not explicitly mechanistic, they can describe which mechanisms would be operative or useful in the physiological context of HSG dispersal. For instance, our data support the newer view that Hsp70s act *upstream* of Hsp104 to load clients into it^69^. Earlier models tended to assign Hsp70s a role in refolding clients after translocation through Hsp104^67^; however, the observation that most of the chaperone’s natural clients can refold unassisted implies this function would not generally be needed. Our data also support the view that Hsp104 can act as a rapid loop-extruder. Rapid translocation would generate extended unfolded conformations, resembling the conformations populated when proteins are unfolded with denaturant, and from which most HSG proteins can refold. In the loop extrusion model, translocation could commence anywhere in the protein and appearance of the unfolded chain is not vectorial. This is consistent with amenable clients being fully refoldable because the client could not preordain the order in which its segments get unfolded or reappear on the *trans* face.

A more subtle question is how to reconcile fast processivity with conflicting data that support partial threading or a non-processive mechanism. Though our assays do not directly observe partial threading, there are cases where it would be useful, because even though the majority of HSG proteins refold spontaneously, some cannot (cf. Figure 4A). Given that most HSG resident proteins are in native-like conformations (cf. Figure 4E) it would then be advantageous for native-but-nonrefolding portions (e.g., the red domain in Figure 6) to avoid being threaded, and instead be allowed to bypass. The biophysical cues that could signal Hsp104 to toggle from a processive form^66, 67^ to another that enables clients to bypass^68^ are currently unknown. Elucidating how these molecular machines alternate between seemingly contradictory functions will be important to understanding how biological refolding systems can work so effectively.

## Methods

### Culturing of *S. cerevisiae* (strain BY4741) Cells for Aggregation, PMSF Inhibition Screening and Limited-Proteolysis Mass Spectrometry (LiP-MS) Studies

3 × 100 mL of SC Complete media (Sunrise Sciences) were inoculated with *S. cerevisiae* (strain BY4741) from frozen cell stocks (−80°C) and grown overnight at 30°C in 250 mL Erlenmeyer flasks with agitation (300 rpm) to a final OD600 of ∼0.8. Culture from each replicate was then divided into 2 × 50 mL falcon tubes and collected by centrifugation at 3000 *g* for 90 s at 4°C. Supernatants were removed and cells were washed with 2 mL of ice cold Tris lysis buffer (20 mM Tris pH 8.2, 100 mM NaCl, 2 mM MgCl2) and combined before centrifuging again at 3000 *g* for 1 min at 4 °C. Washed cell pellets were then stored at -20°C for downstream studies.

## Methods to Study Aggregation

### 1a. Preparation of Normalized Lysates

Frozen cell pellets were resuspended in 1.4 mL of ice-cold Tris lysis buffer (supplemented with DNase I to a final concentration (f.c.) of 0.1 mg mL-1). Resuspended cells were flash frozen by slow drip over liquid nitrogen and cryogenically pulverized with a freezer mill (SPEX Sample Prep) over 8 cycles consisting of 1 min of grinding (9 Hz), and 1 min of cooling. Pulverized lysates were transferred to 50 mL centrifuge tubes and thawed at room temperature for 20 min. Lysates were then transferred to fresh 1.5 mL microfuge tubes and clarified at 16000 g for 15 min at 4 °C to remove insoluble cell debris. The cell lysates were then split into two aliquots, where one aliquot was subjected to ribosomal depletion and the other aliquot was not and kept on ice. To deplete ribosome particles, clarified lysates were transferred to 3 mL konical tubes and ultracentrifuged at 33,300 rpm at 4°C for 90 min without sucrose cushions using a SW55 Ti rotor. The supernatants were then transferred to fresh 1.5 mL microfuge tubes and then protein concentrations of the clarified lysates were determined using the bicinchoninic acid assay (Rapid Gold BCA Assay, Pierce) in a microtiter format with a plate reader (Molecular Devices iD3) using BSA as a calibration standard. Using the results of the BCA assay, the clarified cellular lysates (whether ribosome depleted or not) were normalized to a protein concentration of 2.0 mg mL-1 using Tris lysis buffer.

### 1b. Preparation of Native and Refolded samples for Precipitation Studies

To prepare native samples (with or without ribosomal depletion) for precipitation studies, 28.75 µL of normalized lysates (with or without ribosomal depletion) were diluted with 471.25 µL of native dilution buffer (20 mM Tris pH 8.2, 100 mM NaCl, 2 mM MgCl2, 1.061 mM DTT, 63.66 mM GdmCl) to a final protein concentration of 0.115 mg mL^-1^. Following dilution, the final concentrations are 20 mM Tris pH 8.2, 100 mM NaCl, 2 mM MgCl2, 1 mM DTT and 60 mM GdmCl. Native samples were then stored overnight at 4°C. The refolding samples were prepared as follows: 505 µL of normalized lysate, 50 mg of solid GdmCl, and 1.25 µL of a freshly prepared 700 mM DTT stock solution were added to a fresh 1.5 mL microfuge tube, and solvent was removed using a Vacufuge Plus (Eppendorf) to a final volume of 87 µL, such that the final concentrations of all components were 11.6 mg mL^-1^ protein concentration, 6 M GdmCl, 116.1 mM Tris pH 8.2, 580.5 mM NaCl, 11.61 mM MgCl2 and 10 mM DTT. Unfolded lysates were incubated overnight at room temperature to allow for complete unfolding prior to refolding.

To prepare refolded samples (with or without ribosomal depletion), 495 µL of refolding buffer (19.03 mM Tris pH 8.2, 95.15 mM NaCl, 1.9 mM MgCl2, 0.91 mM DTT) was added to a fresh 1.5 mL microfuge tube. 5 µL of unfolded lysates (with or without ribosomal depletion) was then added to the tube containing the refolding dilution buffer and quickly mixed by rapidly vortexing, diluting the samples 100x, followed by flash centrifugation to collect the liquid to the bottom of the tube. The final concentrations after dilution were, 0.115 mg mL^-1^ protein, 20 mM Tris pH 8.2, 100 mM NaCl, 2 mM MgCl2, 60 mM GdmCl, and 1 mM DTT, compositionally identical to that of the native sample. Refolded samples were then allowed to incubate for 2 hr at room temperature to allow for proteins to refold (or precipitate). Native samples were allowed to equilibrate to room temperature for 2 hrs.

500 μL of native and refolded samples (with or without ribosomal depletion, and both at 0.115 mg mL^-1^, final protein concentration) were centrifuged at 16000 *g* for 15 mins at 4 °C to collect aggregated proteins. The supernatant was carefully removed by pipetting to not disturb the protein pellet. The pellets in all samples were washed with 500 μL of ice-cold Tris lysis buffer to reduce the interference from reducing agents from the refolding buffer with the BCA assay. The washed pellets were then resuspended in 50 μL of 8M urea in MPW and the protein concentrations were quantified with the BCA Assay as described above. Refolding reactions were conducted on biological triplicates and were probed in technical duplicate. The amount of protein in the pellet was determined using the protein concentration and the resuspension volume (50 μL) and converted to fractional precipitation by dividing by the initial amount of protein in the refolding reaction (57.5 μg). The data are reported as a mean ± standard deviations from biological triplicates, which were differentiated at the inoculation stage. Statistical significance between samples refolded were assessed using t-tests with Welch’s correction for unequal population variances as implemented in Prism 9 (GraphPad). The “precipitation” measured for the native samples were treated as the background level of the measurement because they should not possess any precipitated protein.

## 2. Limited Proteolysis Mass Spectrometry (LiP-MS) Refolding Studies

### 2a. Preparation of Normalized Lysates

Frozen cell pellets as prepared above were resuspended in 1.4 mL of Tris lysis buffer (20 mM Tris pH 8.2, 100 mM NaCl, 2 mM MgCl2) supplemented with DNase I to a f.c. of 0.1 mg mL^-1^). In samples prepared for screening of effect of PMSF on the inhibition endogenous proteases, either 14 µL of a 50 mM PMSF stock prepared in DMSO to prevent protein degradation due to endogenous proteases or 14 µL of DMSO as a control was added. For final versions of the experiments, 14 µL of a 50 mM PMSF stock prepared in DMSO was added to all samples. Normalized lysates were prepared as above, by using cryogenic pulverization, removing ribosomes by ultracentrifugation, and normalizing to 2.0 mg mL^-1^ with the BCA assay.

### 2b. Preparation of Native Samples with or without the addition of PMSF

To prepare native samples (with or without PMSF), 23 µL of normalized lysates (with or without the addition of PMSF) were diluted with 377 µL of native dilution buffer (20 mM Tris pH 8.2, 100 mM NaCl, 2 mM MgCl2, 1.061 mM DTT, 63.66 mM GdmCl) to a final protein concentration of 0.115 mg mL^-1^. Following dilution, the final concentrations are 20 mM Tris pH 8.2, 100 mM NaCl, 2 mM MgCl2, 1 mM DTT and 60 mM GdmCl. Native samples were then equilibrated by incubating for 2 hr at room temperature prior to limited proteolysis (LiP). We note that the equilibration time for the native samples is longer than our previously studies to allow for complete quenching of PMSF activity.

Refolded samples were not prepared for the PMSF protease inhibition study. Unfolded lysates (with PMSF) for refolding studies were prepared exactly as describe above and were incubated overnight at room temperature to allow for complete unfolding prior to refolding. To prepare refolded samples for refolding studies, 198 µL of refolding buffer (19.03 mM Tris pH 8.2, 95.15 mM NaCl, 1.9 mM MgCl2, 0.91 mM DTT) was added to a fresh 1.5 mL microfuge tube. 2 µL of unfolded lysates was then added to the tube containing the refolding dilution buffer and quickly mixed by rapidly vortexing, diluting the samples 100x, followed by flash centrifugation to collect the liquid to the bottom of the tube. The final concentrations after dilution were, 0.115 mg mL^-1^ protein, 20 mM Tris pH 8.2, 100 mM NaCl, 2 mM MgCl2, 60 mM GdmCl, and 1 mM DTT, compositionally identical to that of the native sample. Refolded samples were then allowed to incubate for either (1 min, 5 min, or 2 hr) at room temperature to allow for proteins to refold prior to limited proteolysis.

### 2c. Limited Proteolysis Mass Spectrometry (LiP-MS) Sample Preparation

To perform LiP, 2 µL of a PK stock (prepared as either a 0.25 mg mL^-1^ PK for the PMSF proteinase inhibition study, or a 0.115 mg mL^-1^ PK for refolding studies, both in a 1:1 mixture of lysis buffer and 20% glycerol, stored at -20°C and thawed at most only once) were added to a fresh 1.5 mL microfuge tube. After refolded proteins were allowed to refold for the specified amount of time (1 min, 5 min, or 2 h), or native proteins were allowed their 2 hr equilibration, 200 µL of the native lysates (with or without PMSF, all production experiments used PMSF) were added to the PK-containing microfuge tube and quickly mixed by rapid vortexing (enzyme:substrate ratio is a 1:50 w/w ratio in the PMSF proteinase inhibition study, 1:100 w/w ratio in the refolding studies), followed by flash centrifugation to collect liquids to the bottom of the tube. Samples were incubated for exactly 1 min at room temperature before transferring them to a mineral oil bath preequilibrated at 110°C for 5 min to quench PK activity^18^. For samples designated as non-LiP controls for the PMSF protease inhibition study, the same procedure was used as above, except PK was not added. Boiled samples were then flash centrifuged (to collect any condensation on the sides of tube) and transferred to 152 mg of urea such that the final urea concentration was 8 M and the final volume was 316 µL. This method generates 4 native samples for the PMSF protease inhibition study: 2 samples which were lysed with PMSF in buffer and incubated with PK or not, and 2 samples which were lysed without PMSF in buffer and incubated with PK or not.

The final datasets for the refolding studies consisted of a total of 30 samples, which comprise the following, each with three biological replicates each time it was performed: native lysates (for two performances of the experiment), lysates refolded for 1 min (for three performances of the experiment), lysates refolded for 5 min (for three performances of the experiment), and lysates refolded for 2 h (for two performances of the experiment). Similar to a previous study^18^, we opted to **not** utilize a series of parallel non-LiP control samples in which PK was withheld for normalization purposes, as it was deemed not to be needed since native and refolded samples are sourced from the same lysates so it is not possible for protein abundances to be different between them. During the first performance of the experiment, non-LiP controls *were* generated and used to normalize protein abundances (as described in ref. 19). However, we found that the statistical trends were *weaker* than if normalization was *not* carried out. This is presumably because whatever small benefit is accrued from normalization is outweighed by the propagation of error (and false discoveries) from the non-LiP study. Hence, we report all data without normalization and did not perform non-LiP controls for the second and third performance of the experiment.

All protein samples were prepared for mass spectrometry as follows: 4.5 μL of a freshly prepared 700 mM stock of DTT were added to each sample-containing microfuge tube to a final concentration of 10 mM. Samples were incubated at 37°C for 30 minutes at 700 rpm on a thermomixer to reduce cysteine residues. 18 μL of a freshly prepared 700 mM stock of iodoacetamide (IAA) were then added to a final concentration of 40 mM, and samples were incubated at room temperature in the dark for 45 minutes to alkylate reduced cysteine residues. 942 μL of 100 mM ammonium bicarbonate (pH 8) were added to the samples to dilute the urea to a final concentration of 2 M. 1 μL of a 0.46 μg μL^-1^ stock of Trypsin (NEB) were added to the samples (to a final enzyme:substrate ratio of 1:50 w/w) and incubated overnight (15-16 h) at 25°C at 700 rpm (not 37 °C, so as to minimize carbamylation of lysines). After digestion, samples were prepared for MS analysis as described below, in section 7.

## 3. Ncrassa Methods

### 3a. Culturing of *N. crassa* (strain 4637A) Cells for Limited-Proteolysis Mass Spectrometry (LiP-MS) Studies

*N. crassa* spores were received from ATCC (product #42875), rehydrated in 0.5 mL sterile Millipore water, and allowed to sit overnight at room temperature to rehydrate. Rehydrated spores were plated on Neurospora culture agar (ATCC Medium 331) containing 1% sorbose and incubated at 24°C for 2 days. 5 mL of Neurospora liquid culture media (ATCC Medium 331 without agar) was inoculated using a single colony from the sorbose plate and incubated at 24°C with agitation (300rpm) overnight. 4 x 5 mL of Neurospora liquid culture media were inoculated with material from the initial liquid culture and incubated overnight at 24°C with agitation (300rpm).

### 3b. Preparation of Normalized Lysates

Solid *N. crassa* material from liquid cultures were rinsed in 1mL Tris lysis buffer (20 mM Tris pH 8.2, 100 mM NaCl, 2 mM MgCl2), placed in a new tube with 1mL Tris lysis buffer (supplemented with DNase I to a final concentration (f.c.) of 0.1 mg mL^-1^ and PMSF to a f.c. of 87μg mL^-1^) and mechanically disrupted with a spatula. Normalized lysates were prepared as above, by using cryogenic pulverization, removing ribosomes by ultracentrifugation, and normalizing to 2.0 mg mL^-1^ with the BCA assay.

### 3c. Preparation of Native Samples

To prepare native samples 11.5 µL of normalized lysates were diluted with 188.5 µL of native dilution buffer (20 mM Tris pH 8.2, 100 mM NaCl, 2 mM MgCl2, 1.061 mM DTT, 63.7 mM GdmCl) to a final protein concentration of 0.115 mg mL^-1^. Following dilution, the final concentrations are 20 mM Tris pH 8.2, 100 mM NaCl, 2 mM MgCl2, 1 mM DTT and 60 mM GdmCl. Native samples were then equilibrated by incubating for 2 hr at room temperature prior to limited proteolysis (LiP).

### 3d. Preparation of Refolded Samples

Unfolded lysates for refolding studies were prepared by adding 25 mg of GdmCl to 250μL of normalized lysate and reducing in a vacuum centrifuge to a final volume of 43 μL (6M f.c. of GdmCl) and were incubated overnight at room temperature to allow for complete unfolding prior to refolding. To prepare refolded samples for refolding studies, 198 µL of refolding buffer (19.03 mM Tris pH 8.2, 95.15 mM NaCl, 1.9 mM MgCl2, 0.91 mM DTT) was added to a fresh 1.5 mL microfuge tube. 2 µL of unfolded lysates was then added to the tube containing the refolding dilution buffer and quickly mixed by rapidly vortexing, diluting the samples 100x, followed by flash centrifugation to collect the liquid to the bottom of the tube. The final concentrations after dilution were, 0.115 mg mL^-1^ protein, 20 mM Tris pH 8.2, 100 mM NaCl, 2 mM MgCl2, 60 mM GdmCl, and 1 mM DTT, compositionally identical to that of the native sample. Refolded samples were then allowed to incubate for either 1 min, 5 min, or 2 hr at room temperature to allow for proteins to refold prior to limited proteolysis.

### 3e. Limited Proteolysis Mass Spectrometry (LiP-MS) Sample Preparation

To perform LiP, 2 µL of a PK stock (prepared as 0.115 mg mL^-1^ PK for refolding studies in a 1:1 mixture of lysis buffer and 20% glycerol, stored at -20°C and thawed at most only once) were added to a fresh 1.5 mL microfuge tube. After refolded proteins were allowed to refold for the specified amount of time (1 min, 5 min, or 2 h), or native proteins were allowed their 2 hr equilibration, 200 µL of the native lysates were added to the PK containing microfuge tube and quickly mixed by rapid vortexing (enzyme:substrate ratio is 1:100 w/w ratio in the refolding studies), followed by flash centrifugation to collect liquids to the bottom of the tube. Samples were incubated for exactly 1 min at room temperature before transferring them to a mineral oil bath preequilibrated at 110°C for 5 min to quench PK activity^18^. For samples designated as controls (native control in triplicate and refolding control in triplicate) the same procedure was used as above, except PK was not added. Boiled samples were then flash centrifuged (to collect any condensation on the sides of tube) and transferred to 152 mg of urea such that the final urea concentration was 8 M and the final volume was 316 µL. The final datasets for the refolding studies contain a total of 18 separate samples prepared for this experiment, they include: native and refolded controls (in triplicate) and native and refolded LiP, with the refolded LiP at three different timepoints.

All protein samples were prepared for mass spectrometry as follows: 4.5 μL of a freshly prepared 700 mM stock of DTT were added to each sample-containing microfuge tube to a final concentration of 10 mM. Samples were incubated at 37°C for 30 minutes at 700 rpm on a thermomixer to reduce cysteine residues. 18 μL of a freshly prepared 700 mM stock of iodoacetamide (IAA) were then added to a final concentration of 40 mM, and samples were incubated at room temperature in the dark for 45 minutes to alkylate reduced cysteine residues. 942 μL of 100 mM ammonium bicarbonate (pH 8) were added to the samples to dilute the urea to a final concentration of 2 M. 1 μL of a 0.46 μg μL^-1^ stock of Trypsin (NEB) were added to the samples (to a final enzyme:substrate ratio of 1:50 w/w) and incubated overnight (15-16 h) at 25°C at 700 rpm. After digestion, samples were prepared for MS analysis as described below in section 7.

## 4. Heat Shock Structural Perturbation Experiments

### 4a. Culturing of *S. cerevisiae* (strain BY4741) for Heat Shock Experiments

Method adapted from ref. 40. 4 × 150 mL of SC Complete media were inoculated with *S. cerevisiae* from frozen cell stocks and grown overnight at 30°C in 500 mL Erlenmeyer flasks with agitation (300 rpm) to a final OD600 of ∼0.8. Culture from each replicate was then divided into equal fractions in 3 × 50 mL falcon tubes and collected by centrifugation at 3000 *g* for 90 s at 4°C. Supernatants were then decanted leaving ∼1 mL of residual media left in each tube. For each of the biological replicates, one fraction was incubated in a water bath set to 30°C for 8 mins as the non-heat shock control samples (Ctrl), one fraction was incubated in a water bath set to 42°C for 8 mins as the heat shocked sample (HS), and the last fraction was incubated in a water bath set to 42°C for 8 mins, and then transferred to a water bath set at 30°C and incubated for 1 hr as the heat shock recovery sample (HS-R). After each sample has finished incubating in its respective condition, 1 mL of ice-cold Tris lysis buffer was added to each of the samples to quickly cool the temperature of the culture. Each sample was then centrifuged again at 3000 *g* for 1 min at 4°C. Supernatants were removed and cell pellets were then resuspended in 1.0 of ice-cold Tris lysis buffer (supplemented with DTT to a f.c. of 1 mM and DNase I to a f.c. of 0.1 mg mL^-1^). Resuspended cells were flash frozen by slow drip over liquid nitrogen and stored at -80°C until preparations of all samples and replicates were finished. We note that although samples were stored as cell pellets at -20°C until further use for the refolding studies, we did **not** store our samples as cell pellets but rather as flash-frozen cell suspensions in lysis buffer for these studies.

### 4b. Preparation of Normalized Lysates for Heat Shock Experiments

Frozen resuspended cells were cryogenically pulverized as described above. Pulverized lysates were transferred to 50 mL centrifuge tubes and thawed at room temperature for 20 min. Lysates were then transferred to fresh 1.5 mL microfuge tubes and clarified at 3000 *g* for 90 s at 4 °C to remove insoluble cell debris while keeping any biomolecular condensates that may have formed during heat shock in the supernatant. The supernatant was then transferred to a fresh 1.5 mL microfuge tube. For these experiments, careful consideration was taken to minimize the amount of time between the lysis of cells and LiP with PK. As a result, protein concentration was determined by the absorbance value obtained at 280 nM (A280) instead of BCA assays as above. Generally, the raw A280 value would be between 12-13 for the various preparations. Protein concentrations were diluted to an A280 value of 6 (determined to be approximately 1.0 mg mL^-1^ in a separate experiment correlating A280 values to known protein concentrations as determined by BCA assay) in Tris lysis buffer supplemented with 1 mM DTT.

### 4c. LiP-MS of Heat Shock Samples

To perform LiP, 2.5 µL of a PK stock (prepared as a 1 mg mL^-1^ Proteinase K in a 1:1 mixture of lysis buffer and 20% glycerol, stored at -20°C and thawed at most only once) stock was added to a fresh 1.5 mL microfuge tube. 250 µL of the normalized lysates (Ctrl, HS, or recovered) were then added to the PK-containing microfuge tube and quickly mixed by rapid vortexing (enzyme:substrate ratio is a 1:100 w/w ratio), followed by flash centrifugation to collect liquids to the bottom of the tube. Samples were incubated for exactly 1 min at room temperature before transferring them to a mineral oil bath preequilibrated at 110°C for 5 min to quench PK activity. As above, boiled samples were then flash centrifuged (to collect any condensation on the sides of tube) and transferred to a fresh 2.0 mL microfuge tube containing 190 mg of urea such that the final urea concentration was 8 M and the final volume was 392.5 µL. For samples designated as controls, the same procedure was used as above, except PK was not added. In total, 24 separate samples were prepared for this experiment, they include: the non-heat shock control, heat shock, and heat shock recovery conditions prepared as both LiP and non-LiP samples, and the appropriate biological quadruplicates for each category. We note that here, we chose to include parallel “control” samples in which PK is withheld to quantify protein abundance differences during the heat shock and heat shock recovery processes, to be used for normalization.

All protein samples were prepared for mass spectrometry as above: 5.6 μL of a freshly prepared 700 mM stock of DTT were added to each sample-containing microfuge tube to a final concentration of 10 mM. Samples were incubated at 37°C for 30 minutes at 700 rpm on a thermomixer. 22.4 μL of a freshly prepared 700 mM stock of iodoacetamide (IAA) were then added to a final concentration of 40 mM, and samples were incubated at room temperature in the dark for 45 minutes. 1.26 mL of 100 mM ammonium bicarbonate (pH 8) were added to the samples to dilute the urea to a final concentration of 2 M. 5 μL of a 1.0 μg μL^-1^ stock of Trypsin (NEB) were added to the samples (to a final enzyme:substrate ratio of 1:50 w/w) and incubated overnight (15-16 h) at 25°C at 700 rpm. After digestion, samples were prepared for MS analysis as described below in section 7.

## 5. Stress Granule Dispersal Experiments

### 5a. Engineering a Fluorescent and Affinity Tagged Pab1 Strain

An *S. cerevisiae* (strain BY4741) containing a C-terminal Clover-labeled Pab1 at the chromosomal locus, a gift from the Drummond Lab (University of Chicago), was transformed with a Clover-FLAG-Leu2d gene block (see Table S2) ordered from IDT with homology arms to the chromosomal locus to create a fluorescent protein-tagged Pab1 gene that is also affinity tagged (1x FLAG) at the native locus. In addition, the Leu2d gene was inserted into the chromosomal locus downstream of Pab1 to allow for positive selection on SD(–Leu) agar plates. To perform genomic knock-in by homologous recombination, *S. cerevisiae* cells harboring Pab1-Clover were grown in 5 mL of YPD from saturated overnight cultures with a starting OD600 of 0.05 in 14 mL round bottom falcon tubes. Cells were cultured at 30°C with agitation (300 rpm) to log phase at a final OD600 of 0.4, before being collected by centrifugation at 2000 *g* for 5 mins at 4°C. The supernatant was removed, and the cell pellet was resuspended with 1 mL of TE-LiOAc (10 mM Tris pH 8.0, 1 mM EDTA pH 8.0, 0.1 M LiOAc) and transferred to a fresh 1.5 mL microfuge tube and centrifuged again at 2000 *g* for 5 min at 4°C. Supernatants were removed and cells were resuspended in 100 µL of TE-LiOAc. 10 µL of denatured salmon sperm (incubated at 95°C for 10 mins), 25 µL of 100 ng/µL DNA (PCR amplified using Clover FLAG-Leu2d gene block using Q5 DNA polymerase (NEB) according to the manufacturer’s protocol), and 800 µL of PEG-TE-LiOAc (10 mM Tris pH 8.0, 1 mM EDTA pH 8.0, 0.1 M LiOAc, 40% PEG (3350 mw)) were added to resuspended cells in that order and mixed thoroughly between each addition. Cells were incubated at 30°C for 30 mins with agitation (300 rpm) in a thermomixer and then heat-shocked 30 mins in a 42°C water-bath. Cells were then harvested by centrifugation at 14000 *g* for 30 seconds. The supernatants were removed, and cells were washed with 1 mL of water and then pelleted at 3000 *g* for 5 mins. Washed cell pellets were then resuspended in 100 µL of water before being plated on selective media (SD-leu) and allowed to grow for 2 days at 30°C. Colonies from this plate were genotyped by colony PCR followed by Sanger sequencing of the PCR amplicon. A clone positive for recombination was then used to prepare frozen cell stocks by inoculating the colony into 5 mL of SD(-Leu), growing to saturation, adding an equal volume of 20% glycerol in SD(-Leu), aliquoting into 1.5 mL microfuge tubes, and flash freezing. For subsequent downstream experiments, one frozen aliquot of cell stocks was streaked on a SD(-Leu) agar plate, incubated for 2 days at 30°C, and the colonies used to seed overnight cultures.

### 5b. Culturing and Heat Shock of Pab1-Clover-Flag

3 × 400 mL of YPD media were inoculated with *S. cerevisiae* (containing fluorescent and affinity tagged Pab1) from saturated overnight cultures with a starting OD600 of 0.2. Cells were then cultured at 30°C with agitation (300 rpm) to a final OD600 of 0.8 before being combined into one large 1.2 L culture in a 2 L baffled flask. OD600 was remeasured to confirm that it was still at an OD600 of 0.8. Cells were then transferred to 8 × 50 mL falcon tubes and collected by centrifugation at 4000 *g* for 1 min at 25°C. Supernatants were removed and an additional 50 mL of cells were transferred to each 50 mL falcon tube and collected again by centrifugation at 4000 *g* for 1 min at 25°C. This step was repeated once more such that after all centrifugations, each 50 mL falcon tube contained the equivalent of 150 mL of culture at an OD600 of 0.8. Each of the cell pellets were then resuspended in 1 mL of YPD and subjected to different treatments. This experiment was conducted twice, with differences in heat shock temperature and recovery times. For both experiments, one falcon tube was incubated for 15 mins in a 30°C water bath, serving as the non-heat shock sample. The other 7 falcon tubes were incubated for 15 mins in either a 42°C or 46°C water bath. After incubation, 1 mL of ice cold stress granule (SG) lysis buffer (50 mM Tris pH 7.4, 100 mM Potassium acetate, 2 mM Magnesium Acetate, 0.5 mM DTT, 0.5% NP40, 50 µg / mL Heparin and 1x Protease Inhibitor Cocktail (1 mM AEBSF.HCl, 0.8 µM Aprotonin, 50 µM Bestatin, 15 µM E-64, 20 µM Leupeptin, and 10 µM Pepstatin A)) was added to the non-heat-shocked culture and one to the heat-shocked culture. The two tubes were then centrifuged at 4000 *g* for 1 min at 4°C and the supernatants were removed. The cell pellets were resuspended in 1.5 mL of ice-cold SG lysis buffer supplemented with 37.5 µL of RNase Inhibitor (Murine) to prevent RNA degradation after lysis. Resuspended cells were flash frozen by slow drip over liquid nitrogen and stored at -80°C until preparation of all samples and replicates were finished.

The remaining 6 heat shocked samples were then quickly transferred to 150 mL of fresh YPD in a 500 mL Erlenmeyer flask preequilibrated at 30°C and recovered in an orbital shaker at 30°C with agitation (300 rpm). Each culture was allowed to recover for different time intervals (the time between the heat-shocked cultures being transferred to YPD until being removed from the orbital shaker) before being removed and split into 3 × 50 mL falcon tubes. The 3 tubes were rapidly transferred to a centrifuge, and spun at 4000 *g* for 1 min at 4°C and the supernatants were quickly removed. Each tube was resuspended in 500 µL of stress granule lysis buffer and pooled into one 50 mL falcon tube. 37.5 µL of RNase Inhibitor (Murine) was added. As above, resuspended cells were flash frozen by slow drip over liquid nitrogen and stored at -80°C until all recovery times finished. In total, 8 samples were generated for each heat shock temperature: a non-heat shock (control) sample, a heat shock sample where the cells were not allowed to recover, and 6 heat shock samples where the cells were allowed to recover for different amounts of time. For the samples heat shocked at 42°C, the time points were 5 min, 15 min, 30 min, 45 min, 60 min, and 90 min. For the samples heat shocked at 46°C, we note that there were two separate performances of the experiment, one in which cells were allowed to recover for 15 min, 30 min, 45 min, 60 min, 75 min and 90 min; a second in which cells were allowed to recover for 2 min, 15 min, 30 min, 45 min, 60 min and 90 min. We note that due to the number of samples that need to be processed simultaneously, these experiments require three experimentalists working simultaneously.

### 5c. Isolation of Stress Granule Fractions

Frozen cell suspensions were cryogenically pulverized as described above. Pulverized lysates were transferred to a 50 mL centrifuge tubes and thawed at room temperature for 20 min. Lysates were then transferred to fresh 1.5 mL microfuge tubes and clarified by centrifugation at 18000 *g* for 10 mins at 4°C. The supernatant was then discarded, and the remaining pellet was resuspended in 1 mL of stress granule lysis buffer. The resuspended pellet was then centrifuged at 14000 *g* for 10 min at 4°C and the supernatant was again discarded. The remaining pellet was resuspended in 150 µL of stress granule lysis buffer and centrifuged at 850 *g* for 2 min at 4°C. The supernatant, enriched for heat stress granules, was then transferred to a fresh 1.5 mL microfuge tube and diluted with 350 µL of SG lysis buffer to a final volume of 500 µL. 5 µL of RNase inhibitor was added to each sample. This generates the heat stress granule fraction for each condition.

### 5d. Absorbance, Fluorescence and Protein Quantification

3 × 125 µL of the heat stress granule fraction was transferred to black-walled clear-bottom 96 well plates creating technical triplicates for each sample. Absorbance readings at 342 nm and Clover fluorescence (excitation at 480 nm, emission at 516 nm, with an integration time of 40 µs obtained from 20 mm from the plate) were obtained using a Tecan Spark Plate Reader. Relative fluorescence was determined by taking the ratio of the fluorescence for a given sample by the highest fluorescence value obtained. Protein concentrations of the stress granule enriched samples were determined using the bicinchoninic acid assay (BCA Protein Assay, Pierce) in a microtiter format with a plate reader (Molecular Devices iD3) using BSA as a calibration standard. We note here that we use the BCA assay kit to determine the protein concentration as the rapid gold BCA assay kit is incompatible with DTT.

### 5e. Microscopy Analysis

125 µL of samples were transferred to an 8 well coverglass (Nunc Lab-Tek II Chambered Coverglass, Thermofisher). Images were captured with a vt-instant Structured Illumination Microscope (vt-iSIM; BioVision Technologies) using the 488nm 150Mw laser, an ORCA-Fusion sCMOS camera and the Leica HC PL APO 63x/1.40 OIL CS2 objective. All images were obtained with a laser power of 30% with a 150 ms exposure time. Fluorescence images were analyzed using the Fiji software^70^. For the heat shock experiment conducted at 42°C, particle sizes, average intensities were determined using the “Analyzer Particles” function in FIJI with a pixel size greater than 20 and threshold set to 135. Samples heat shocked at 46°C were identified the same way, except that the threshold was set to 170 for samples with recovery times 0 min, 2 min and 15 min, and 135 for all subsequent recovery times. Particle sizes were determined by counting the number of pixels obtained for each stress granule and multiplying by the area of each pixel (0.0106 µm^2^).

To estimate the number of stress granules per cell we used the following heuristic. The particles/image was multiplied by the area ratio (area of well/area of field of view = 0.9 cm^2^/2.76 ×10^-5^ cm^2^) and the height ratio (height of sample in well/heigh of illuminated volume = 1.4 mm / 100 nm) to give the total particles/well. To estimate the number of yeast cells whose lysates entered the well, we multiplied the percent of the heat shock granule fraction that got loaded into the microscope (125 µL/500 µL) by the recovery yield of the lysis (80%) by the number of cells in the culture (1.8 × 10^9^) = 3.6 × 10^8^. As the conversion factor of particles/image to particles/well (4.6 × 10^8^) is very similar to the number of cells whose lysates entered the well, the particles/image roughly equals the number of stress granules per cell.

### 5f. Immunoprecipitation of Pab1-Clover-FLAG and Mass Spec Prep

Anti-FLAG M2 magnetic beads (Sigma) were prepared by thoroughly resuspending and then taking the 30 µL of slurry and transferring into a fresh 1.5 mL microfuge tube. The tube was placed into a magnetic separator and the supernatant was removed leaving behind 15 µL of beads. Beads were equilibrated twice with 150 µL of SG lysis buffer. 250 µL of stress granule enriched samples were then added to the beads incubated overnight (∼16 hr) at 4°C on a tube rotator. The next day, beads were washed twice with by removing the flow through, adding 500 µL of a high salt SG lysis buffer (50 mM Tris pH 7.4, 300 mM Potassium acetate, 2 mM Magnesium Acetate, 0.5 mM DTT, 0.5%NP40, 50 µg / mL Heparin) and incubating on a tube rotator for 5 mins at 4°C between each wash step. Beads were then washed twice again as above with 500 µL of a high salt SG lysis buffer but withholding DTT, NP40 and Heparin (50 mM Tris pH 7.4, 300 mM Potassium acetate, 2 mM Magnesium Acetate). The heat shock stress granules were then eluted by adding 150 µL of an elution buffer (8 M Urea, 50 mM Ambic) to the beads and incubating for 45 mins at 4°C on a tube rotator. The elute was then transferred to a fresh 1.5 mL microfuge tube and prepared for mass spec as above. 2.14 μL of a freshly prepared 700 mM stock of DTT were added to each sample containing microfuge tube to a final concentration of 10 mM. Samples were incubated at 37°C for 30 minutes at 700 rpm on a thermomixer. 8.56 μL of a freshly prepared 700 mM stock of iodoacetamide (IAA) were then added to a final concentration of 40 mM, and samples were incubated at room temperature in the dark for 45 minutes. 450 µL of 50 mM ammonium bicarbonate (pH 8) were added to the samples to dilute the urea to a final concentration of 2 M. 1 μL of a 0.2 μg μL^-1^ stock of Trypsin (NEB) were added to the samples (to a final enzyme:substrate ratio of 1:50 w/w) and incubated overnight at 25°C at 700 rpm. After digestion, samples were prepared for MS analysis as described below in section 7.

We note that as based on quantification assays (section 5d) and the capacity of the anti-FLAG magnetic beads, the input was in high excess such that all elutions are expected to produce similar overall levels of protein (Extended Data Figure 12A-B). Hence to convert mass spec-derived abundances to relative abundance of each protein in heat stress granules relative to the initial time-point, they were multiplied by the ratio of overall protein amount at the given time point divided by overall protein amount at initial time point (Extended Data Figure 12C-D).

## 6. Hom6p, Trp5p and Abp1p Western Blot Experiments

### 6a. Cloning pESC-Leu2d-Trp5p, pESC-Leu2d-HOM6p, and pESC-Leu2d-ABP1p Construct

To create plasmids to express full length Trp5p, Hom6p, or Abp1p proteins that are C-terminally His tagged with a short GSSGSS linker separating the two, geneblocks were ordered from IDT (Table S1) containing the full-length protein of interest, a ∼15 bp or 18 bp homology arm at the 5’ or 3’ end with pESC_Leu2d at the 5’ end, and either a short GSSGSS linker serving as the homology arms at the 3’end (for geneblocks used to insert Hom6p or Abp1p) or a GSSGSS linker followed by a His tag, a stop codon at the end of the His tag and homology arms to the pESC_Leu2d vector at the 3’end (for geneblock used to insert Trp5). To create pESC_Leu2d-Trp5p, a linearized pESC-Leu2d vector was generated using pESC_Leu2d as a template for PCR with primers pESC_Leu2d_f and pESC_Leu2d_r (Table S1) using the Phusion polymerase (Thermofisher) according to the manufacturer’s protocol. Purity and accuracy of PCR products was assessed using agarose gel electrophoresis (0.8% agarose, 0.5 x TBE), DpnI digested to remove template DNA (37°C for 30 mins and then 80°C for 20 mins to inactivate DpnI), and then purified by column purification using a DNA Clean up and Concentration kit (Zymo) according to manufacturer’s protocol. DNA concentration was then quantified by nanodrop using a Nanodrop OneC (ThermoFisher). The geneblock was then ligated to the pESC_Leu2d_vector using Gibson assembly using HiFi 2 × Gibson Master Mix (NEB) according to manufacturer’s protocol and transformed into chemically competent 10beta cells. Transformation was accomplished by taking 1 µL of ligated DNA and combining with 25 µL of chemical component cells, incubating on ice for 25 mins and then subjecting cells to a heat pulse in a 42°C water bath for 40 s. Cells were then returned to ice for 2 mins and recovered by inoculating into 1 mL of SOC media and incubated at 37°C for 1 h with agitation (700 rpm in a thermomixer). 100 µL of the transformant was plated on selective media (LB agar supplemented with 100 µg mL^-1^ of Ampicillin) using sterile coli rollers and incubated at 37°C overnight (∼16 hr). 5 × 5 mL of selective media (LB supplemented with 100 µg mL^-1^ of Ampicillin) in 14 mL round-bottom flasks were then each inoculated with single colonies and incubated overnight at 37°C with agitation (220 rpm). The next day, saturated overnight cultures were centrifuged at 3000 *g* for 15 min at 4°C and plasmid DNA was extracted using a ZymoPURE Plasmid Miniprep Kit (Zymo) according to the manufacturer’s protocol. Plasmid DNA were sent for Sanger sequencing (Genewiz) according to Genewiz specifications. A single tube containing the correct DNA sequencing would then be used for all subsequent transformations. The same protocol above was used to generate pESC_Leu2d_Hom6p and pESC_Leu2d_Abp1p except that a pESC_Leu2d vector containing a His tag was generated using pESC_Leu2d-Trp5 as template for PCR with primers pESC_Leu2d_His_f and pESC_Leu2d_r.

### 6b. Creating Truncations of pESC_Leu2d_Trp5 and pESC_Leu2d_ABP1

To create a plasmid to express Trp5p with its structured linker removed (Trp5p_RL), pESC_Leu2d_Trp5p was used as a template for PCR with primers pESC_Leu2d_Trp5p_RL_r and pESC_Leu2d_Trp5p_RL_f using the Phusion polymerase as above. The amplified DNA was ligated using the QuickChange method by directly transforming DNA obtained from PCR into chemical competent 10-beta cells as described above. To create a plasmid to express Trp5p where its structure linker has been replaced with a flexible linker of similar length (Trp5p_GSS), a linearized pESC-Leu2d-Trp5p vector with its native linker removed was generated using pESC_Leu2d-Trp5p as a template for PCR with primers pESC_Leu2d_Trp5p_GSS_f and pESC_Leu2d_Trp5p_GSS_r using the Phusion polymerase as described above. A geneblock ordered from IDT containing a 16× GSS flexible linker with homology arms at the 5’ and 3’ end with Trp5p with ligated with the pESC_Leu2d_Trp5p vector using Gibson assembly as described above and transformed into chemical competent 10-beta cells as described above. To create a plasmid to express Abp1p with its structured linker removed (Abp1p_RL), pESC_Leu2d_Abp1p was used as a template for PCR with primers pESC_Leu2d_Abp1p_RL_r and pESC_Leu2d_Abp1p_RL_f using the Phusion polymerase as above. The amplified DNA was ligated using the QuickChange method by directly transforming DNA obtained from PCR into chemical competent 10-beta cells as described above.

### 6c. Transformation of plasmids into *S. cerevisiae* (strain BY4741)

The 6 different plasmids (pESC_Leu2d_Hom6p, pESC_Leu2d_Trp5p, pESC_Leu2d_Trp5p_RL, pESC_Leu2d_Trp5p_GSS, pESC_Leu2d_Abp1p, pESC_Leu2d_Abp1p_RL) were transformed using the same protocol for yeast homologous recombination as described above using 1 µL of plasmid DNA. This generates 6 separate plates of yeast cells, each plate containing yeast colonies harboring one of the 6 different plasmids.

### 6d. Cell culture and Expression

10 mL of SD-Leu media were inoculated with transformed *S. cerevisiae* (harboring the appropriate pESC_Leu2d plasmid) at 30°C with agitation (300 rpm) and grown overnight (∼16 hr) to saturation. To express Hom6p, Trp5p, Trp5p_RL and Trp5p_GSS proteins, 3 x 50 mL of SD-Leu-glucose (+2% Galactose / + 2% Raffinose) were inoculated from overnight saturated cultures and grown for 16 hr at 30°C with agitation (300 rpm) as follows: a volume of the overnight saturated cultures was selected such that expression cultures would commence at an OD600 of 0.1. This volume was transferred to fresh 1.5 mL microfuge tubes and centrifuged at 3000 *g* for 5 mins at 4°C and the supernatants were discarded. The cell pellets were then resuspended in 1 mL of SD-Leu-glucose (+2% Galactose / + 2% Raffinose) before being transferred to 49 mL of the same media in a 250 mL Erlenmeyer flask and cultured for 16 hr at 30°C with agitation (300 rpm). This is done to minimize the amount of glucose in the cell culture as it is an inhibitor of the Gal,1,10 promoter. Generally, after 16 hr of growth, cells reach an OD600 of 2.0. After overnight expression, cells were transferred into 50 mL falcon tubes and collected by centrifugation at 3000 *g* for 90 s at 4°C. Cells were washed with 2 mL of ice-cold Tris lysis buffer and centrifuged again at 3000 *g* for 1 min at 4°C. Cell pellets were then resuspended in 2 mL of ice-cold Tris lysis buffer (supplemented with DNase to a f.c. of 0.1 mg mL^-1^ and PMSF to a f.c. of 0.5 mM) and flash frozen by slow drip over liquid nitrogen. Abp1p and Abp1p_RL were expressed the same way except were cultured in 3 x 150 mL of SD-leu-glucose (+2% Galactose / + 2% Raffinose). For cultures expressing Abp1p and Abp1p_RL, final OD600 values of 0.6 were typically obtained, hence larger culture volumes were needed to get the same amount of protein.

### 6e. Preparation of Native Sample and Refolded Samples

Cell lysates were prepared as previously, by performing cryogenic pulverization, thawing, and normalizing to 3.5 mg mL^-1^ using the Rapid Gold BCA assay. Ribosomal depletion with ultracentrifugation was not carried out (based on results showing it to be unnecessary, Figure 3A). To prepare native samples of each protein, 33 µL of normalized lysates were diluted with Tris dilution buffer (20 mM Tris pH 8.2, 100 mM NaCl, 2 mM MgCl2, 1.034 mM DTT, 62.04 mM GdmCl) to a final protein concentration of 0.115 mg mL^-1^. Following dilution, the final concentrations are 20 mM Tris pH 8.2, 100 mM NaCl, 2 mM MgCl2, 1 mM DTT and 60 mM GdmCl. Native samples were incubated for 2 hr at room temperature prior to further steps.

The refolding samples were prepared as follows: 288 µL of normalized lysate, 50 mg of solid GdmCl, and 1.25 µL of a freshly prepared 700 mM DTT stock solution were added to a fresh 1.5 mL microfuge tube, and solvent was removed using a Vacufuge Plus (Eppendorf) to a final volume of 87 µL, such that the final concentrations of all components were 11.6 mg mL^-1^ protein concentration, 6 M GdmCl, 66.3 mM Tris pH 8.2, 331.4 mM NaCl, 6.63 mM MgCl2 and 10 mM DTT. Unfolded lysates were incubated overnight at room temperature to allow for complete unfolding prior to refolding. To prepare refolded samples for aggregation, 990 µL of refolding buffer (19.53 mM Tris pH 8.2, 97.65 mM NaCl, 1.95 mM MgCl2, 0.91 mM DTT) was added to a fresh 1.5 mL microfuge tube. 10 µL of unfolded lysates was then added to the tube containing the refolding dilution buffer and quickly mixed by rapidly vortexing, diluting the samples 100x, followed by flash centrifugation to collect the liquid to the bottom of the tube. The final concentrations after dilution were, 0.115 mg mL^-1^ protein, 20 mM Tris pH 8.2, 100 mM NaCl, 2 mM MgCl2, 60 mM GdmCl, and 1 mM DTT, compositionally identical to that of the native sample. Refolded samples were then allowed to incubate for 2hr at room temperature to allow for proteins to refold prior to aggregation studies.

### 6f. Aggregation Studies by BCA and Western Blot analysis

After incubation times, native and refolded samples were mixed by vortexing and then 25 µL of the samples was transferred to a fresh 1.5 mL microfuge tube, creating the total (T) fraction. The remaining 975 µL of native and refolded samples were then centrifuged at 16000 *g* for 15 mins at 4°C to collect any aggregated proteins. The supernatants were carefully removed by pipetting to not disturb the protein pellet and transferred to a fresh 1.5 mL microfuge tube. This is the supernatant (S) fraction. The pellets were washed with 1 mL of ice-cold Tris lysis buffer to reduce the interference from reducing agents from the refolding buffer with the BCA assay. The washed pellets were then resuspended in 100 µL of 8M Urea. 50 µL of the resuspended pellet was removed and transferred to a fresh 1.5 mL microfuge tube for western blot analysis. This is the pellet (P) fraction. The protein concentration of the remaining 50 µL of resuspended pellet was quantified by BCA Assay as describe above. The amount of protein in the pellet was determined using the protein concentration and the total resuspension volume (100 µL) and then converted to fractional precipitation by diving by the initial amount of protein the refolding reaction (115 µg) and statistical significance was determined as described above.

Western blot analysis for the Total (T), Supernatant (S) and Pellet (P) were conducted as follows: 20 μL of each resuspension was transferred to a microfuge tube, mixed with 5 μL of 5× Tris-glycine-SDS loading buffer via vortexing, heated in a 90 °C water bath for 5 min. 25 μL of each lysate was loaded onto precast Novex WedgeWell 4–12% Tris-glycine Mini protein gels (ThermoFisher Scientific) with 4 μL of prestained PAGE ruler as the ladder (ThermoFisher Scientific; 26619) and was separated by electrophoresis at 200 V for approximately 40 min, using 1× Tris-glycine-SDS electrophoresis running buffer (BioRad). The resulting gel was incubated for approximately 5 min in 0.8× Tris-glycine buffer (BioRad) and 20% (v/v) methanol. The gel was trimmed to remove the wells and foot, and electroblotting was performed using an iBlot 2 gel transfer device (ThermoFisher) and its matching transfer packet (Invitrogen; iBlot 2 Transfer Stacks, PVDF, regular size, according to the manufacturer’s protocol (7 min: 20 V). After electroblotting, the PVDF membrane was incubated in 15 mL of 5% (w/v) nonfat milk (Nestle—Carnation, instant nonfat dry milk)–TBST solution for approximately 1 h with rocking to block the membrane. 5% nonfat milk–TBST solution was made by combining 1× TBS (Quality Biological; pH 7.4) with evaporated milk and Tween 20 to a final concentration of 0.1% (v/v). The blocked membrane was incubated ∼16 h in 8 mL of diluted primary anti-His antibody (mouse; Invitrogen) solution at 4 °C with rocking; the diluted primary antibody was made by diluting the antibody by 1:2000 in 5% (w/v) nonfat milk–TBST. The membrane was rinsed in 1× TBST 3 times for 10 min each and incubated in 8 mL of diluted secondary antimouse-HRP antibody (goat; Invitrogen) solution at room temperature for 40 min with rocking; the diluted secondary antibody was made by diluting the antibody 1:10 000 in 5% (w/v) nonfat milk–TBST. The incubated membrane was rinsed in 1× TBST 3 times for 10 min each, incubated for 1 min in 4 mL of chemiluminescence reagents (Signal West Femto Maximum Sensitive Substrate; ThermoFisher Scientific) that were mixed in a 1:1 ratio, and then, images were acquired using a ChemiDoc Touch imaging system (BioRad). Gel images were quantified by taking the ratio of the different conditions of the ChemiDoc Touch Images Gel.

## 7. MS Data Analysis and Computational Workflows

### 7a. Solid Phase Extraction and Sample Storage

Peptides were desalted with Sep-Pak C18 1 cc Vac Cartridges (Waters) over a vacuum manifold. Tryptic digests were first acidified by addition of trifluoroacetic acid (TFA, Acros) to a final concentration of 1% (vol/vol). Cartridges were first conditioned (1 mL 80% ACN, 0.5% TFA) and equilibrated (4 x 1 mL 0.5% TFA) before loading the sample slowly under a diminished vacuum (ca. 1 mL/min). The columns were then washed (4 x 1 mL 0.5% TFA), and peptides were eluted by addition of 1 mL elution buffer (80% ACN, 0.5% TFA). During elution, vacuum cartridges were suspended above 15 mL conical tubes, placed in a swing-bucket rotor (Eppendorf 5910R), and spun for 3 min at 350 g. Eluted peptides were transferred from Falcon tubes back into microfuge tubes and dried using a vacuum centrifuge (Eppendorf Vacufuge). Dried peptides were stored at -80°C until analysis. For analysis, samples were vigorously resuspended in 0.1% formic acid in Optima water (ThermoFisher) to a final concentration of 0.5 mg mL^-1^.

### 7b. LC-MS/MS Acquisition

Chromatographic separation of digests were carried out on a Thermo UltiMate3000 UHPLC system with an Acclaim Pepmap RSLC, C18, 75 μm × 25 cm, 2 μm, 100 Å column. Approximately, 1 μg of protein was injected onto the column. The column temperature was maintained at 40 °C, and the flow rate was set to 0.300 μL min^-1^ for the duration of the run. Solvent A (0.1% FA) and Solvent B (0.1% FA in ACN) were used as the chromatography solvents. The samples were run through the UHPLC System as follows: peptides were allowed to accumulate onto the trap column (Acclaim PepMap 100, C18, 75 μm x 2 cm, 3 μm, 100 Å column) for 10 min (during which the column was held at 2% Solvent B). The peptides were resolved by switching the trap column to be in-line with the separating column, quickly increasing the gradient to 5% B over 5 min and then applying a 95 min linear gradient from 5% B to 40% B. Subsequently, the gradient held at 40% B for 5 min and then increased again from 40% B to 90% B over 5 min. The column was then cleaned with a sawtooth gradient to purge residual peptides between runs in a sequence.

A Thermo Q-Exactive HF-X Orbitrap mass spectrometer was used to analyze protein digests. A full MS scan in positive ion mode was followed by 20 data-dependent MS scans. The full MS scan was collected using a resolution of 120000 (@ m/z 200), an AGC target of 3E6, a maximum injection time of 64 ms, and a scan range from 350 to 1500 m/z. The data-dependent scans were collected with a resolution of 15000 (@ m/z 200), an AGC target of 1E5, a minimum AGC target of 8E3, a maximum injection time of 55 ms, and an isolation window of 1.4 m/z units. To dissociate precursors prior to their reanalysis by MS2, peptides were subjected to an HCD of 28% normalized collision energies. Fragments with charges of 1, 6, 7, or higher and unassigned were excluded from analysis, and a dynamic exclusion window of 30.0 s was used for the data-dependent scans.

### 7c. LC-MS/MS Data Analysis

Proteome Discoverer (PD) Software Suite (v2.4, Thermo Fisher) and the Minora Algorithm were used to analyze mass spectra and perform Label Free Quantification (LFQ) of detected peptides. Default settings for all analysis nodes were used except where specified. The data were searched against *Saccharomyces cerevisiae* (UP000002311, Uniprot) or *Neurospora crassa* (UP000001805, Uniprot) reference proteome databases. For peptide identification, the PD MSFragger node were used, each using a semi-tryptic search allowing up to 2 missed cleavages. A precursor mass tolerance of 10 ppm was used for the MS1 level, and a fragment ion tolerance was set to 0.02 Da at the MS2 level for both search algorithms. For MSFragger, a peptide length between 7 and 50 amino acid residues was allowed with a peptide mass between 500 and 5000 Da. Additionally, a maximum charge state for theoretical fragments was set at 2 for MSFragger. Oxidation of methionine and acetylation of the N-terminus were allowed as dynamic modifications while carbamidomethylation on cysteines was set as a static modification. All parameters for MSFragger search algorithms are provided in the table below. For peptides identified, the Philosopher PD node was used for FDR validation. Raw normalized extracted ion intensity data for the identified peptides were exported from the .pdResult file using a three-level hierarchy (protein > peptide group > consensus feature). These data were further processed utilizing custom Python analyzer scripts (available on GitHub, and described in depth previously in ref. 18). Briefly, normalized ion counts were collected across the refolded replicates and the native replicates for each successfully identified peptide group. Effect sizes are the ratio of averages (reported in log2) and P-values (reported as –log10) were assessed using *t* tests with Welch’s correction for unequal population variances. Missing data are treated in a special manner. If a feature is not detected in all three native (or refolded) injections and is detected in all three refolded (or native) injections, we use those data, and fill the missing values with 1000 (the ion limit of detection for this mass analyzer); this peptide becomes classified as an all-or-nothing peptide. If a feature is not detected in one out of six injections, the missing value is dropped. Any other permutation of missing data (e.g., missing in two injections) results in the quantification getting discarded. In many situations, our data provide multiple independent sets of quantifications for the same peptide group. This happens most frequently because the peptide is detected in multiple charge states. In this case, we calculate effect size and P-value for all features that map to the same peptide group. If the features all agree with each other in sign, they are combined: the quantification associated with the median amongst available features is used and the P-values are combined with Fisher’s method. If the features disagree with each other in sign, the P-value is set to 1. Coefficients of variation (CV) for the peptide abundance in the three replicate refolded samples are also calculated. Analyzer returns a file listing all the peptides that can be confidently quantified, and provides their effect size, P-value, refolded CV, proteinase K site (if half-tryptic), and associated protein metadata.

### 7d. Refoldability Analysis

Results from analyzer are digested in the following way. Proteins with only one peptide confidently quantified are discounted; proteins with more than two are kept. Peptides are considered to have significantly different abundance in the refolded sample if the effect size is 2 or greater (more than double or less than half the abundance of native), and the P-value is less than 0.01 by Welch’s *t* test. All-or-nothing peptides must have abundance differences greater than 64-fold, and use a relaxed P-value cut-off of 0.0158. The number of significant and all-or-nothing peptides is counted for each protein. Proteins (or domains) are deemed nonrefoldable if two or more peptides with significantly different abundances in the refolded sample are identified.

Protein-level refoldability analyses proceed by counting the number of refoldable and nonrefoldable proteins within a set of categories (e.g., 5 < pI < 6) associated with a feature (e.g., pI) and calculating the fraction refolding within the category. To determine if there is a significant enrichment for (non-)refolders within certain categories, we calculate the expected number of (non-)refolders for each category by taking the total number of proteins that are assigned a value under the feature in question, times the fraction (non-)refolding, times the fraction of proteins in that category. The chi-square test is used to determine if the observed counts and expected counts significantly differ, for all cases in which the feature has three or more categories. If it only has two, Fisher’s exact test is used instead.

Domain-level refoldability is conducted in a similar way to protein-level analyses, except that instead of considering all peptides that map to a particular protein, only peptides that map to a particular domain is considered (which is defined based on its host protein and the appropriate residue range, as determined by DomainMapper^27^). Half-tryptic peptides are considered if the proteinase K cut-site lies within the domain in question. Full-tryptic peptides are considered only if the whole range of residues lie within the domain in question.

Peptide (site)-level refoldability analyses are performed in a similar way. The total number of refoldable (nonsignificant) and nonrefoldable (significant difference in abundance between refolded samples and native references) sites mapped to proteins associated with a particular feature are counted and the frequency refoldable calculated. To determine if there is a significant enrichment for (non)refoldable peptides associated with certain categories, we calculate the expected number of (non)refoldable peptides for each category by taking the total number of peptides associated with proteins that are assigned a value under the feature in question, times the fraction of peptides that are (non)refoldable, times the fraction of peptides associated with that category. The chi-square test is used to determine if the observed counts and expected counts significantly differ, for all cases in which the feature has three or more categories. If it only has two, Fisher’s exact test is used instead.

### 7e. Heat Shock Structural Perturbation Analysis

For heat shock structural perturbation analysis, we performed 12-way LFQs, these experiments were done in quadruplicates), and created a slightly modified analyzer script that assesses peptide quantifications separately for the samples associated with condition 1 (heat shocked at 42°C) and the samples associated with condition 2 (heat shocked at 42°C and recovered). In this LFQ, both conditions shared the same four native samples. In addition, we choose to utilize PD’s in-built algorithms to estimate protein abundance differences based on the available peptide group data for the control experiment. If a protein abundance difference is greater than 2-fold and the P-value calculated by PD is less than 0.01, this was considered to be a significant protein abundance difference, in which case the value of (protein abundance-heat shock sample)/(protein abundance-Native) is used as a normalization factor for all peptides that map to that protein in the analysis of the LiP experiment. If either of those thresholds are not met (or if no quantification data is available for said protein, or if that protein was not identified in the control experiment), then no normalization is conducted for the peptides that map to that specific protein. The analyzer returns a file listing all the peptides that can be confidently quantified, and provides their effect-size, P-value, proteinase K site (if half-tryptic), and associated protein metadata. Similar to described above, the number of significant and all-or-nothing peptides are counted for each protein in condition 1 and 2. Proteins are only admitted into the comparison if two or more peptides are identified. To classify a protein as structurally perturbed (either during heat shock or recovery), it must have two or more peptides with significant abundance differences relative to non-heat-shocked controls.

### 7f. Dispersal Analysis

For heat stress granule dispersal assays, protein abundances in each of the immunoprecipitations were determined using the protein-level output from ProteomeDiscoverer’s Minora feature detector node. The abundances were normalized using the quantifications from BCA assays described in section 5d. Dispersal kinetics generally did not fit well to single-phase exponentials (although Pab1 itself did), hence dispersal curves are presented as datapoints rather than as fits. Fast proteins (those which are dispersed quickly) were defined as those that had a 50% reduction in their population relative to the heat-shocked sample, and then did not rise in population after. As a caveat to the second rule, we found that 20% fluctuations between datapoints was common and that the measurements were less accurate when the quantification was 0.1 or less; hence we allowed for increases in upto 20% between consecutive datapoints or if the quantification was below 0.1. Reaggregators were defined as those that had a large (2-fold) increases between any two consecutive time-points. This was discounted if the quantification was 0.1 or less.

### 7g. Orthology Analysis

For orthology analysis (i.e., comparing proteins that have refolded in *E. coli* vs *S. cerevisae*), orthologous pairs were determined from InParanoiDB 9^71, 72^. Using the list of proteins in each orthogroup (as provided by InParanoiDB 9), we combined the subset of proteins that were identified as orthologous with a match > 0.6 in organism 1 and organism 2. We compiled together the total number of peptides, the number of significant peptides, the number of and all-or-nothing peptides that were observed for each protein in organism 1 or organism 2. Orthologous pairs are only admitted into the analysis if 2 or more peptides were quantified in both organisms. Proteins are classified as refolding in both organisms, refolding only in organism 1, refolding only in organism 2, or nonrefolding in either organism. Orthologous pairs are discarded from the analysis if they are on the border; e.g., one significant peptide was found in organism 1, and two significant peptides were found in organism 2.

## 8. Bioinformatics and Metadata

### 8a. Bioinformatics

Saccharomyces Genome Database^73^ (SGD) was employed as the primary resource for metadata about yeast proteins, and its website was scraped with a script to obtain information about length, molecular weight, isoelectric point, median abundance, cellular components, complexes, and sequences for each protein. CYC2008^74^ was used to obtain information about cellular complexes to couple information from SGD, or to identify complexes not provided by SGD. Data from ref. 40 was used to obtain percentages of proteins found within a supernatant fraction upon heat-shock (pSup). pSup values for 42°C heat shock, 46°C heat shock, and 30°C control conditions were taken where available. The difference between pSup at 30°C – 42°C and 30°C – 46°C were calculated by subtraction. DomainMapper^27^ was used to obtain unique domain assignments utilizing domain definitions from the Evolutionary Classification of Domains^26^ (ECOD, http://http://prodata.swmed.edu/ecod/). E-values for domain matches, residue ranges, architectures, X-groups, T-groups, and F-groups were obtained for each protein. The number of domains were calculated as the number of architectures identified for each protein. The percent of the protein’s sequence found within an ECOD domain was calculated as proportion of residues identified as within a fold region by DomainMapper to the length of the protein. MetaPredict^25^ was used to identify the percent disorder, disordered ranges, and ordered ranges for each protein by feeding the sequence from SGD into MetaPredict using the commands (meta.percent_disorder or meta.predict_disorder_domains). The number of times a given protein engages the CCT chaperonin or the Ssb Hsp70 was taken from the dataset reported in ref. 24. Several “lists of stress granule proteins” exist for stress granule resident proteins, and the definitions do not fully overlap. Generally we apply the definition of pSup42°C < 0.4 from ref. 40, as the operating definition for a “heat stress granule” protein. Other proteomic investigations were undertaken by Jain et al.^42^, Cherkasov et al.^41^ and the semi-overlapping protein lists were compiled by Zhu et al.^43^. SGD also lists ‘cytoplasmic stress granule’ as a potential “cellular component” for each protein it indexes. We compiled all these definitions and include them as part of metadata, though we note that only the definition based on pSup has a notably high association with refoldability.

### 8b. Quantification and statistical analysis

All analyses of aggregation were conducted on independent refolding reactions from independent biological replicates (n = 3). Raw values shown for pelleting assay and significance by *t* test with Welch’s correction for unequal population variances. Standard target-decoy based approaches were used to filter protein identifications to an FDR < 1%, as implemented by Philosopher. All mass spectrometry experiments were conducted on three biological replicates used to generate three native samples and three independent refolding reactions from the same biological replicates. For heat shock perturbation studies, four biological replicates used to generate three native samples and four independent refolding reactions from the same biological quadruplicates. For each peptide group, abundance difference in refolded relative to native was judged by the *t* test with Welch’s correction for unequal population variances. Fisher’s method was used to combine P-values when there were multiple quantifiable features per peptide group. P-values less than 0.01 were used as a requirement to consider a region structurally distinct in the refolded form. Differences in means of distributions are assessed with the Mann-Whitney rank-sum test. To test whether particular categories are enriched with (or de-enriched with) (non)refoldable proteins, the chi square test or Fisher’s exact test is used.

## 9. Linker Length Analysis

Linker length analyses uses one of two distinct starting points. One starting point distinguishes between ‘ordered’ and ‘disordered’ regions within a protein, and this employed the artificial intelligence-based tools, MetaPredict^25^. Boundaries for ordered and disordered regions were calculated by calling the function meta.predict_disorder_domains.folded_domain_boundaries and meta.predict_disorder_domains.disordered_domain_boundaries. Another starting point distinguishes between residues assigned to structured domains and residues outside such boundaries, using the tool DomainMapper^27^. DomainMapper uses the ECOD system^26^ of Hidden Markov Model Profile-based domain definitions and parses the hits into a unique domain structure solution. The percent of the protein’s sequence found within an ECOD domain was calculated as the number of residues assigned to a domain by DomainMapper divided by the length of the protein. Both MetaPredict and DomainMapper were provided amino acid sequences from SGD and were used to determine ordered/disordered and in domain/linker residue ranges.

Depending on the figure, different analyses were conducted using these starting points.

### 9a. (Dis)Ordered vs. (Non)refoldable at the level of individual PK cut sites (Figure 2F, Extended Data Figure 6C)

Peptide-level quantifications were parsed to determine for each quantified peptide (which corresponds to a cut site) whether it falls within ordered or disordered regions (as classified by MetaPredict), and whether the site peptide was considered to be nonrefoldable (i.e., present at significantly different abundance in refolded samples compared to native references). Full-tryptic peptides were only considered if the entire peptide fell within an ordered region or disordered region. Half-tryptic peptides were considered ordered if the proteinase K cut site fell within an ordered region or disordered if in a disordered region. Fisher’s exact test was used to determine the significance of the resulting 2×2 contingency table.

### 9b. Refoldability vs. Linker Length for Two-domain proteins (Figure 2G, Extended Data Figure 6A)

Proteins with only two ECOD domains (based on DomainMapper) were isolated from the dataset. “Linker length” was calculated as the arithmetic difference between the terminal residue position of the first domain and the initial residue position of the subsequent domain. Protein-level refoldability for each protein was determined using the criteria stated above. The chi-square test was used to determine statistical significance for the relationship between refoldability of each two-domain protein and the length of its calculated linker length.

### 9c. Refoldability vs. Linker Length for Multidomain proteins (Figure 2H, Extended Data Figure 6B)

“Domain pairs” were identified as any set of two consecutive ECOD-identified domains within proteins with 2 or more domains, referred to as “multidomain” proteins. For each domain pair, the linker length was calculated as described above. Both the number of total and nonrefoldable sites were documented for both domains in the domain pair. A domain pair was considered to be refoldable if both domains were considered refoldable under the criteria described above, or nonrefoldable if one or both domains was found to be nonrefoldable. The chi-square test was used to determine statistical significance for the relationship between the fraction of domain-pairs found to be refoldable and the linker length intervening between the domain pair.

### 9d. (Non)Refoldability vs. Linker Length or Distance to Domain at the level of individual PK cut sites (Extended Data Figure 4D-E)

Peptide-level quantifications were parsed to determine for each quantified peptide (which corresponds to a cut site), if it fell in a linker region: (i) how many residues lie between it and the closest folded domain; (ii) how long the linker region it fell in was; (iii) whether the linker region, in the context of the protein, was leading (before any folded domain), trailing (after all folded domains), or interdomain; and (iv) whether the site peptide was considered to be nonrefoldable (i.e., present at significantly different abundance in refolded samples compared to native references). The length of an interdomain linker was calculated as described above. The length of a leading or trailing linker was calculated as the arithmetic difference between either the beginning of the protein and the first residue within a domain, or the terminal position of the last domain and the last position of the protein, respectively. For full-tryptic peptides, the smaller difference between the beginning residue of the tryptic peptide and the terminal residue of a preceding domain, if available, or terminal residue of the tryptic peptide and beginning residue of a subsequent domain, if available, was calculated. In this analysis, full-tryptic peptides were only counted if they were fully contained within a linker region. For half-tryptic peptides, the smaller difference between the cut site residue position and either terminal residue position of a preceding domain or beginning residue position of a subsequent domain was calculated. The chi-square test was used to determine statistical significance of the relationship between ranges of distances (or linker lengths) with refoldability. Sites that corresponded to leading/trailing linkers or interdomain linkers were considered separately.

### 9e. Average X-group Refoldability vs. Linker Length (Extended Data Figure 11A)

For each ECOD-identified X-group, the average linker length was calculated as the average length of all linkers connecting that X-group with another preceding or succeeding X-group for all proteins. Average refoldability for was calculated as the fraction of examples for a given X-group found to be refoldable based on the above refoldability criteria compared to the total number of representations for that X-group across all proteins. The number of representations for each X-group is shown on a logarithmic scale.

**Table S1:** Reagent and Resources.

| Reagent or Resource | Source | Identifier |
| --- | --- | --- |
| **Bacterial Strains** |  |  |
| *E. coli* (Strain K12) | New England BioLabs | ER2738 |
| *S. cerevisiae* (Strain BY4741) | ATCC | ATCC 4040002 |
| *S. cerevisiae* (Strain BY4741)<br>genotype: PAB1-Clover::KanMX | Drummond Lab | N/A |
| **Chemicals and Reagents** |  |  |
| Agar | Fisher | BP1423-2 |
| Ammonium Bicarbonate | Acros Organics | 393210050 |
| D-(+)-Raffinose pentahydrate | Sigma-Aldrich | R7630-100g |
| D-Glucose (Dextrose) | Fisher | D16-500 |
| Dithiothreitol (DTT) | Sigma-Aldrich | D06632 |
| DMSO | Sigma-Aldrich | D8418 |
| EDTA | Sigma-Aldrich | 60-00-4 |
| Galactose | Fisher | 59-23-4 |
| Glycerol | Fisher | G33-1 |
| Guanidium Chloride (GdmCl) | Acros Organics | 50-01-1 |
| Heparin | Sigma-Aldrich | H3393 |
| Hydrochloric Acid (HCl) | Fisher Scientific | A142-212 |
| Iodoacetamide (IAA) | Acros organics | 122271000 |
| KCl | Fisher | BP366-1 |
| L-Leucine | Fisher | BP384-100 |
| L-Tryptophan | Acros Organics | 140590250 |
| Lithium Acetate | Sigma-Aldrich | 517992 |
| Magnesium Chloride | Fisher | BP214-500 |
| NP-40 Surfact-Amps Detergent | Thermo Scientific | 85124 |
| PEG (3350 MW) | Sigma Aldrich | 202444 |
| Peptone | Gibco | 211677 |
| PMSF | ThermoFisher | 36978 |
| Potassium Acetate | Sigma Aldrich | P1190 |
| SC Complete-Ura-Glucose Powder | Sunrise Science | 1461-500 |
| SD-Leu-Trp-Ura-Glucose | Sunrise Science | 1805-500 |
| Sodium Chloride | Fisher | BP358-212 |
| Tris-Base | Sigma-Aldrich | RD008 |
| Tris-HCl | Sigma | RD009 |
| Tryptone | Fisher | BP1421-2 |
| Uracil | Alfa Aesar | A15570 |
| Urea | Fisher | U17-212 |
| Yeast Extract | Fisher | BP1422-500 |
| **Components for Western Blot** |  |  |
| iBlot 2 Transfer Stacks, PVDF | Invitrogen | IB24001 |
| Nestle Carnation Instant Nonfat Dry Milk | Nestle | N/A |
| Novex Tris-Glycine Mini Protein Gels, 4-12%, 1.0 mm | ThermoFisher | XP04122BOX |
| PageRuler Prestained Protein Ladder | ThermoFisher | 26619 |
| SuperSignal West Femto Maximum Sensitivity Substrate | ThermoFisher | 34095 |
| TBS (10X), pH 7.4 | Quality Biological | 351-086-131 |
| TRIS-buffered saline (TBS, 10X) pH 8.0 | ThermoFisher | J62780.K2 |
| Tris/Glycine/SDS Running Buffer | Bio-Rad | 1610732 |
| Tween 20 | Sigma Aldrich | P1379 |
| **Components to Make Protease Inhibitor Cocktail** |  |  |
| AEBSF.HCl | Thermo Scientific | 78431 |
| Aprotinin from bovine lung | Sigma Aldrich | 10820 |
| Bestatin | Thermo Scientific | 78433 |
| E-64 | Thermo Scientific | 78434 |
| Leupeptin | Sigma Aldrich | EI8 |
| Pepstatin A | Thermo Scientific | 78436 |
| **Recombinant Proteins and Enzymes** |  |  |
| 6x-His Tag Monoclonal Antibody (HIS.H8) | Invitrogen | MA1-21315 |
| DNase I from Bovine Pancreas | Sigma Aldrich | 31136 |
| DpnI | New England BioLabs | R0176S |
| Goat anti-Mouse IgG (H+L) Secondary Antibody, HRP | Invitrogen | 31430 |
| Proteinase K | Fisher Scientific | BP1700 |
| Q5 High-Fidelity 2X Master Mix | New England BioLabs | M0492S |
| Trypsin-ultra, Mass Spectrometry Grade | New England BioLabs | P8101S |
| UltraPure Salmon Sperm DNA | Invitrogen | 15632011 |
| **Commercial Assays** |  |  |
| BCA Assay (Pierce) | Thermo Scientific | 23227 |
| Rapid Gold BCA Assay (Pierce) | Thermo Scientific | A53225 |
| **LC-MS Reagents** |  |  |
| Acetonitrile (HPLC Grade) | Sigma-Aldrich | 34851 |
| Formic Acid (Optima LC/MS Grade) | Fisher Scientific | A117-50 |
| Trifluoroacetic acid (TFA) (HPLC Grade) | Alfa Aesar | 44630 |
| Water (Optima LC-MS Grade) | Fisher Scientific | W6500 |
| **Other** |  |  |
| 3 mL *Konical Tubes* | Beckman Coulter | N/A |
| Anti-FLAG M2 Magnetic Beads | Sigma Aldrich | M8823 |
| Greiner Bio-One Black μClear Bottom 96-well Polystyrene Microplates | Fisher Scientific | 07-000-088 |
| Mass Spec Vials | Thermo Scientific | N/A |
| Nunc Lab-Tek II Chambered Coverglass (8 Well) | Thermo Scientific | 155409 |
| Sep Pak C18 1 cc 50 mg Resin | Waters | 186000308 |
| Viva-Spin 15R 2K MWCO Hydrosart | Sartorius | VS15RH92 |
| **Instrumentation** |  |  |
| AN60 Ti Rotor | Beckman Coulter | N/A |
| Eppendorf Centrifuge (5910R) | Eppendorf | N/A |
| iBlot 2 Gel Transfer Device | Thermo Scientific | IB21001 |
| pH Meter | Mettler-Toledo | N/A |
| Plate Reader (ID3) | Molecular Devices | N/A |
| Plate Reader (Spark) | Tecan | N/A |
| Q-Exactive HF-X Orbitrap Mass Spectrometer | Thermo Scientific | N/A |
| SPEX™ Sample Prep Dual Freezer/Mill | SPEX | N/A |
| SW55 Ti Rotor | Beckman Coulter | N/A |
| UltiMate3000 UHPLC | Thermo Scientific | N/A |
| Ultracentrifuge (Optima XL-A) | Beckman Coulter | N/A |
| **Software and Algorithms** |  |  |
| AnalyzerV18 | In House | https://github.com/FriedLabJHU/ |
|  |  | Refoldability-Tools |
| FIJI | Schindelin et al., 2012 | N/A |
| MSFragger Node in PD | Nesvizhskii Lab | N/A |
| Proteome Discoverer v 2.4 (PD) | Thermo Scientific | N/A |
| VisiView | Visitron Systems | N/A |
| XCalibur | Thermo Scientific | N/A |
| **Databases** |  |  |
| CYC2008 | Wodak Lab | https://wodaklab.org/cyc2008/ |
| Evolutionary Classification of Protein Domains | Grishin Lab | http://prodata.swmed.edu/ecod/ |
| SGD | Stanford University | https://www.yeastgenome.org/ |

**Table S2:**
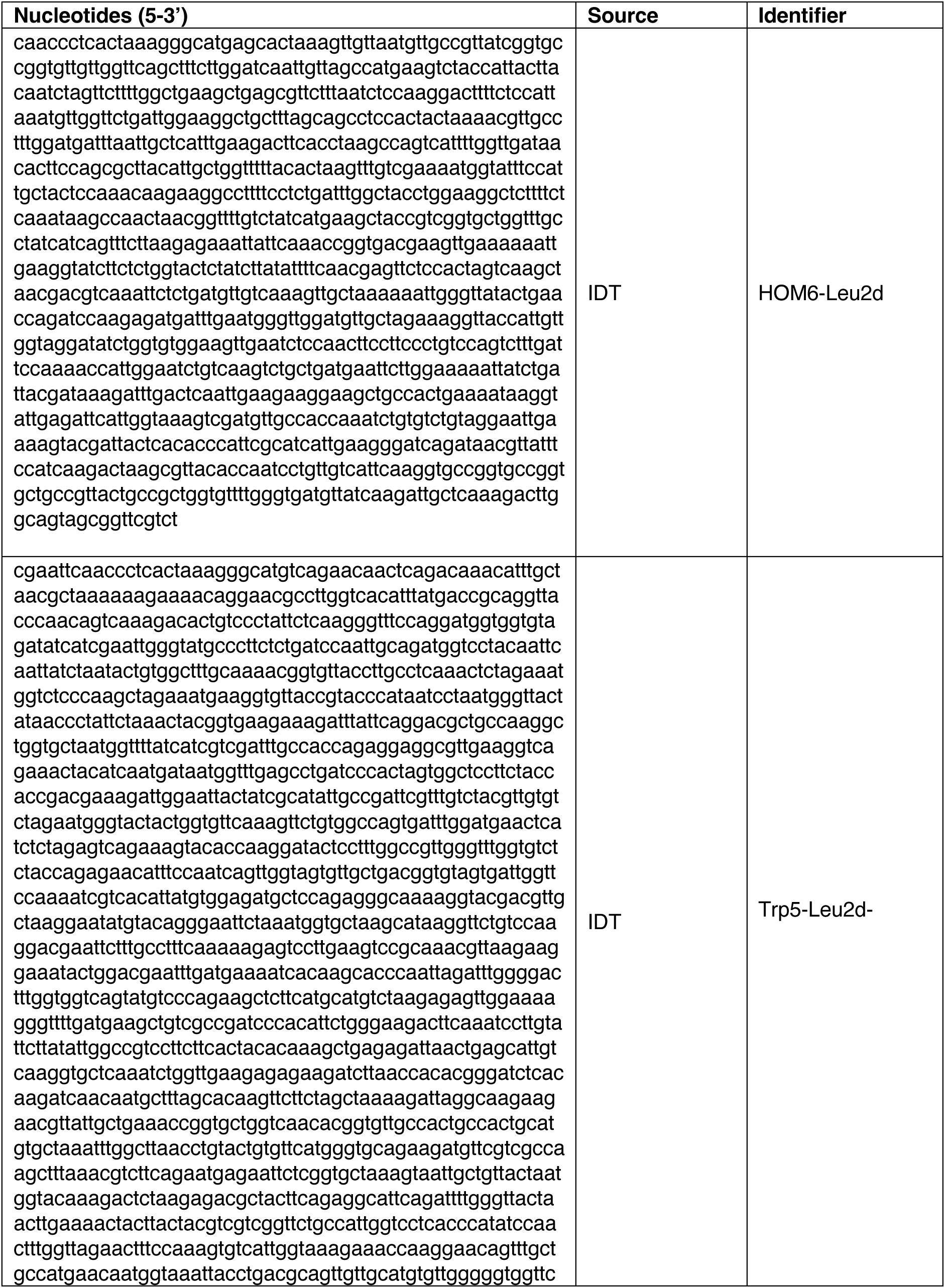
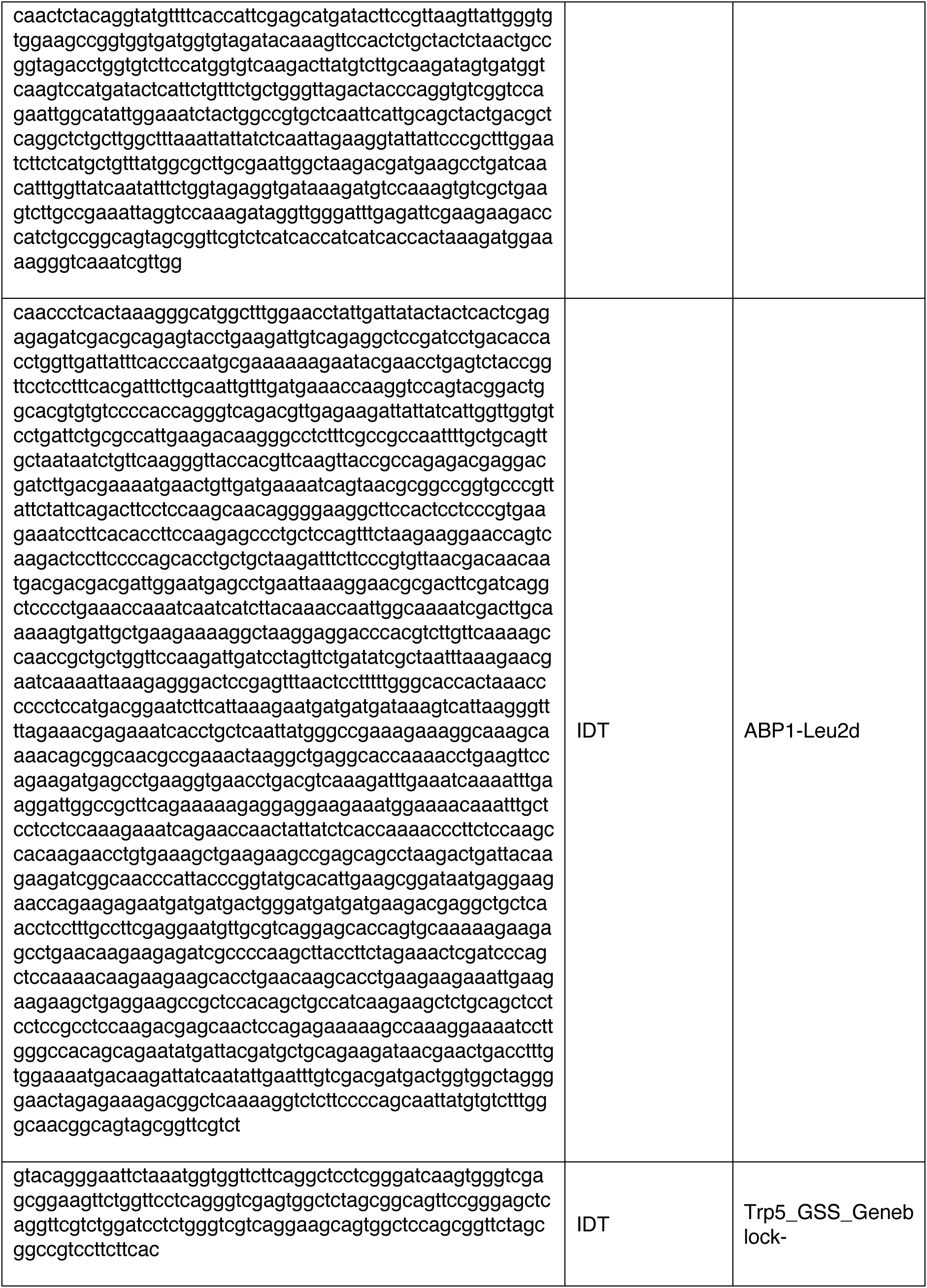
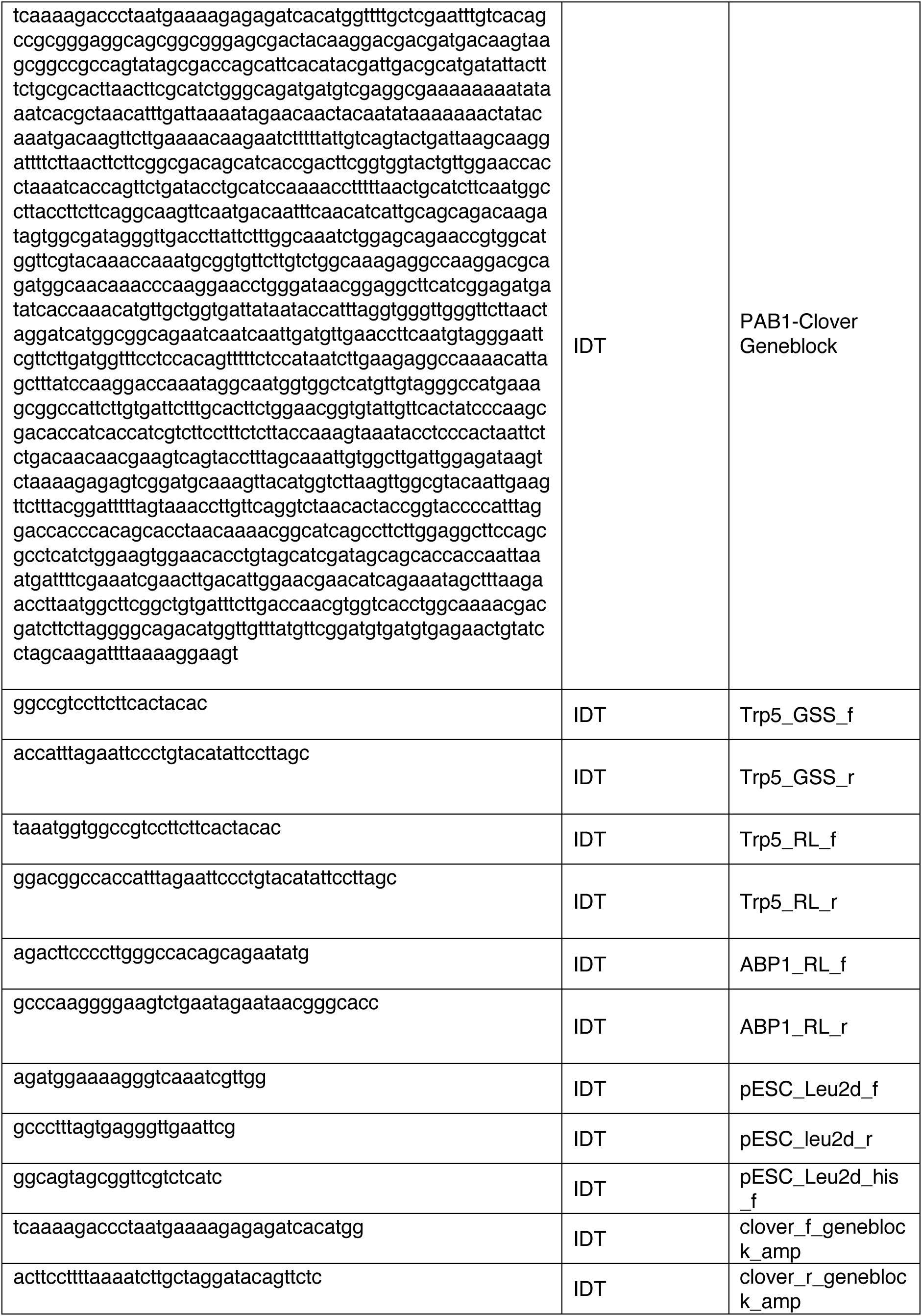
Geneblocks and Oligonucleotides.

**Table S3:** Resource Availability Table.

| Resource | Source | Location |
| --- | --- | --- |
| Domain Summary Files | In House | Supplemental Data Files (this paper) |
| Protein Metadata Files | In House | Supplemental Data Files (this paper) |
| PD v2.4 LFQ Output Files (.pdresult) | In House | PRIDE Accession XXX (doi: XXX) |
| Peptide Quantifications (*_out.txt) | In House | Zenodo (doi: XXX) |
| Protein Summary Data | In House | Supplemental Data Files (this paper) |
| Raw Mass Spectra Files (.raw) | In House | PRIDE Accession XXX (doi: XXX) |

**Table S4:** Proteome Discoverer Peptide Identification Node Parameters.

|  | <u>MSFragger</u> |
| --- | --- |
| **Peak Matching and Outputting** |  |
| Precursor Mass Tolerance | 10 ppm |
| Fragment Mass Tolerance | 20 ppm |
| Use Average Precursor Mass | N/A |
| Use Average Fragment Mass | N/A |
| Isotope Error | 0/1/2 |
| Localized Delta Mass | 0 |
| Delta Mass Exclude Ranges | (-1.5,3,5) |
| Max. Precursor Charge | 4 |
| **Digestion** |  |
| Enzyme | Trypsin (semi) |
| Min. Peptide Length | 7 |
| Max. Peptide Length | 50 |
| Max. Missed Cleavage Sites | 2 |
| Min. Peptide Mass | 500 |
| Max. Peptide Mass | 5000 |
| Maximum Charge State for Theoretical Fragments to Match | 2 |
| Clip n Term M | TRUE |
| **Spectrum Matching** |  |
| Use of Neutral Loss a Ions | N/A |
| Use of Neutral Loss b Ions | N/A |
| Use of Neutral Loss y Ions | N/A |
| Use Ranking Ions | N/A |
| Weight of a Ions | N/A |
| Weight of a Ions | N/A |
| Weight of a Ions | N/A |
| Weight of a Ions | N/A |
| Weight of a Ions | N/A |
| Weight of a Ions | N/A |
| **Spectral Processing** |  |
| Min. Peaks | 15 |
| Min. Fragments Modeling | 2 |
| Min. Ratio | 0.01 |
| Use Top N Peaks | 150 |
| Min. Matched Fragments | 3 |
| Min. Clear m/z Range | 0 |
| Max. Clear m/z Range | 0 |
| Mass Calibration | On and find optimal parameters |
| **Modifications** |  |
| Max. Equal Modifications Per Peptide | N/A |
| Multiple Variable Mods on Residue | TRUE |
| Max. Variable Mods per Peptide | 3 |
| Max. Variable Mods Combinations | 5000 |
| Mass Offsets | 0 |
| **Dynamic Modifications** |  |
| Oxidation | +15.995 Da (M) |
| Acetyl | +42.011 Da (Any N-Terminus) |
| **Static Modifications** |  |
| Carbamidomethyl | +57.021 Da (C) |

## Data Availability

All raw proteomic data and label free quantification (LFQ) outputs (corresponding to global refolding assays in yeast [three replicates], global refolding assays in *N. crassa*, LiP-MS assays in response to heat shock, and heat street granule dispersal assays) are available on PRIDE under the accession code XXXXX. LFQ outputs were processed by home-made scripts to generate peptide-level quantifications; these out files are available on Zenodo under the accession code XXXX. The peptide-level quantifications were further analyzed to generate summary files that represent refoldability (or structural perturbation) at the protein-level; these are available as supplemental tables to this paper. Several additional analyses were performed to analyze domains, linker lengths, orthology, etc.; these are available as supplemental tables to this paper.

## Acknowledgments

We thank Dr. Philip Mortimer for maintaining the mass spectrometry core facility at JHU chemistry. We thank Edgar Manriquez-Sandoval for performing DomainMapper analyses. We thank Dr. John Kim for providing advice on stress granule isolation procedures. We acknowledge Jiaxin Li and Dr. Kyle Cunningham for providing advice on yeast transformation and culturing. We are deeply indebted to Drs. Allan Drummond and Jared Bard for introducing us to heat stress response in yeast, and for providing us with a *S. cerevisiae* strain containing Pab1 C-terminally fused with Clover. We thank Drs. Sua Myong, Taekjip Ha, John Kim, and Geraldine Seydoux for insightful conversations and commenting on the manuscript.

H.E.T. and H.W. acknowledge support from the Chemistry-Biology Interface NIH training grant program (T32-GM080189 and T32-GM149382). H.E.T. acknowledges support from the age-related neuroscience NIH training grant (T32-AG027668). A.E. acknowledges support from the Cellular, Molecular, and Developmental Biology NIH training grant program (T32-GM141804). S.D.F. acknowledges grants from the NIH Director’s New Innovator Award (DP2-GM140926), from the NSF Division of Molecular and Cellular Biology (MCB2045844), from the Human Frontiers in Science Program (RGY0074/2019) and from the RCSA foundation (Award ID 28413).

## Author Information

These authors contributed equally: Philip To, Atharva Bhagwat

## Authors and Affiliations

Department of Chemistry, Johns Hopkins University, Baltimore, MD, USA

-Philip To, Atharva Bhagwat, Haley E. Tarbox, Zanya Jamieson, Stephen D. Fried

Department of Biology, Johns Hopkins University, Baltimore, MD, USA

-Ayse Ecer, Tatjana Trcek

T. C. Jenkins Department of Biophysics, Johns Hopkins University, Baltimore, MD, USA-Stephen D. Fried

Department of Biophysics and Biophysical Chemistry, Johns Hopkins University School of Medicine, Baltimore, MD 21205, USA

-Hannah Wendorff

## Author Contributions

S.D.F. designed the study. P.T. and A.M.B. optimized and performed refolding experiments in yeast. H.E.T. and Z.J. optimized and performed refolding studies in *N. crassa*. P.T. and H.W. performed aggregation assays and related Western blot experiments. P.T. and A.M.B. performed proteomics experiments on yeast subjected to heat stress. P.T. and A.E. performed stress granule isolation, imaging, and fluorescence studies. T.T. advised imaging experiments and related data analysis. P.T. and S.D.F. performed stress granule dispersal assays. P.T., A.M.B., and S.D.F. curated metadata and performed bioinformatic analyses. P.T., A.M.B., and S.D.F. analyzed and visualized data from proteomics experiments. P.T. analyzed and visualized data from aggregation assays, fluorescence, and imaging. P.T. and S.D.F. generated figures. S.D.F. wrote the first draft of the manuscript. P.T. and S.D.F. revised the manuscript with feedback from all authors. P.T. and A.M.B. prepared supplementary data tables in a disseminatable form and collected proteomic data for accession to PRIDE. S.D.F. mentored trainees and secured funding for the project.

**The authors have no competing interests to declare.**

## Extended Data Figures

**Extended Data Figure 1.**
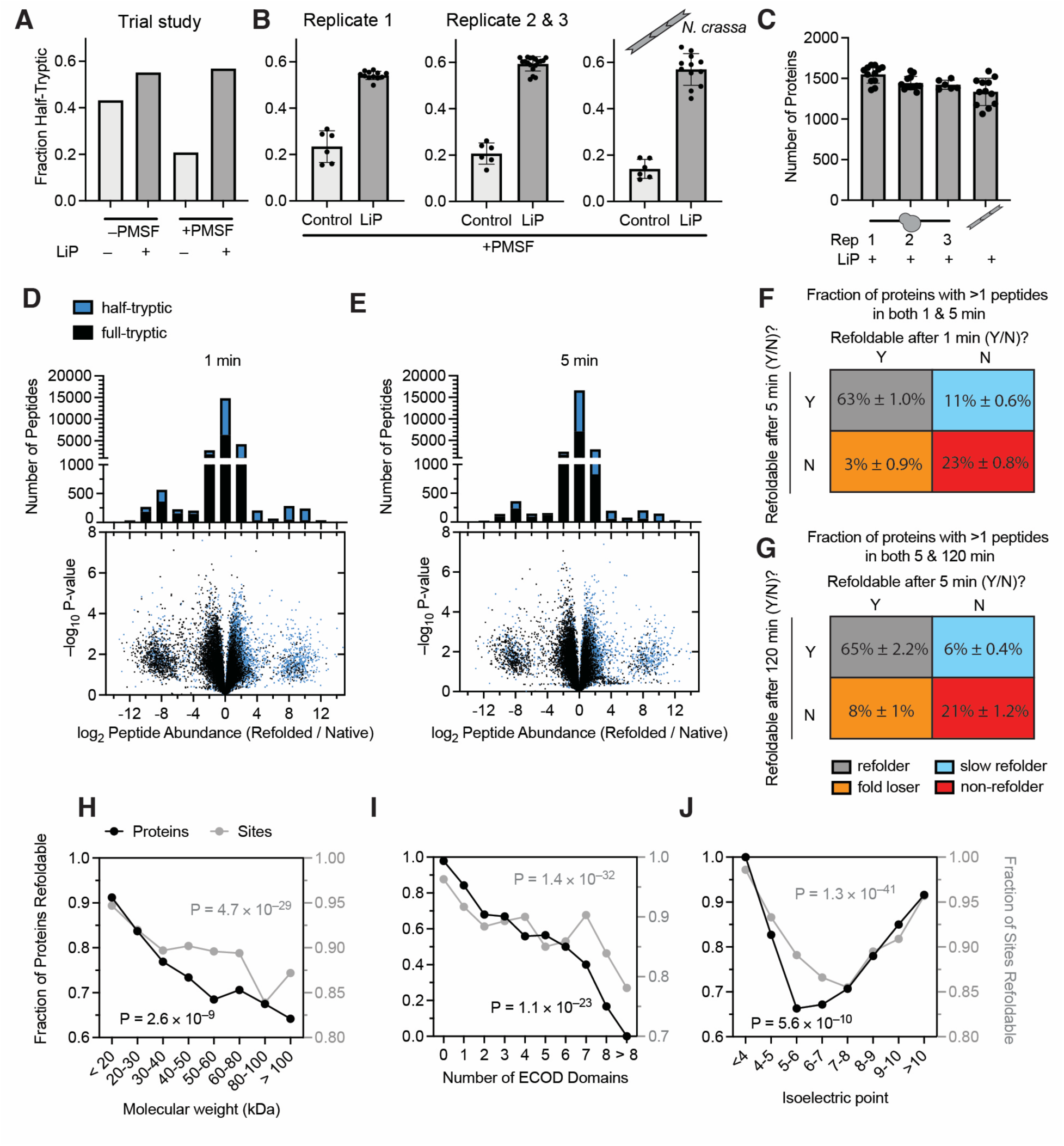
Development, Evaluation, and Robustness Analysis of Global Refolding Assays in Yeast. (A) Effect of adding the irreversible serine protease inhibitor phenylmethylsulonyl fluoride (PMSF) to lysis buffer during global refolding limited-proteolysis (LiP) mass spectrometry assays, measured by the fraction of identified peptides that are half tryptic in a semi-specific search (a quality control metric). (B) Half trypticity values for the four production refolding experiments reported in this study, three separate replicates of yeast refolding and one for *N. crassa* refolding. Control implies no limited proteolysis step was performed. Replicates two and three were performed concurrently. (D, E) Volcano plots showing all peptides quantified in a yeast global refolding LiP-MS experiment, along with histograms displaying the number of peptides quantified with refolded/native ratios in various ranges. Black dots are full-tryptic peptides (representing sites less susceptible following refolding), blue dots are half-tryptic peptides (sites more susceptible following refolding). Panel D is after 1 min of refolding, E is after 5 min of refolding. (F, G) Contingency tables summarizing refolding kinetics by showing the percent of proteins in one of four kinetic categories. Percentages are over three replicates of yeast refolding experiment. Panel F compares 1 min to 5 min refolding times; panel G compares 1 min to 2 h refolding times. (H, I, J) Trends between refolding propensity and (H) molecular weight; (I) isoelectric point; and (J) number of domains. Black dots represent percent of proteins in that category which refold (as in Figure 1H-J). Gray dots represent the fraction of sites associated with proteins in that category which are refoldable. P-values are from chi-square test, and come from replicate 1.

**Extended Data Figure 2.**
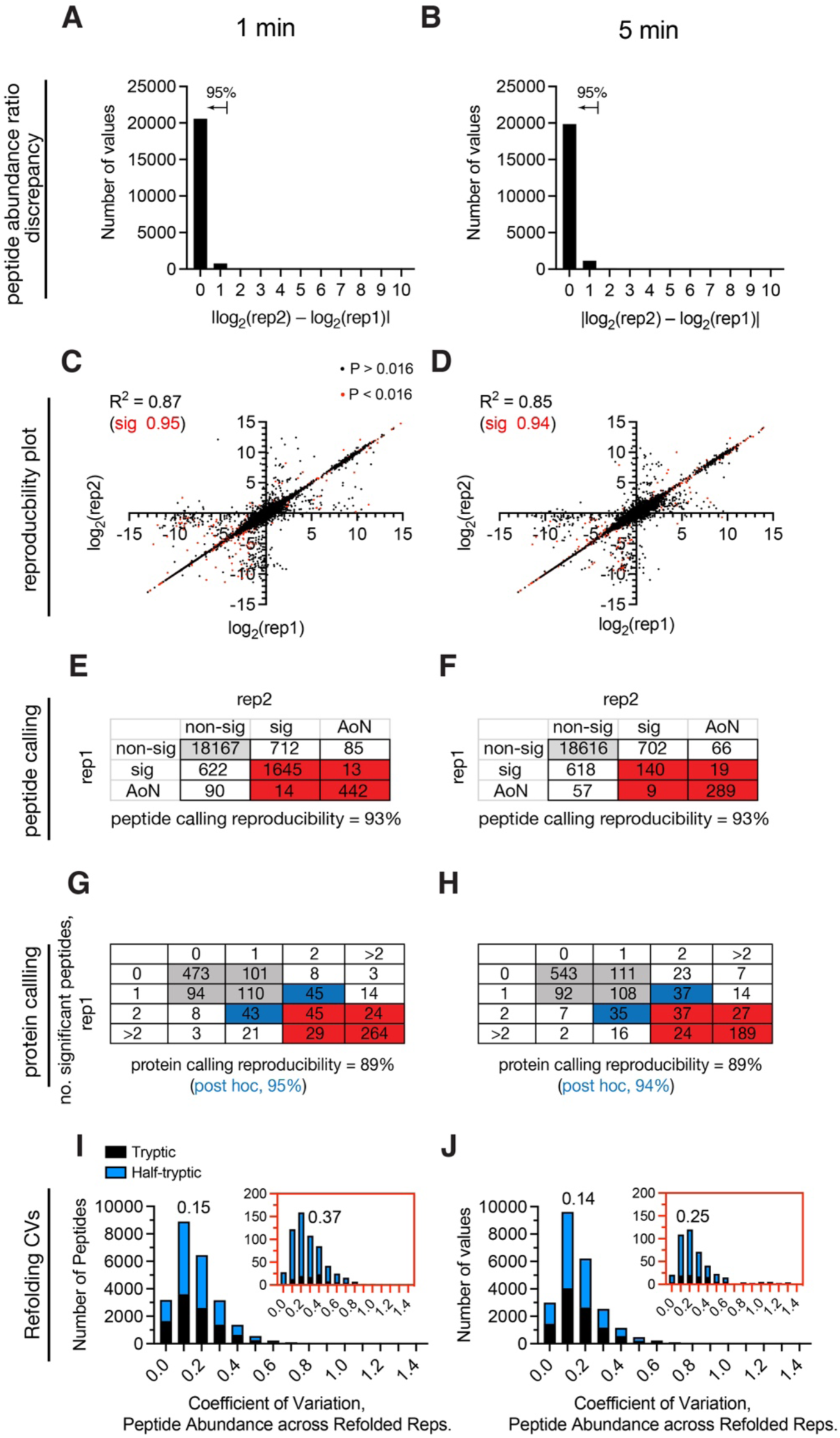
Reproducibility analysis. **(A-B)** Histograms showing the peptide quantification discrepancies between two replicates of the experiment in which proteins were refolded for **(A)** 1 min or **(B)** 5 min. Note that each of these replicates of the experiment involved three separate biological replicates of native and refolded. Peptides that were identified in both experiments were collected and the refolded/native ratio in each replicate was compared to each other. Histograms show the absolute value of the difference of the log2 quantifications. **(C-D)** Scatter plots showing the relationship between the peptide log2(refolded/native) quantification in one replicate versus its value in the other replicate for two replicates of the experiment in which proteins were refolded for **(C)** 1 min or (**D**) 5 min. Points in red were considered significant (P < 0.016 by Welch’s t-test) in both experiments. The coefficients of determination (*R*^2^) are given first for all points in common (black), and then for the subset of points that were considered significant in both replicates of the experiment (red). **(E-F)** Calling reproducibility of peptides (classified as either non-significant, significant, or all-or-nothing (AoN)) between two replicates of the experiment in which proteins were refolded for **(E)** 1 min or **(F)** 5 min. **(G-H)** Calling reproducibility of proteins between two replicates of the experiment in which proteins were refolded for **(G)** 1 min or **(H)** 5 min. Rows correspond to the number of peptides that were significantly different between native and refolded samples in the first replicate of the experiment, and columns correspond to the number of peptides that were significantly different in the duplication. Numbers in the table correspond to the number of proteins with that many significant peptides in each replicate. Gray cells correspond to proteins that would be called refoldable in both iterations. Red cells correspond to proteins that would be called nonrefoldable in both iterations. Cells in white would have been called differently, resulting in reproducibility of 89%. In all comparisons, we exclude proteins that only differ by one significant peptide at the cut-off, shown as blue cells. With these proteins removed *post hoc*, reproducibility increases to 94–95%. **(I-J)** Histograms of the coefficients of variation (CV) for the peptide abundances in refolded samples, from 3 independent refolding reactions, in which proteins were refolded **(I)** 1 min or **(J)** 5 min. Insets in red correspond to the CV histograms for the peptides detected only in the refolded samples. Numbers represent medians of distributions.

**Extended Data Figure 3.**
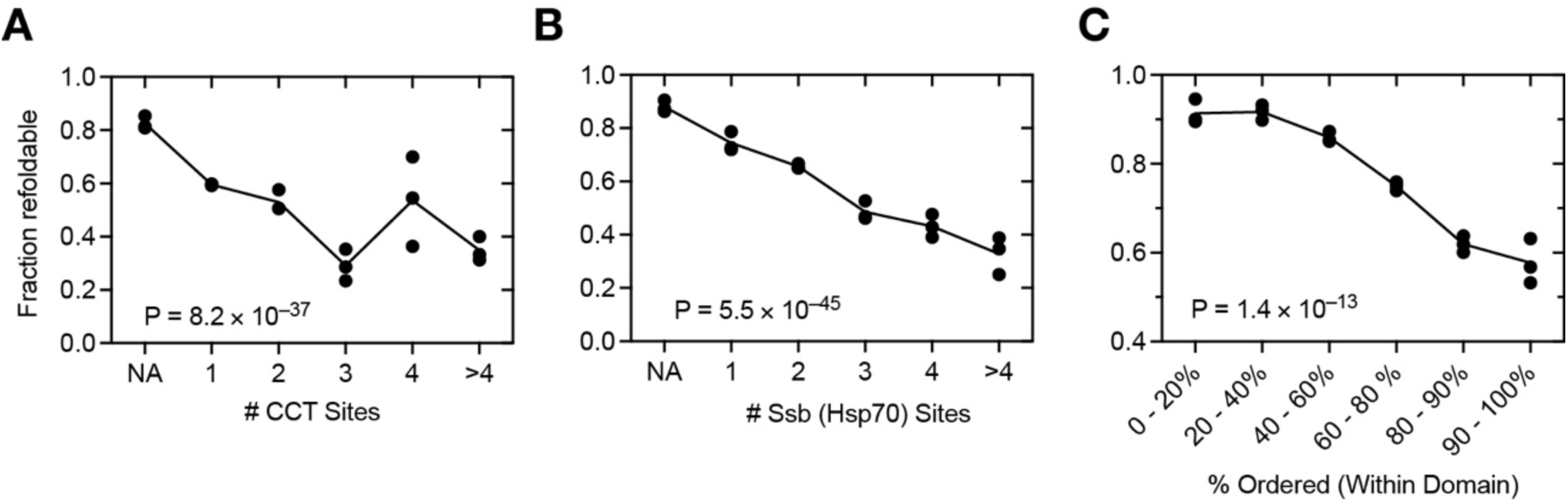
Refoldability Anti-Correlates with Chaperone Usage During Primary Biogenesis. Fractional refoldability for proteins divided by (A) the number of times the nascent protein engages TRiC/CCT chaperonin, (B) the number of times the nascent protein engages Ssb1/2p (Hsp70) chaperone, and (C) the percentage of the residues within the protein associated with an ECOD domain. Data from ref. 24 (Ribo-Seq assays) for panels A, B. Three separate performances of the assay were performed for yeast, each time with three biological replicates. P-values are from chi-square test; for yeast all P-values come from replicate 1.

**Extended Data Figure 4.**
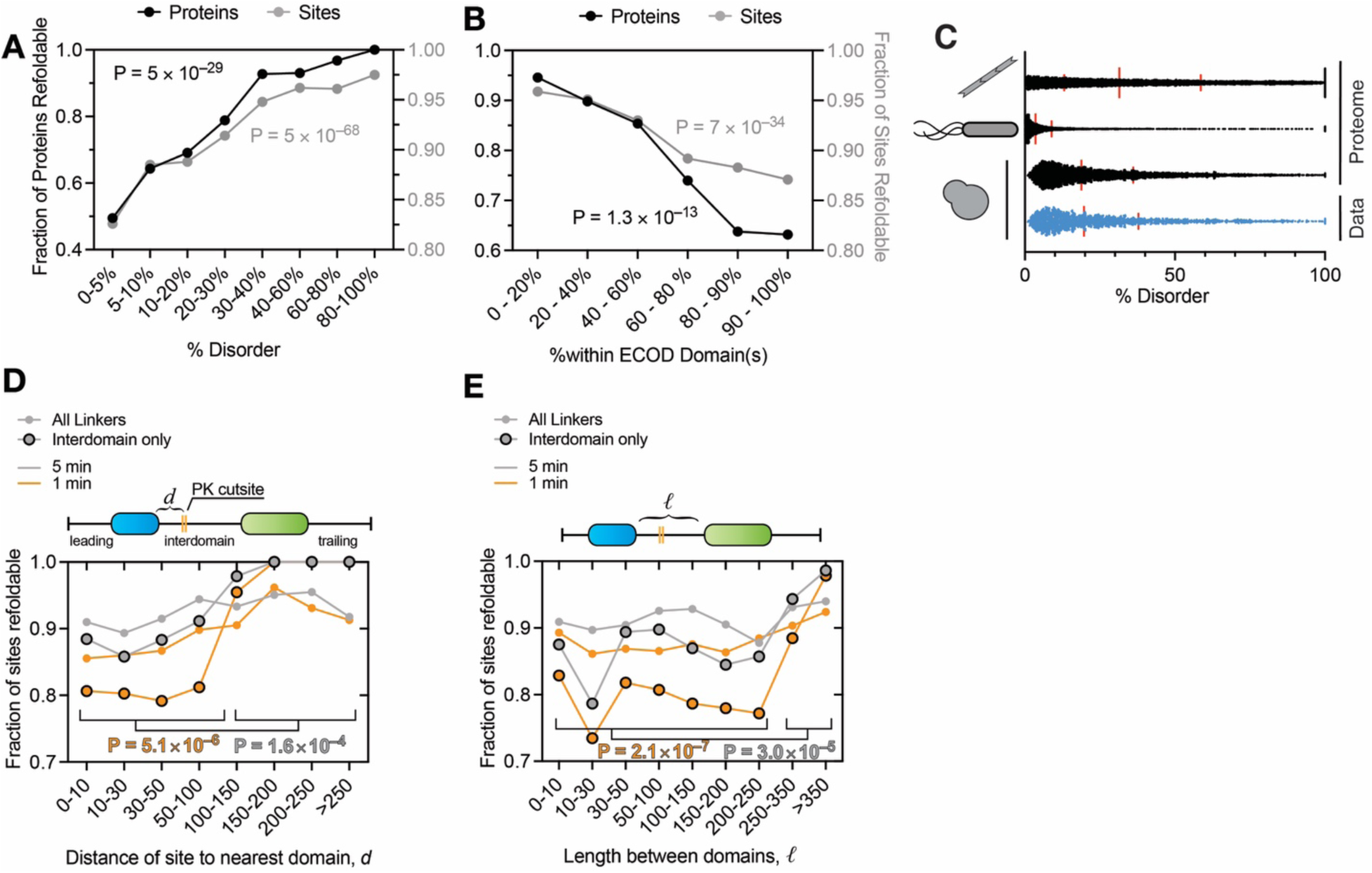
Robustness Analysis of Refoldability Trends Related to Disorder. (A, B) Trends between refolding propensity and (A) % disorder as calculated by metapredict and (B) % of amino acid residues within ECOD domains. Black dots represent percent of proteins in that category which refold. Gray dots represent the fraction of sites associated with proteins in that category which are refoldable. P-values are from chi-square test, and come from replicate 1. (C) Distribution of the %disorder of the *N. crassa*, *E. coli*, and yeast protomes. Red lines represent 1^st^ quartile, median, and 3^rd^ quartile. Median %disorders are 31.45%, 3.53%, 18.9%. Shown in blue is the %disorder distribution on the yeast proteins for which refoldability data is available (median = 19.7%). (D) Fraction of sites outside domains (in linker regions) that are refoldable (i.e., same susceptibility to PK after unfolding/refolding) grouped by the distance of the site to the closest domain. Orange, refoldability after 1 min; gray, refoldability after 5 min. Open circles, all linker regions included; bordered circles, only considering linkers between domains. P-values calculated by pooling together interdomain sites “close” to domains (0–100 residues) or “far” from domains (>101 residues) and evaluating with Fisher’s exact test. (E) Fraction of sites outside domains that are refoldable grouped by the length of the interdomain segment. Colours and borders have same meaning as panel D.

**Extended Data Figure 5.**
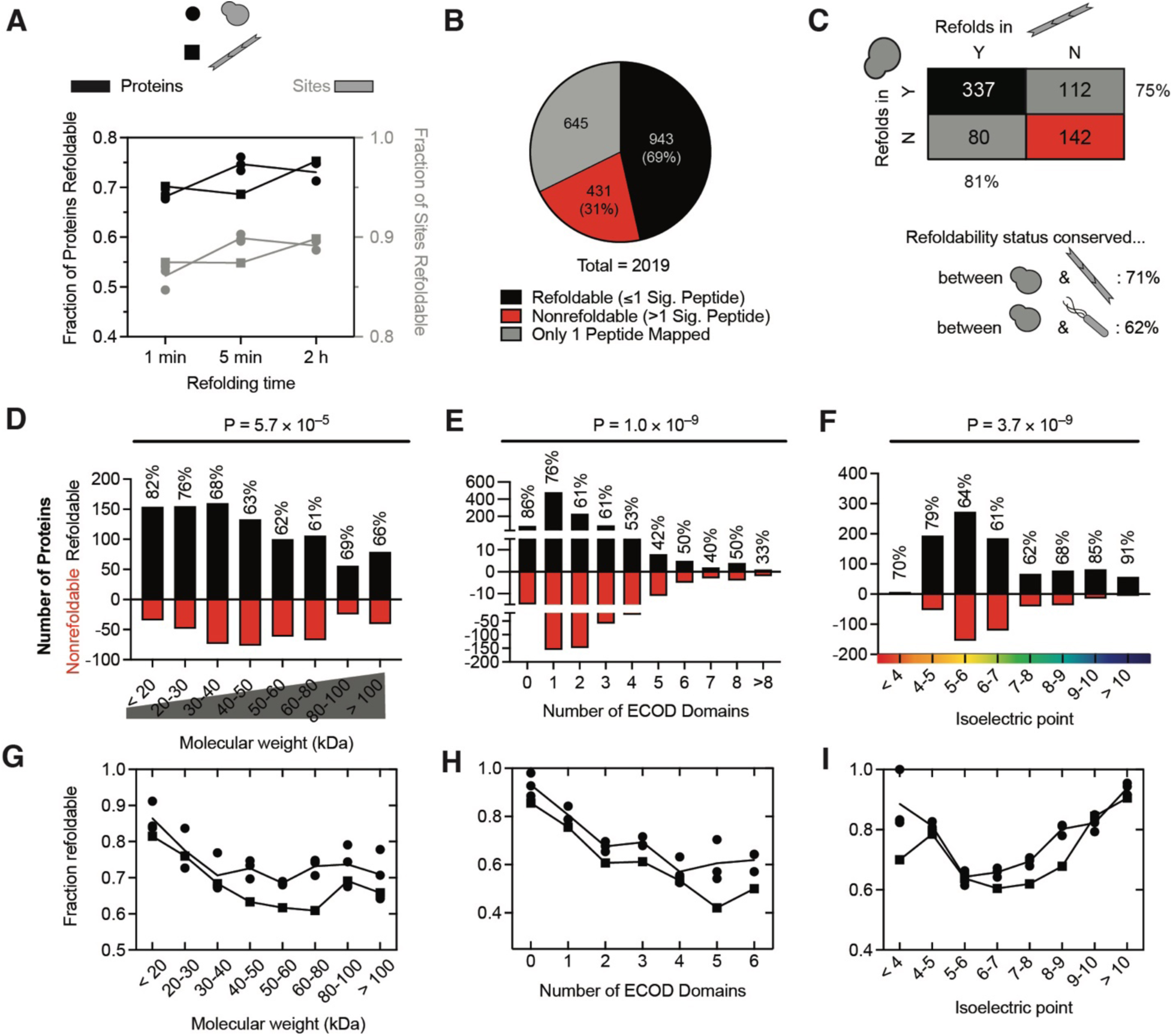
The *Neurospora crassa* Proteome’s Refoldability Profile Mimics Yeast’s. (A) Fraction of sites (i.e., mapped peptides; gray) and proteins (black) that are refoldable in yeast (circles) or *N. crassa* (squares), based on if their susceptibility to proteinase K is the same or different after unfolding/refolding compared to a native reference. Refolding reactions were analyzed at three times following dilution from denaturant. (B) Pie chart showing (non)refolding protein counts after 5 min. (C) Contingency table showing the number of *N. crassa*/yeast orthologous pairs with a given refoldability status in either organism (5 min refolding time). Shown below is the refoldability conservation for both species pairs. (D-F) The number of refoldable (black) and nonrefoldable (red) proteins in *N. crassa* divided by (D) molecular weight, (E) number of folded domains, and (F) isoelectric point at the 5 min refolding time. (G-I) Fractional refoldability for proteins in the categories shown in D-F, for *N. crassa* (squares) and yeast (circles). P-values are from chi-square test.

**Extended Data Figure 6.**
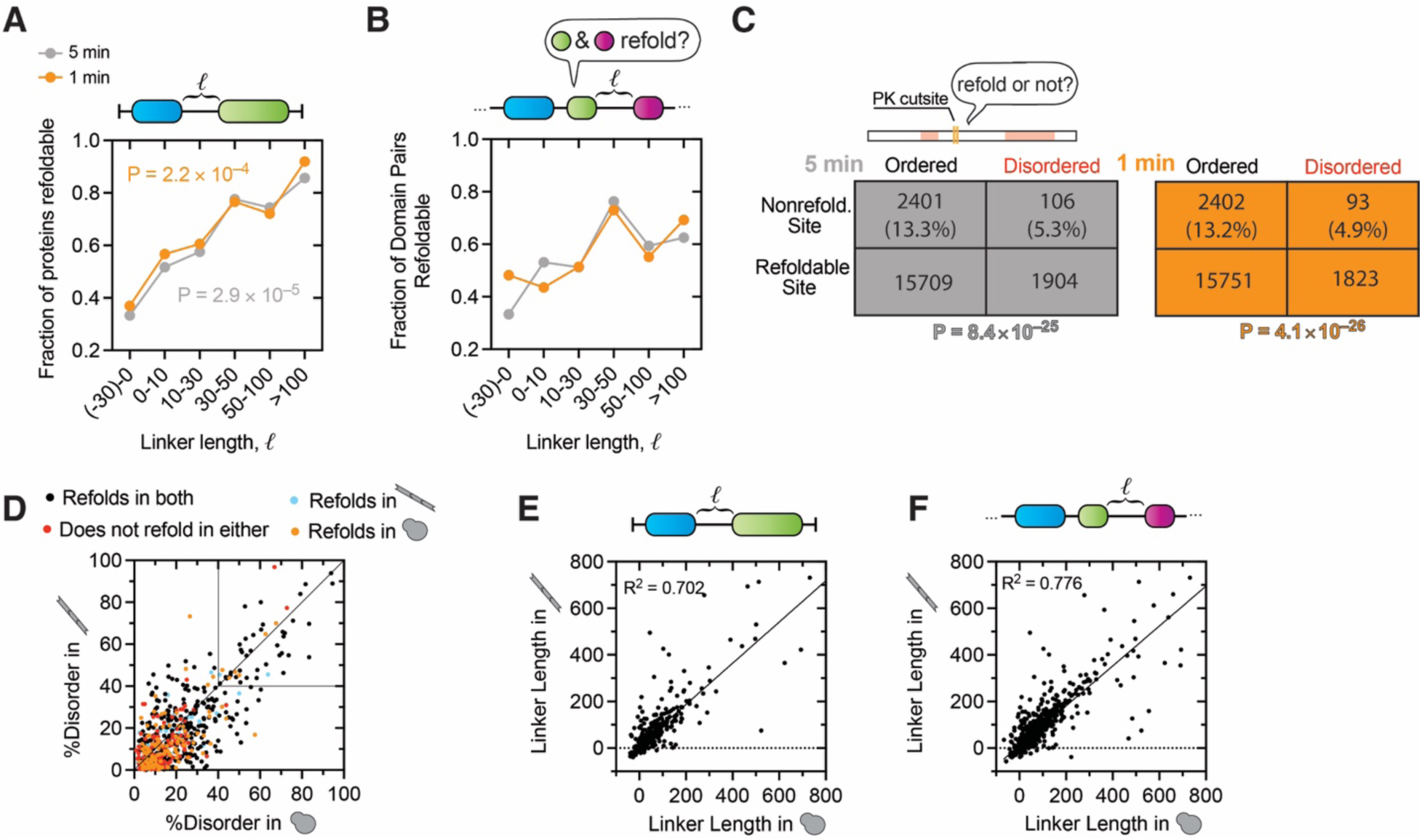
Disorder Promotes Refoldability in *Neurospora crassa* in a Manner Similar to Yeast. (A) Fractional refoldability of two-domain proteins in *N. crassa* as a function of linker length between domains, calculated with DomainMapper. Orange, 1 min refolding time; Gray, 5 min refolding time. P-values calculated by chi-square test. (B) Fractional refoldability of consecutive domain pairs within all *N. crassa* proteins with ≥2 domains as a function of linker length between pair of domains. (C) Contingency tables showing the number of assessed sites that are refoldable or not in terms of whether they fall in an ordered or disordered region. Sites in disordered regions are substantially (and according to Fisher’s exact test, significantly, P-values given) more likely to refold. Colour denotes refolding time (5 min, gray; 1 min, orange). (D) Plot of each *N. crassa*/yeast orthologous pair with the %disorder of the yeast orthologue plotted on *x*-axis and %disorder of the *N. crassa* orthologue plotted on *y*-axis. Orthologous pair coloured based on whether they refold in both organisms (black), in neither (red), only in yeast (orange) or only in *N. crassa* (blue). Refoldable yeast proteins whose *N. crassa* orthologues do not refold tend to have more disorder. (E) Similar to panel D, except only two-domain proteins are included and linker length between the two domains in the two orthologous proteins are plotted. Coefficient of determination from a least-square regression line given. (F) Similar to panel E, except linker lengths are plotted for all domain-pairs in all proteins with ≥2 domains.

**Extended Data Figure 7.**
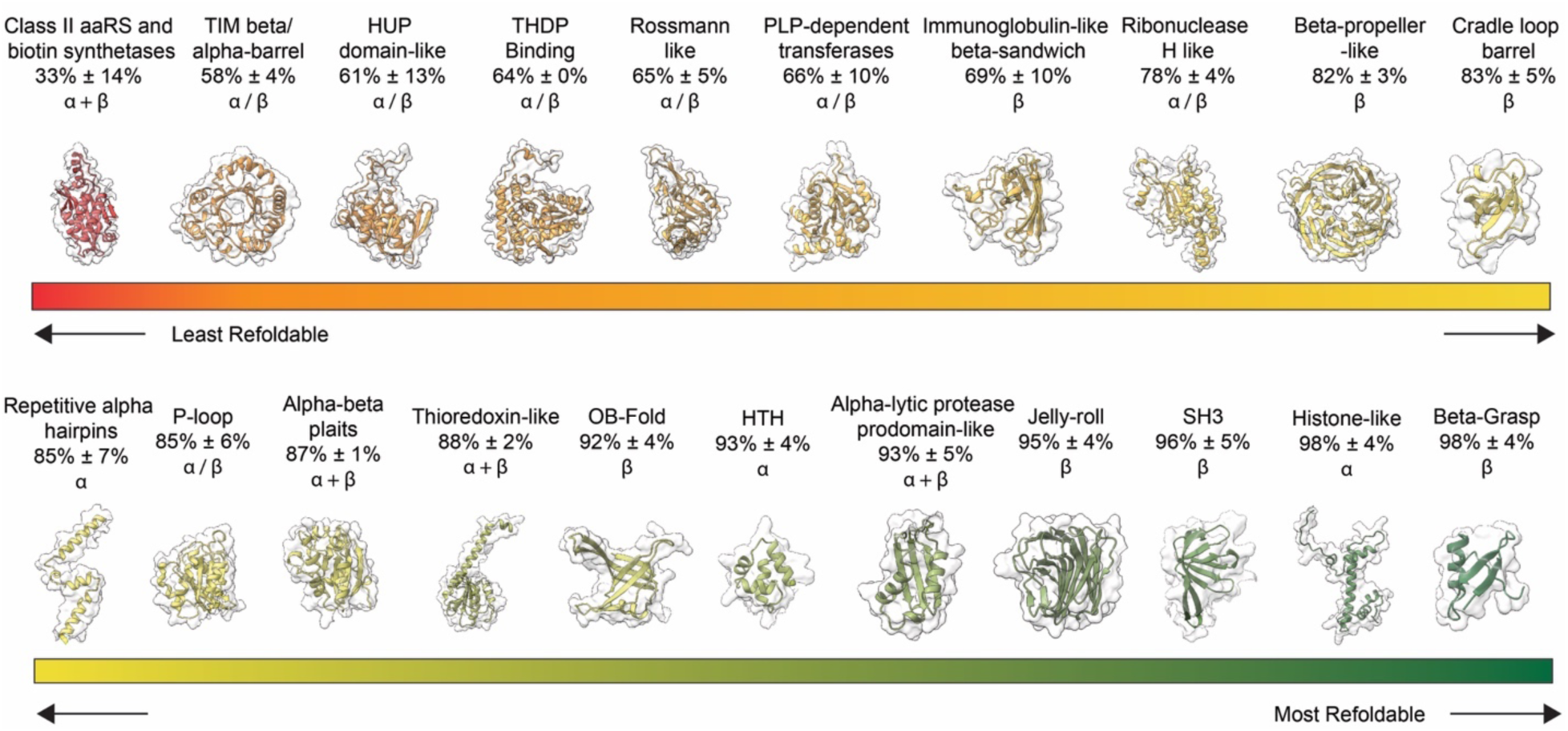
Refoldability of Different Fold Topologies. Sites were grouped to individual domains (instead of proteins) using ECOD domain definitions (ref. 26) and boundaries demarcated by DomainMapper (ref. 27). Domains were grouped by X-group (equivalent to a superfamily), and X-groups are sorted in order of least refoldable (upper left) to most refoldable (lower right). Numbers show fraction refoldability ± std. dev. from three separate performances of the global refolding assay in yeast.

**Extended Data Figure 8.**
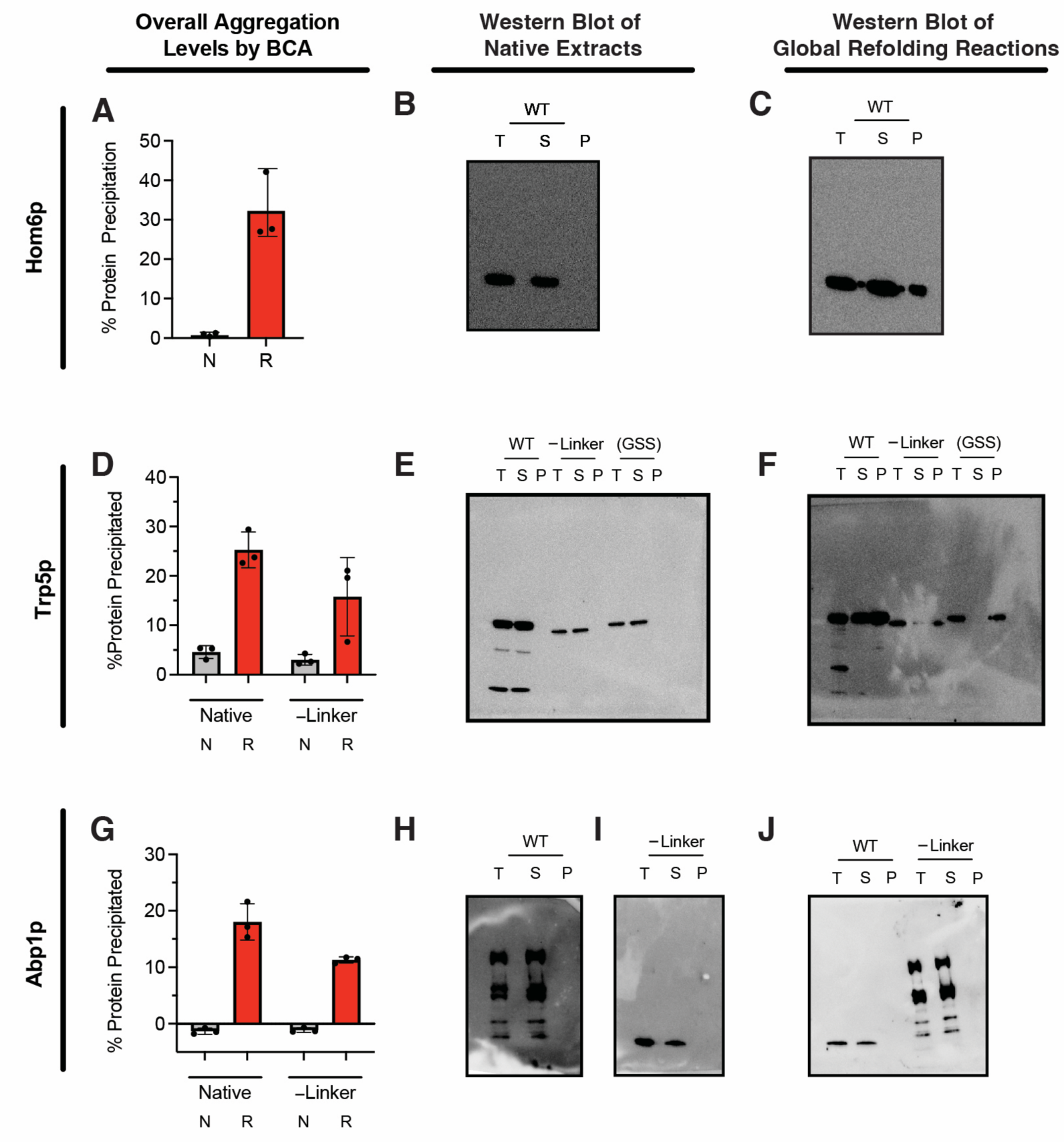
Aggregation Assays and Full Images of Western Blots. (A, D, G) Yeast strains harbouring 2µ plasmids over-expressing proteins of interest were lysed, and extracts were assessed for precipitating aggregates (by centrifugation at 16,000 *g* for 15 min) via the bicinchoninic acid (BCA) assay, either in their native forms (N) or following a cycle of global unfolding/refolding (R). Assay was performed on strains over-expressing (A) Hom6, (D) Trp5, or (G) Abp1. (B, E, H, I) Full Western blot images (probed with α-His6) of native extracts, either in total (T), or taken from the supernatant (S) or a resuspended pellet (P) following centrifugation at 16,000 g for 15 min. (C, F, J) Full Western blot images (probed with α-His6) of refolded extracts, either in total (T), or taken from the supernatant (S) or a resuspended pellet (P) following centrifugation at 16,000 g for 15 min.

**Extended Data Figure 9.**
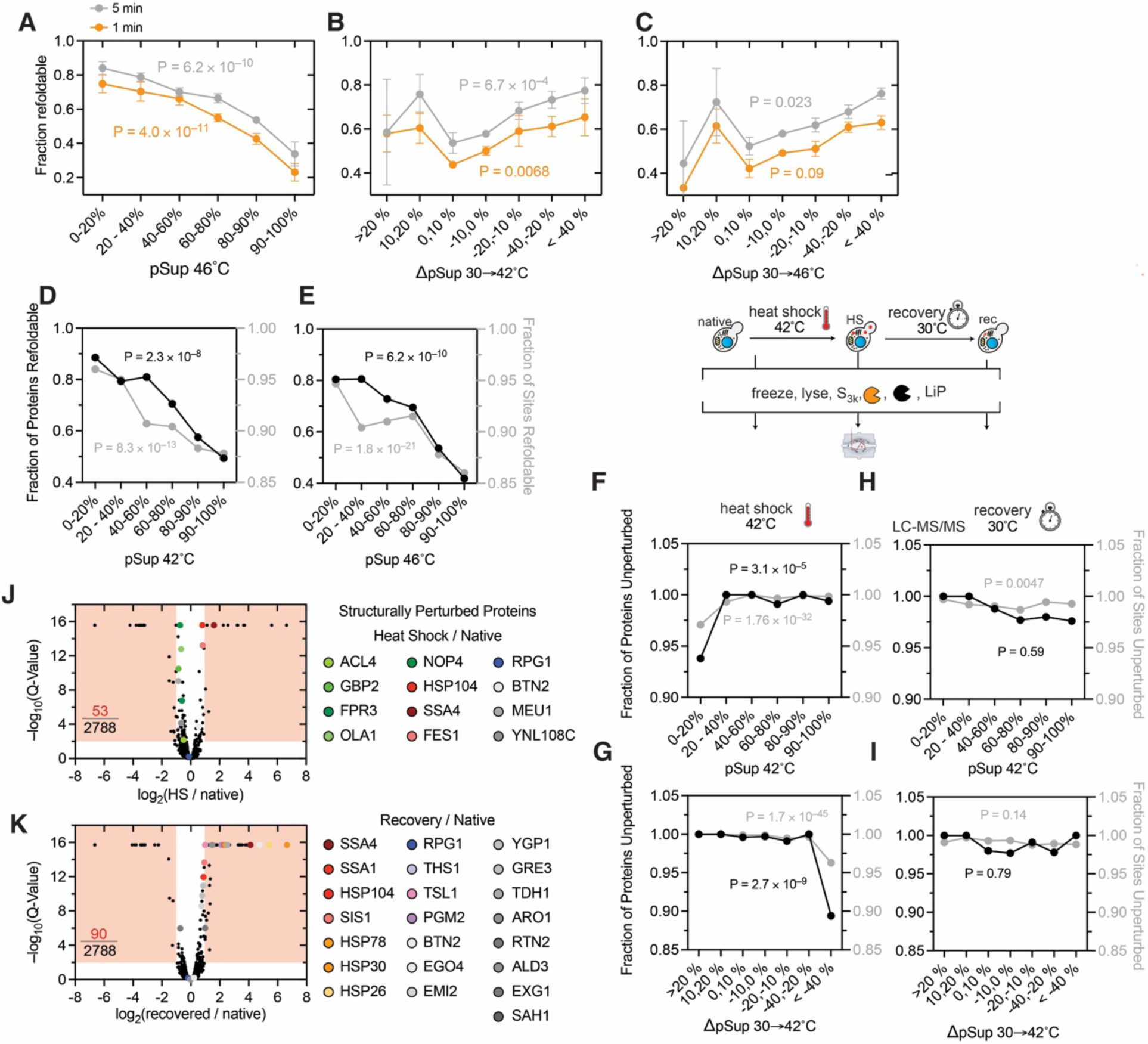
Supporting Analyses Concerning the Linkage between Condensability upon Heat Shock, Refoldability, and Structural Change upon Heat Shock. (A) Fraction of yeast proteins that refold after either 1 min (orange) or 5 min (gray) as a function of pSup46°C (proportion in the supernatant after heat shock at 46 °C), a measure of the protein’s tendency to partition to heat stress granules. P-values are from chi-square test from replicate 1 of the experiment. (B, C) Similar to panel A except using ΔpSup metrics for the same set of yeast proteins for which pSup metrics were available from ref. 40. ΔpSup(30→42°C) = pSup42°C – pSup30°C. ΔpSup(30→46°C) = pSup46°C – pSup30°C. For the most part, proteins with low pSup correspond to those with large negative ΔpSup, and those with high pSup correspond to those with ΔpSup ≈ 0. (D,E) Trends between refolding propensity and pSup at (D) 42°C and (E) 46°C. Black dots represent percent of proteins in that category which refold. Gray dots represent the fraction of sites associated with proteins in that category which are refoldable. P-values are from chi-square test, and come from replicate 1. (F-I) Trends between pSup (either (F,H) pSup42°C or (G,I) ΔpSup(30→42°C)) and (F,G) structural changes after a heat shock at 42°C for 8 min or (H,I) structural changes after recovery at 30°C for 60 min. Black dots represent percent of proteins in that category which are structurally unperturbed. Gray dots represent the fraction of sites associated with proteins in that category which are structurally unperturbed. P-values are from chi-square test. (J, K) Volcano plots showing proteins whose abundances are changed during (J) heat shock at 42°C for 8 min, or (K) recovery at 30°C for 60 min relatively to yeast grown at 30°C. *y*-axis provides Benjami Hochberg corrected P-values. Regions in red represent cut-offs for significance; in lower-left corner is the fraction of total proteins quantified with significant abundant changes. To the right are all proteins with significant *structural* changes (taken from Figure 4E-F), and the colour code they’re assigned.

**Extended Data Figure 10.**
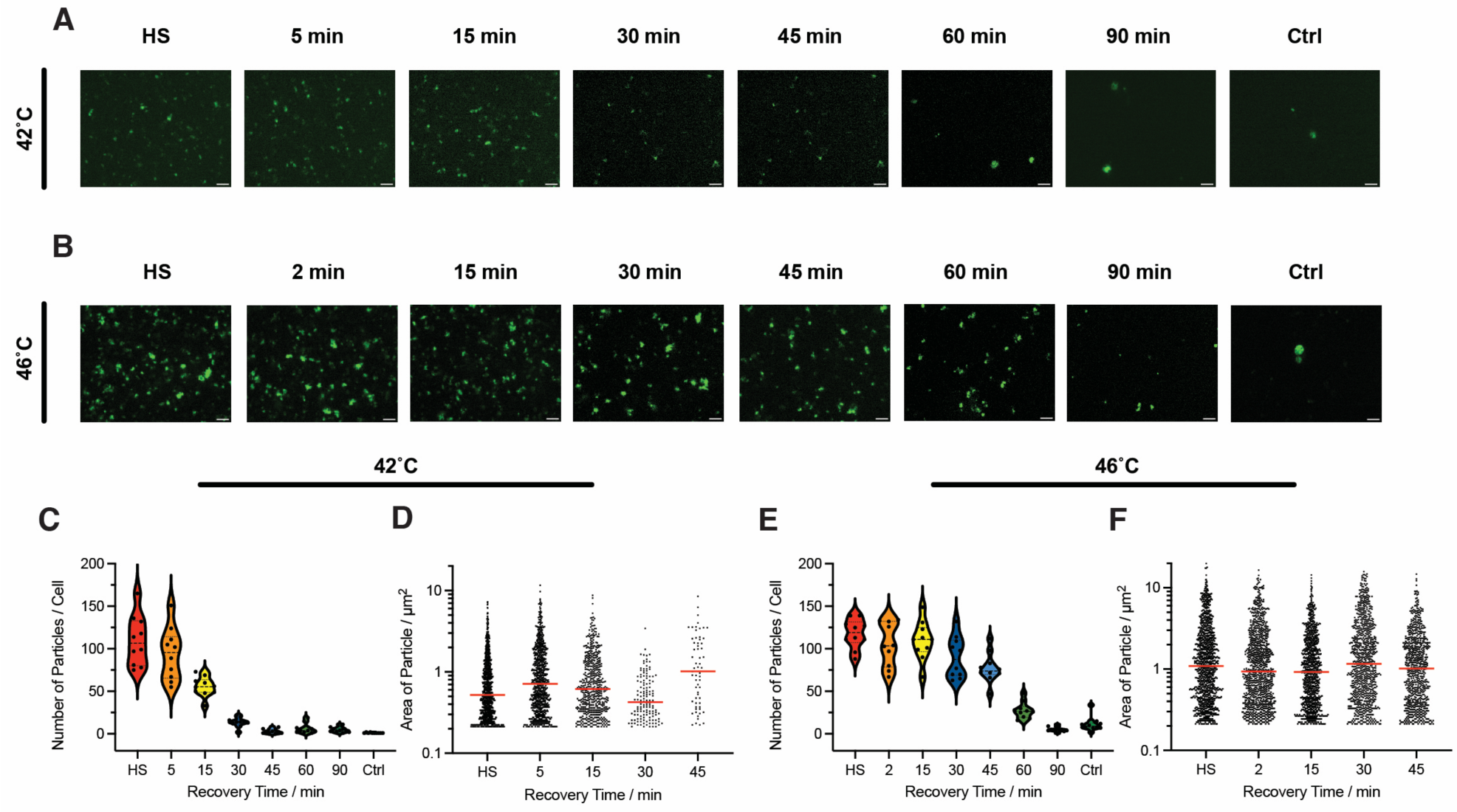
Image Analysis of Isolated Heat Stress Granules. (A,B) Representative fluorescence images of Pab1-Clover heat stress granules (HSG) formed by incubating yeast at (A) 42°C or (B) 46°C for 8 min and then recovering at 30°C for distinct periods of time thereafter. HS, immediately after heat shock; control, cells that were never subject to heat shock. Scale bar represents 5 µm. (C) Distribution of the number of HSGs per yeast cell following heat stress at 42°C and then recovering for distinct periods of time thereafter, with each datapoint representing one of ten images acquired under each condition. (D) Distribution of the area of each HSG particle following heat stress at 42°C and then recovering for distinct periods of time thereafter, with each datapoint representing one particle. (E) Similar to panel C, except following heat stress at 46°C. (F) Similar to panel D, except following heat stress at 46°C.

**Extended Data Figure 11.**
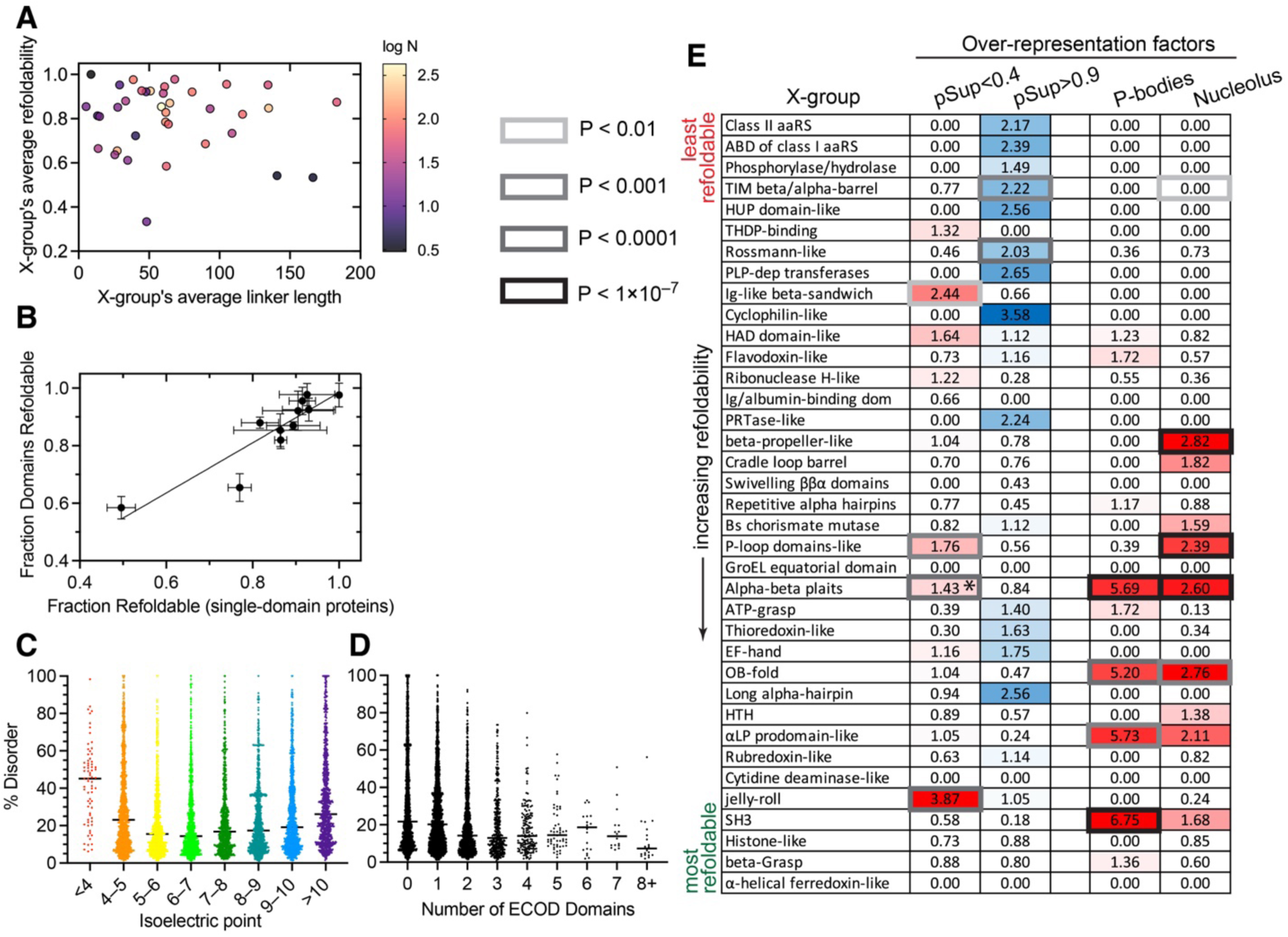
Fold Topology, Linker Length, %Disorder, Isoelectric Point, and Number of Domains Provide Independent Explanatory Information on Refoldability with Minimal Covariation. (A) Covariation analysis conducted on domains. Each domain in the yeast proteome is assigned to a fold topology (X-group) and the two linker lengths surrounding each domain is calculated. On the *x*-axis is shown the average linker length to/from domains of this X-group, and on the *y*-axis is the X-group’s average refoldability. No significant correlation is found (R^2^ = 0.01, P-value = 0.6); hence, the explanatory power of linker length is not a covariate of fold topologies. (B) Domain-wise refoldability of a given X-group is consistent with protein-wise refoldability for single-domain proteins consisting of that X-group (R^2^ = 0.76, P-value < 0.0001). (C) %Disorder for all proteins divided by their isoelectric point. The high refoldability of acidic (pI<5) and basic (pI>10) proteins could be partially explained in terms of %disorder, but there is little change in median disorder between 5<pI<10 wherein refoldability trends upward. (D) %Disorder for all proteins divided by the number of domains. The trend whereby proteins with more domains refold more poorly is not a co-variate of proteins with more domains having less disorder. (E) A table of frequent fold topologies (X-groups) in increasing order of refoldability and whether that X-group is enriched (shades of red) or depleted (shades of blue) from heat stress granules (assigned by pSup<0.4), P-bodies or the nucleolus (according to the *Saccharomyces* Genome Database under Cellular Component). Over-representation factors given as numbers in cells, and P-values according to the chi-square test that the over-representation would occur due to chance is indicated with the border shade around the cell. *was not significant using pSup<0.4 definition, but is significant using ΔpSup(30→42°C) < –0.2.

**Extended Data Figure 12.**
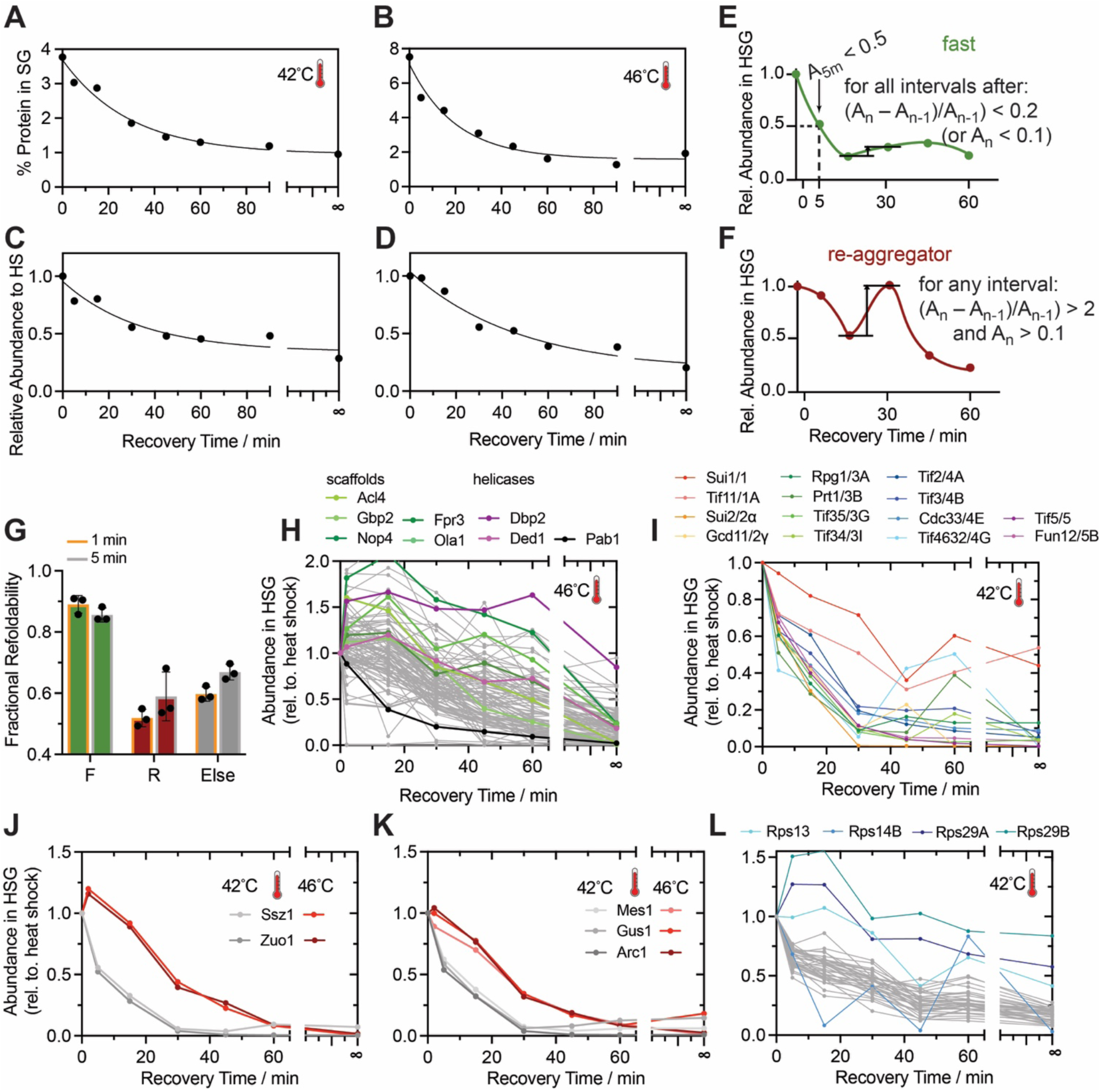
Supplemental Analyses Related to Heat Stress Granule Dispersal Assays. (A,B) The percentage of total protein content in yeast cells contained in the heat stress granule fraction, as a function of recovery time for yeast cultures subjected to heat stress at (A) 42°C, or (B) 46°C for 15 min, as determined by the bicinchoninic acid assay. (C, D) The relative amount of protein in the heat stress granule fraction, as a function of recovery time for yeast cultured subjected to heat stress at (C) 42°C, or (D) 46°C for 15 min, compared to the sample collected immediately after heat shock. (E) Definition of a “rapidly retrieved” protein. The relative abundance must be below 50% at the 5 min timepoint, and the abundance must not increase thereafter allowing for upto 20% change due to noise. (F) Definition of a “re-aggregating” protein. The relative abundance must increase by 200% between two successive timepoints and not be less than 10% abundance relative to heat shock. (G) Fractional refoldability for each dispersal category after both 1 min and 5 min refolding times. (H) Dispersal curves for all HSG proteins (pSup < 0.4) during recovery (30°C) from heat stress at 46°C. Scaffold proteins highlighted in shades of green; RNA helicases in shades of violet; Pab1p in black. Curves recapitulate behaviour of the 42°C heat stress, except abundances decay slower. “Infinite” time refers to control sample not subject to heat shock. (I) Dispersal curves for all translation initiation factors (yeast gene names and eIF names provided) during recovery (30°C) following heat stress at 42°C. (J) Dispersal curves for the two proteins of the ribosome-associated complex (RAC) during recovery following heat stress at 42°C and 46°C. (J) Dispersal curves for the three proteins of the multi-synthetase complex during recovery following heat stress at 42°C and 46°C. (K) Dispersal curves for the proteins of the small ribosomal subunit during recovery following heat stress at 42°C. All the ribosomal proteins are coherently dispersed from HSGs, except Rps13, Rps14B, Rps29A, and Rps29B.

